# Pre-Settlement Forests of Southwest Washington: Witness Statements

**DOI:** 10.1101/2021.01.04.425257

**Authors:** Tom Schroeder

## Abstract

In the mid-nineteenth century overland immigration into western Washington State passed through lands bracketed by the lower Columbia River and the Pacific Ocean. Witness trees from the region’s first GLO surveys (General Land Office), which preceded settlement, are used to reconstruct the composition, character, and distribution of the region’s natural forests. As such, this investigation augments a similar study of early forests around Puget Sound, situated immediately to the north (Schroeder, 2019). A retrospective species map is constructed from locational information from more than thirty-five thousand witness trees; accompanying tree diameters elucidate size differences by species and geographic locales. Three principal forest types were noted: western hemlock in the rainy western hills, with some Sitka spruce near the coast; Douglas-fir with woodland tree species in the rain-shadowed central plains; and hemlock/Douglas-fir/redcedar mixtures on the lower flanks of the Cascade Range. Although the majority of trees were small or medium in size, a significant fraction was large. All forest types displayed significant amounts of old growth, as judged by screening witness trees against a quantitative model. Mensuration exercises estimate that the region’s pre-settlement tree population approached one-half billion specimens with a timber volume of nearly 50 billion cubic feet.

## Introduction

The southwestern part of Washington State (Figure 1) is a relatively unfamiliar region framed by the Columbia River, Pacific Ocean, Puget Sound, the Olympic Range, and the Mt. Saint Helens stretch of the Cascade Range. This study reconstructs the region’s 4,400 sq miles of natural forests as they were at the dawn of Euro-American settlement in the mid-19^th^ century. The findings complement a similar investigation that focused on 6,300 sq miles of early forests around Puget Sound (Schroeder, 2019). Together, the two studies cover all non-mountainous parts of western Washington, except for coastal strips of the Olympic Peninsula.

**Figure 1.**
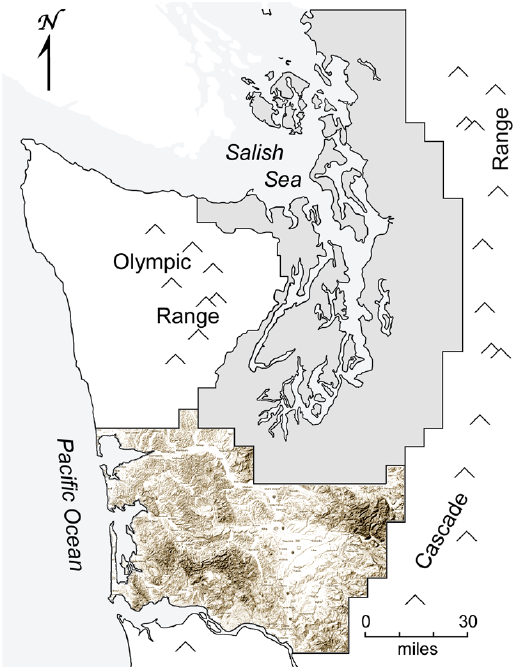
Area map of western Washington State. The study area (sepia terrain) adjoins Puget Sound country (grey), whose early forests were previously studied.

The work employs witness tree information from the region’s first land surveys conducted by the U. S. General Land Office (GLO, forerunner of the Bureau of Land Management). Beginning in 1853 and progressing for four decades, the surveys were executed in 36-sq-mile township units in a rectilinear grid across the landscape, each with data and observations recorded in field notes. Primary tree data for this study were extracted from 10,500 pages of handwritten survey field notes, all archived and available online (BLM online, n.d.).

As an adjunct to the main text of this report, a short Appendix contains important geophysical information about the region’s features. It provides much of the abiotic environmental context for understanding the forests.

## Region-wide Results

### Inventory of Species

It is assumed that witness trees accurately mirror the character of SW Washington’s pre-settlement forests. Table 1 displays basic information about the region’s 35,431 witness trees from a master database aggregated from 138 township surveys. It introduces species common names (as recorded in the surveys), numerical tallies, proportional representation (relative frequencies, i. e., tallies divided by 354.31), and average trunk diameters (linear means, in inches).

**Table 1.**
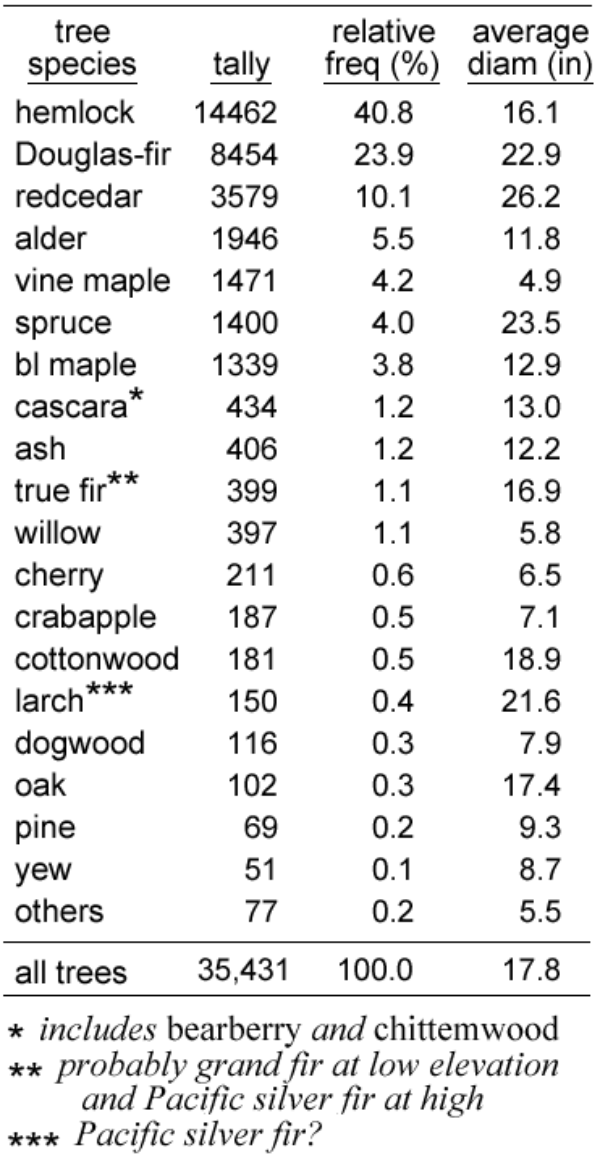
Inventory of witness trees.

Twenty-eight species of witness trees were recorded for SW Washington: eight conifers and 20 hardwoods. About 80% of stems were conifers, mostly western hemlock, Douglas-fir, western redcedar, and Sitka spruce. Of the 20% of stems that were hardwoods, the most common were red alder, vine maple, and bigleaf maple. These seven named species comprise over 92% of all witness trees.

Most common names for trees are familiar and unambiguous, but some are problematic: “pine” could mean either shore/lodgepole pine or western white pine; “true fir” could mean either grand fir or Pacific silver fir; and both “bearberry” and “chittemwood” (each with spelling variations) refer to cascara. The presence of western larch west of the Cascade crest is untenable, so “larch” in the records probably refers to one of the true firs, at high elevation undoubtedly silver fir (Van Pelt, 2007, p. 11, Fig. 3).

Diameters among witness trees ranged from 1 to 148 inches (12+ feet!), with an overall average diameter of 17.8 inches (1.5 feet). On average, conifer diameters were more than twice those of hardwoods (19.5 versus 9.0 inches), a differential that translates exponentially as an average cross-sectional area (basal area) for conifers being five times greater than that of hardwoods.

### Witness Tree Map

Figure 2 is a rectilinear, thematic species map of SW Washington’s witness trees that graphically models spatial distributions of tree species of the early forests. Each so-called corner and quarter-corner tree is represented by a small square, color-coded by species and positioned according to the GLO record: the map’s inset illustrates the eight witness trees of a survey section arranged in a “hollow square” around the section’s number – four corner trees and four quarter-corner trees at the section’s corners and sides. Thirty-six sections, numbered in a boustrophedon, compose each of the region’s townships. As elaborated more fully elsewhere (Schroeder, 2019), the map was constructed by tiling townships into numbered rows (<u>T</u>ownships) and columns (<u>R</u>anges); accordingly, T/R axes provide a convenient shorthand for locating details in the map, e. g., Grays Harbor is centered on T17/R10W. Boundaries and river courses were fitted secondarily. Being strictly rectilinear, the completed map’s format includes no meridional convergence, but it otherwise closely resembles a Lambert Conformal Conic projection, as in WDNR (2010), which includes township boundaries.

**Figure 2.**
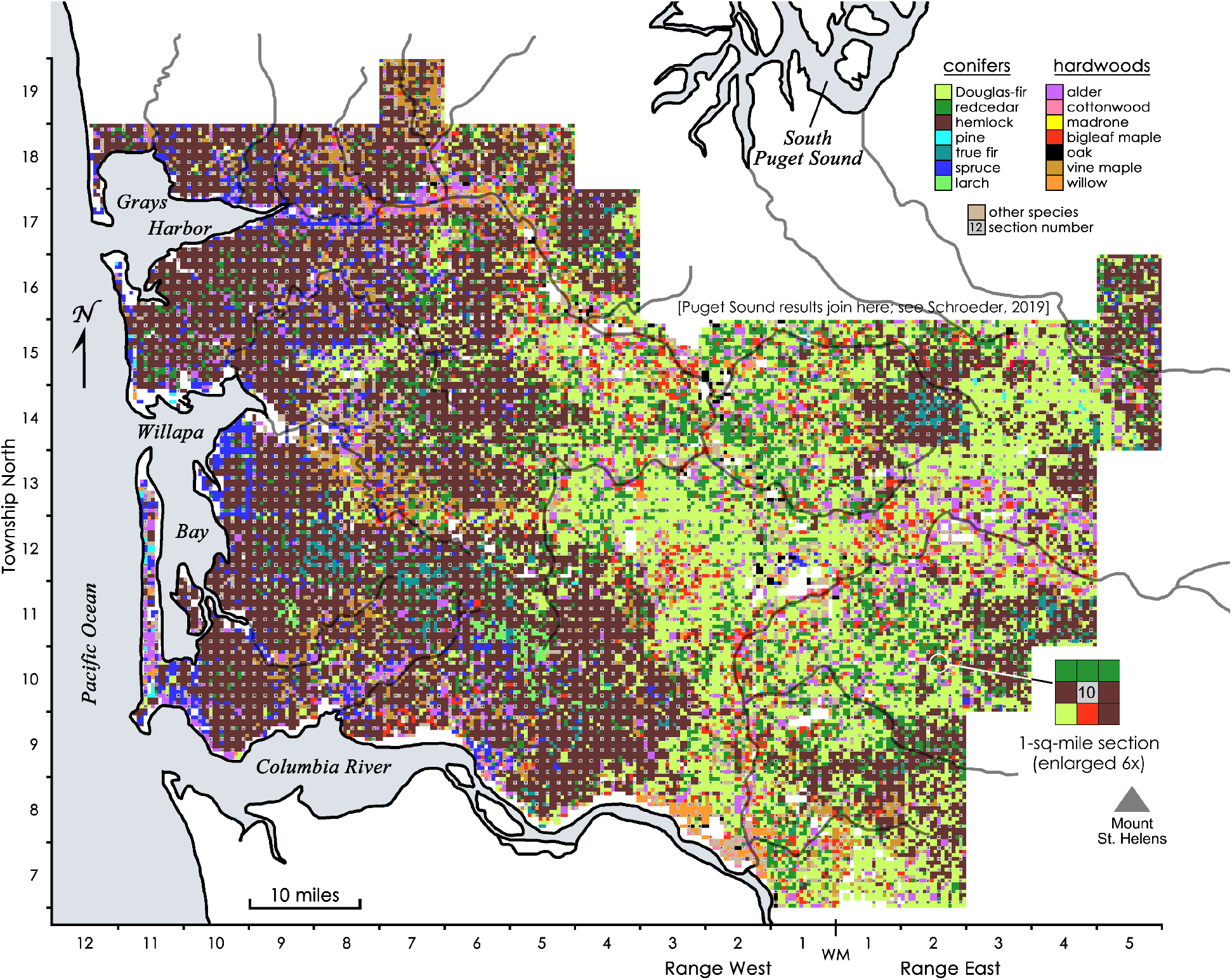
Color-coded species map of all witness trees in SW Washington; the image can be enlarged for details. Major rivers are named in the Appendix. Inset (right), a map module, section 10 of T10/R2E, enlarged to display its eight witness trees. WM, Willamette Meridian.

## Forest Types & Analysis

### Type Discrimination

Species composition across SW Washington was not homogeneous, as the species map clearly shows, but discriminating forest types is a matter of perspective. Micro-focusing on local details (Figure 3) reveals a very high degree of compositional diversity from near-monoculture to a jumble of up to ten species; while interesting, such variety constitutes more of a hodgepodge than a coherent pattern of forest types. More broadly, Figure 2 suggests that the landscape can be divided into three major forest types in three geographically discrete zones (Figure 4): western (mostly hemlock), central (mostly Douglas-fir with some redcedar and bigleaf maple), and eastern (mixed conifers); for completeness, a fourth minor zone must be added to account for a lacework of leftover forested slivers that course through and around the main zones. These four zones are delineated, nicknamed and analyzed in Figure 5.

**Figure 3.**
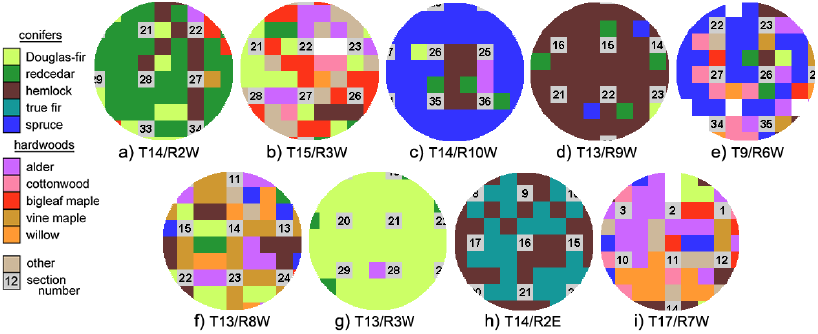
Extreme compositional diversity at small spatial scales. Each 6-sq-mile circle contains ~50 witness trees. Locations by T/R coordinates.

**Figure 4.**
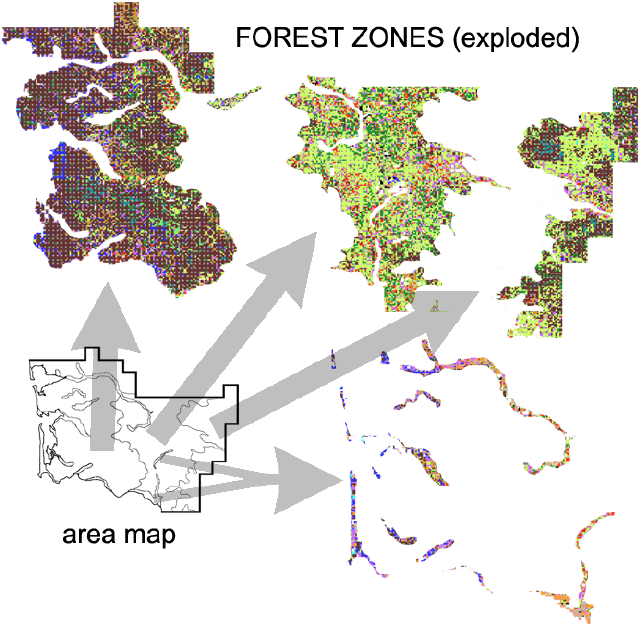
SW Washington’s forests dissected into four distinct types.

**Figure 5.**
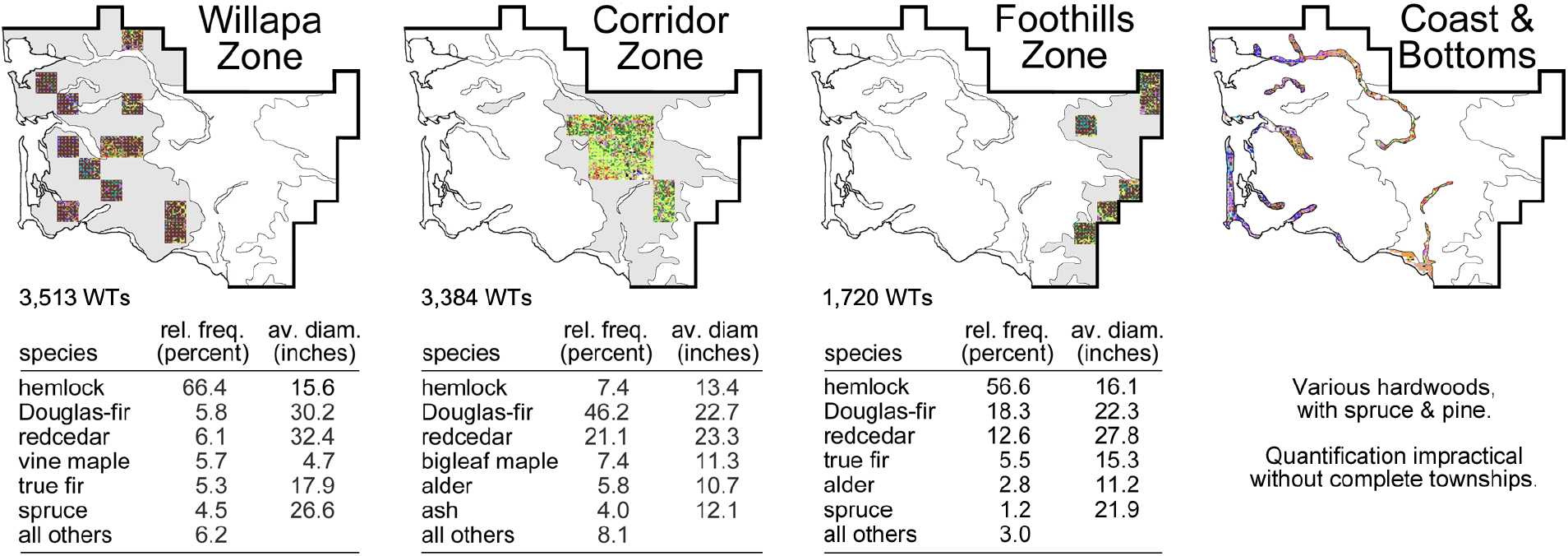
Forest zones (outlined and shaded grey) and quantitative tabulations of witness trees (WTs) pooled from representative townships (species-colored squares); rel. freq., relative frequency within the data pool; av. diam., average diameter.

Quantifying a zone’s forest type by including all of its constituent witness trees might seem an appropriate strategy, except for two drawbacks: 1) witness tree data in this study are unitized by township, whose size and squareness do not suit a zone’s irregular shapes; and 2) zone boundaries are semi-arbitrary, so gradations in species makeup near a zone’s margins dilute the more characteristic “core” composition. Accordingly, tabulated quantifications in Figure 5 are based on trees from a selected few core townships: twelve townships for each of the larger zones (Willapa and Corridor) and six for Foothills Zone. Witness tree data pooled from each subset of townships were sorted by species and analyzed; pool sizes are indicated as tallies beside each table. Coast & Bottoms components were too narrow to permit quantification by township, but their tree species may be recognized by coded colors.

Shapes and locations of the three major zones strongly correlate with visible subdivisions of geo-climatic features identified and described in the Appendix, especially topography, precipitation, and soil moisture regime. For example, Willapa Zone coincides closely with areas of high precipitation, elevations above 1000 feet, and soils that rarely dry out. Even more fundamentally, the major zones also follow the region’s three ancient tectonic realms, also described in Appendix: the uplifted hills of the Coast Range (Willapa), the planar subduction trough (Corridor), and the lower ramparts of Cascade Range volcanos (Foothills). These observations imply that the pre-settlement forest types were constrained in development by their abiotic environments. The terrains that immediately underlaid each forest type were also highly distinctive (Figure 6).

**Figure 6.**
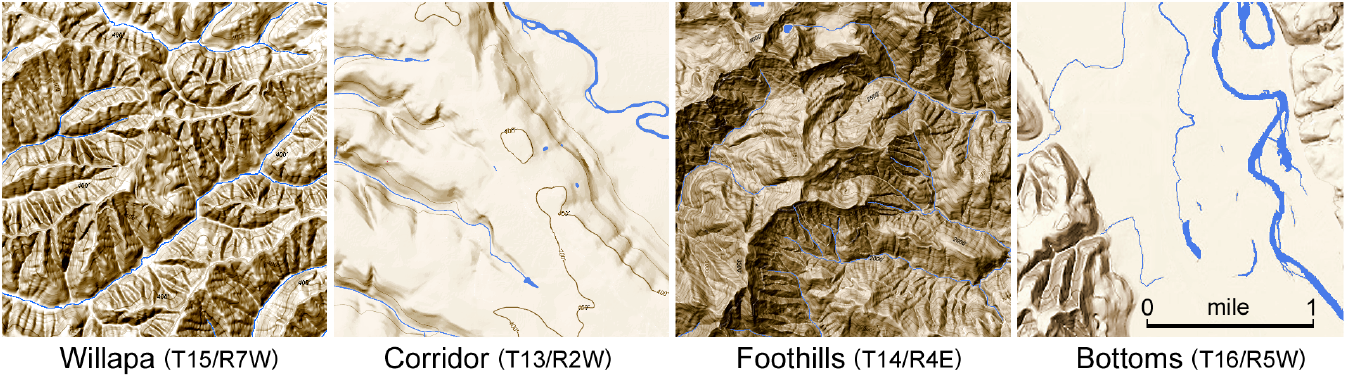
Contrasting zone terrains. Willapa’s fractal complex of ridges and incised gullies; Corridor’s low plains with shallow channels; Foothills’ mountain flanks; and Bottoms ‘floodplains (here the broad Chehalis Gap). (from Google Maps)

Forest zones will be further discussed in the chronological sequence of their surveys, namely: 1850s for Coast & Bottoms; 1860s for Corridor Zone; 1880s for Willapa Zone; and 1890s for Foothills Zone. That timeline reflects the progression of Euro-American immigration into SW Washington, including periodic slowdowns during the Panic of 1873 and the monumental Depression of 1893. The earliest land claims were generally by newcomers interested in homesteading and opening small businesses; they preferred lands that were level, accessible, and suitable for habitation, agriculture, and transportation. Later arrivals increasingly included moneyed speculators in search of exploitable natural resources rather than onsite habitation. Investors recognized that potential wealth lay in the inhospitable, but well timbered, hinterlands, and their eventual success would rely on a financial revolution, new land laws, and advances in industrial technology.

### Coast & Bottoms

These long strips of forested land were generally 0.5 to 2 miles wide, too narrow to be represented by township-wide databases, yet their land area amounts to about 180 sq miles. They included low-lying tidal plains and riparian benches, both of which featured fine-grained soils built from silts; the former experienced marine influences (including storm surges), and the latter suffered seasonal river dynamics (including floods). Hardwood tree species dominated both situations, but on sandy sites, such as the 25-mile-long spit between Willapa Bay and the ocean (T10-13/R11W), red alder shared space with hardy conifers, such as spruce and pine (probably shore pine). The soggy flood-prone bench along the Columbia, west of the mouth of the Cowlitz River (T7&8/R2W), was strongly dominated by willow. Stream bottoms were treed with red alder and other fast-growing, shade-intolerant pioneers that prospered in moist soils, such as willow, cottonwood, and Oregon ash; most of those trees remained small in diameter, likely because of periodic flooding. Alongside a tributary of the Chehalis River a few miles upstream from Grays Harbor a GLO surveyor observed:

> “Different kinds of maple, with ash, cottonwood, crab-apple, chitwood, alder & spruce are common to all the bottom lands” (GLO, T18/R8W, p. 608, 1875).

Rich soils in the generously wide Chehalis Gap were particularly attractive to early immigrant farmers, despite the risk of periodic flooding, as comparable sites had been in the Floodplains Zone of Puget Sound country (Schroeder, 2019). Riverside farms along the narrower benches of the Cowlitz River were even less secure, being more subject to seasonal melt waters and more regular flooding. Tidal sites close to the Pacific shore were not suitable for ordinary farming, but fishing, clamming, and shipping allowed small seaside communities to develop nonetheless.

### Corridor Zone

Long before Europeans arrived, the center of SW Washington presented the favored inland route for travelers and traders between the lower Columbia River and the southern inlets of Puget Sound, hence the zone’s nickname Corridor. Beginning in the 1830s, employees of the Hudson’s Bay Company, headquartered at Fort Vancouver (T2/R1E), also used the Cowlitz River and parallel trails to access their agricultural operations at Cowlitz Farm, situated on the prairie near the modern town of Toledo (T11/R1W), and Fort Nisqually near Dupont (T19/R2E). Many early settlers followed the route in search of homesteads in SW Washington or to continue on to Puget Sound. Travel on or along the river was not altogether trivial in the early days, but at least the terrain was relatively flat:

> “The route traveled from the Columbia River (to Puget Sound) is by canoes, for twenty-eight miles, up the Cowlitz to the settlement at ‘Cowlitz Landing,’ (or by horse over a somewhat bad path) and then by horses or mules to Olympia, fifty-two miles, over a tolerably level country, and by a road moderately good in summer, but bad in winter…. The Cowlitz has a rapid current, and at a low stage of water, canoes are poled up its channel; during freshets they are dragged up, the crews cling to the branches of the trees upon its banks.” (Davidson, 1869).

Cowlitz Landing, consisting of a store and two rustic hotels for travelers, acquired a place in history in 1851 when a handful of pioneers started a petition to separate Washington Territory from Oregon Territory, which became a reality two years later. The route beside the Cowlitz River soon thereafter supported a military road, telegraph lines, and mail service; in the 1870s a rail line connected Olympia and Longview at the mouth of the Cowlitz, a development that enhanced timber extraction. The same transportation axis today supports Interstate 5 highway, Amtrak rail, and Bonneville power lines.

The 1,500 sq miles of Corridor Zone forest traversed by early settlers generally resembled those previously described for the southern parts of Puget Sound country (Schroeder, 2019), all of which lie in a rain shadow and experience 45-55 inches of rainfall annually. As quantified in Figure 5, nearly half of Corridor stems were Douglas-fir and another one-fifth were redcedar, both with tree diameters that averaged nearly 2 feet. Though widely distributed overall, redcedar were noticeably more concentrated in a few places (e. g., T14/R2W and T11/R1&2E), whereas other places were so devoid of redcedar that they resembled Douglas-fir monocultures (T12&13/R3W, near the modern town of Vader) with diameters that impressively averaged more than 28 inches. Bigleaf maple were also well represented in a few locales, e. g., upper and lower Cowlitz River (T12/R3W) and along the Chehalis between the Newaukum and Black Rivers (T15/R3W). Hemlock, however, were strikingly absent from Corridor, except where vegetation graded into neighboring forest zones.

The principal species suggest that conditions in the zone were somewhat droughty. Indeed, about twenty small patches across Corridor Zone lacked witness trees altogether and appear as uncolored blanks in Figure 2. According to surveyors’ notes, most blank patches were identified as grassy prairies, except for a few smaller flood sites or marshes. The largest prairie covered at least 10 sq miles and was occupied by the Company’s Cowlitz Farm for raising livestock and wheat; soils at the location display an unusual aquic moisture regime (see Appendix).

Garry oak specimens were distributed in and around the putative prairies. Figure 7 identifies twenty-three townships whose witness trees included at least one oak; the six townships with the highest abundance of oak witness trees (representing the top quartile of townships) are highlighted. In SW Washington, oak trees occurred virtually nowhere outside of the Corridor Zone and were less abundant than around Puget Sound.

**Figure 7.**
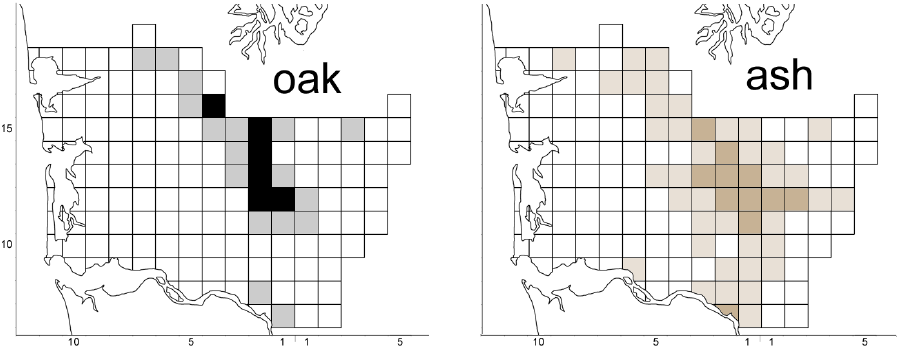
Distributions of Garry oak and Oregon ash coincided in Corridor Zone. Tints: nominal presence of species in township; full colors: the quartile of townships with greatest abundance; uncolored: species absent.

Another Corridor-specific tree was Oregon ash (Figure 7). Although ash is more partial to moist sites than oak, the two sun-loving hardwoods shared loci of occurrence and concentration. Their co-distribution in parts of SW Washington, apparently without an obvious understory of shade-tolerant species, may indicate that forest density in much of Corridor Zone was lower than elsewhere, perhaps more like semi-sunny parklands than closed-canopy forest; such landscapes, including outright prairies, would have been particularly attractive to early homesteaders keen on raising livestock.

As elaborated in the Appendix, Corridor Zone is geologically distinct from other parts of SW Washington. The land is a triangular plain composed of volcanic breccia and finer debris emanating from Mts. Rainier and St. Helens. It gradually inclines from a low point near the the confluence of the Chehalis and Skookumchuck Rivers in the north (T14/R2W) to a 500-foot-high ridge that separates the Cowlitz/ Toutle and Newaukum/Chehalis watersheds (T11//R1&2W).

### Willapa Zone

Both the physical character and forest cover here were most unusual. The zone’s approximately 2,100 sq miles of unrelentingly rugged and inhospitable land results from its geology and climate. Superabundant precipitation from onshore weather systems (up to 120 inches at elevations above 2000 feet, see Appendix) feeds innumerable rills and perennial streams that have eroded decomposing basalt (Willapa and Doty Hills) and bedded sediments (Black and Satsop Hills) into a fractal of steep-sided gullies, sharp-angled ridges, and hardly any level ground (Figure 6).

Throughout their arduous traverses, GLO surveyors noted that Willapa’s land was disturbingly uneven and the vegetation was intimidating, using such phrases as:

> “…unending roughness, broken & mountainous character …everywhere steep and broken …every stream large or small has cut a very sharp and steep gorge …sharp backbone ridges extending in every direction …exceptionally difficult to survey …not worth a Continental Dollar per Township …so dense that sunshine never reaches the ground …always green and wet …underbrush cannot be described” (GLO, various).

Hemlock was, by far, the most abundant tree species in Willapa Zone forests, comprising 66.4% of witness trees pooled from the twelve core townships (Figure 5) and as high as 82.9% in T9/R4W. Oddly, no single species clearly ranked second in abundance: rather, Douglas-fir, redcedar, vine maple, silver fir (putative), and spruce each contributed about 5% of the zone’s trees, on average. Furthermore, rather than being widely distributed throughout the zone, each “secondary” species had preferred locales: spruce near the seashore, vine maple in some river valleys, large-diameter Douglas-fir and redcedar in separate “islands”, and silver fir at the highest elevations (teal color at T12/R7-8W; light green at T11/R9W and T10/R5-6W). [N. B. Some uncertainty lingers about silver fir identification: surveyors applied various common names to true firs. The teal color in the species map probably refers to grand fir at low elevations, but to Pacific silver fir at high elevation, as suggested by Van Pelt (2007, p. 11, Fig. 3). As a further complication, some high-elevation trees were misidentified as “larch” (light green), which are also thought to have been silver fir.]

Spruce and Douglas-fir demonstrated a degree of mutual exclusivity (Figure 8). Although spruce occurred throughout Willapa Zone and beyond, the quartile of townships having the highest proportion was concentrated near the coast. Conversely, many of the areas rich in spruce were notably deficient in Douglas-fir; indeed, a dozen coastal townships, amounting to about 200 sq miles of Willapa forests had no Douglas-fir witness trees whatsoever. The paucity of Douglas-fir begs an explanation: leached soil nutrients, marine salts, selective windthrow, inhibited germination, unstable substrate, over-saturated conditions?

**Figure 8.**
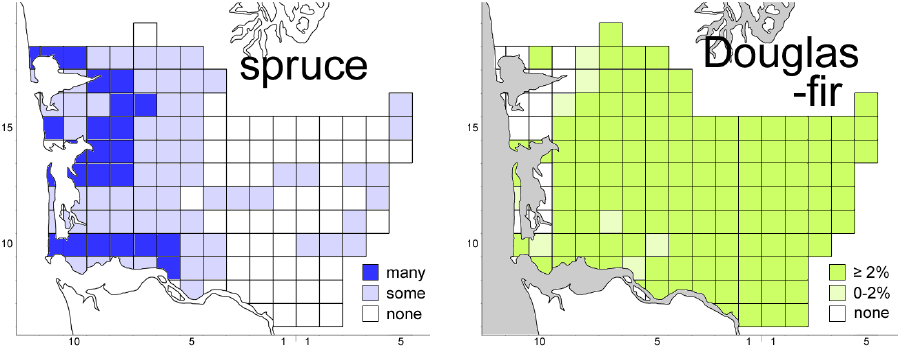
Spruce was concentrated in western townships (intense blue for quartile with highest abundance), whereas Douglas-fir in similar areas was very sparse or completely absent.

Forests overwhelmingly dominated by hemlock have been classified by forest scientists as extreme variants of the Western Hemlock type, even though the relative absence of Douglas-fir and redcedar seem contrary to conventional processes of ecological succession. A somewhat similar forest type found in the Sitka Spruce Zone is characterized by spruce dominance over hemlock; it forms a narrow belt along the state’s entire Pacific coast with fingers that reach up into Olympic Peninsula’s valleys. In that instance, Franklin and Dyrness (1973, p. 62 and Fig. 27) postulate that reproduction of faster-growing spruce is favored over hemlock when forest openings are created by windthrow, resulting from strong onshore windstorms along the coast. One might conjecture that the westernmost parts of Willapa Zone, where spruce was likewise common, represented a transition between Western Hemlock and Sitka Spruce Zones in forest character or, indeed, that the whole of Willapa Zone more properly belongs with the Sitka Spruce Zone, recognizing that spruce recruitment diminishes eastward as the strength of onshore windstorms declines. In any case, GLO survey notes occasionally refer to downed trees in the western stands of Willapa Zone, most explicitly in the following example from near the headwaters of Willapa River:

> “during the past 10 or 15 years, a terrible wind storm has passed over it, almost completely ruining the timber. What timber is still standing is so obstructed by that which is down, as to render it almost worthless” (GLO T12/R6W, p. 288, 1890).

That storm was likely the “Great Gale” of 1880 whose gusts reportedly peaked at 138 mph. Similar extratropical cyclones – known locally as “Big Blows” with winds exceeding 75 mph – strike coastal areas of Washington once every 25 to 50 years (State Climatologist, n.d.); the Columbus Day Storm of 1962 downed more timber in half a day than the state’s typical annual harvest in that decade.

### Foothills Zone

Coherent forest patterns are less clear in this narrow 600-sq-mile zone, whose integrity is intruded by low-lying valleys (T12/R3&4E), irregular projections of Douglas-fir from Corridor (T10/R2E), and high-elevation subalpine volcanic ridges; only a few townships are free from such irregularly shaped encroachments. Regardless, quantifications in Figure 5 do confirm the expectation that the zone’s forests would be a southerly extension of “Hemlock Heights” of Puget Sound country described in a previous study (Schroeder, 2019): more than half of witness trees pooled from its six “core” townships were hemlock (56.6%), and Douglas-fir and redcedar abundances were intermediate (respectively 18.3% and 12.6%, much higher than those species’ values in Willapa Zone).

The selection of townships to represent the core forest was complicated by a large anomaly in the zone’s far north end. That irregularity appears as a conspicuous “hemlock island” (centered on T14/ R2E) incongruously surrounded by a seemingly out-of-place 100-sq-mile stretch of Douglas-fir (T14/ R3E and adjacent townships), even though the opposing forest types shared the same bold tongue of terrain at 2000 to 3000 feet (Figure 9) and identical levels of precipitation (see Appendix). Hemlock dominance in the first township (brown) is no surprise, given the elevation, but the abrupt transition to Douglas-fir (chartreuse) in townships to the east is unexpected. Fortunately, GLO field notes offers an explanatory clue:

> “… covered with second growth Fir and Hemlock timber from 6 to 24 ins in diameter…. At one time some 75 to 100 years since it was covered with heavy timber, as shown by large stubs of burnt timber, which are still standing among the second growth timber” (GLO T14/R3E, p. 291, 1910).

**Figure 9.**
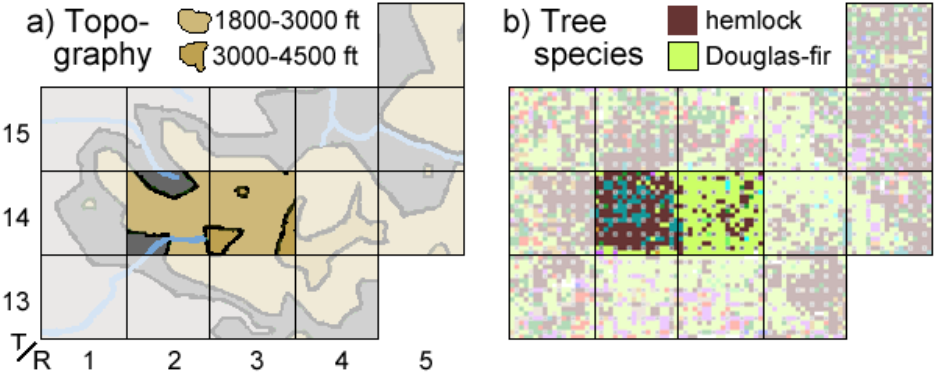
Corresponding maps of topography (from Appendix) and species (Figure 2). Hemlock dominance (T14/R2E-4E) displaced by Douglas-fir (T14/R3E) despite similar high-elevation terrain. Full colors: townships for comparison; pale colors, adjacent townships. Grey areas in (a) below 1800 feet.

Evidently, the “hemlock island” retained its late-seral hemlock tree cover by somehow escaping a large-scale wildfire that occurred around 1810-1835, unlike forest stands to the north, east, and south (presumably also covered in hemlock before the fire) which were fully burned over. Accordingly, after several decades of regrowth, when T14/R3E was surveyed in 1910, Douglas-fir (relative frequency 74.1%) and a few pine (1.4%) had = reestablished a new stand of early-successional pioneers: their diameters averaged 10.9 and 9.8 inches, respectively – modest values indicative of moderately young trees. In 1896, when the “hemlock island” T14/R3E was surveyed, its hemlock abundance was 60.3% and diameters averaged 15.8 inches – still not particularly old, despite being a slower-growing species; an additional 21.6% of trees in the township were putatively silver fir (teal color), whose average diameter was 16.7 inches.

## Exploratory Exercises

So far this report has focused on spatially segregated forest types in SW Washington that probably developed in response to geographic factors. Before concluding, two additional quantitative topics will now be examined: large-diameter trees and successional status as old growth.

### Very Large Trees

The average diameter of witness trees in SW Washington’s pre-settlement forests was 17.8 inches and the median (the population’s middle value) was 14 inches. These simplifying reductions, however, fail to reveal the full range of diameters; indeed, the size profile of all 35,341 trees in the database is shaped like a hockey stick (Figure 10a): three-fourths of all trees were 1-20 inches in diameter and form the long “shaft”. The shorter “blade” comprises 1,873 trees that ranged from 48 to 148 inches across; these specimens (5.3% of all witness trees, average diameter 57.4 inches) constitute the size-class that henceforth will be referenced as “very large trees” (≥ 48”).

**Figure 10.**
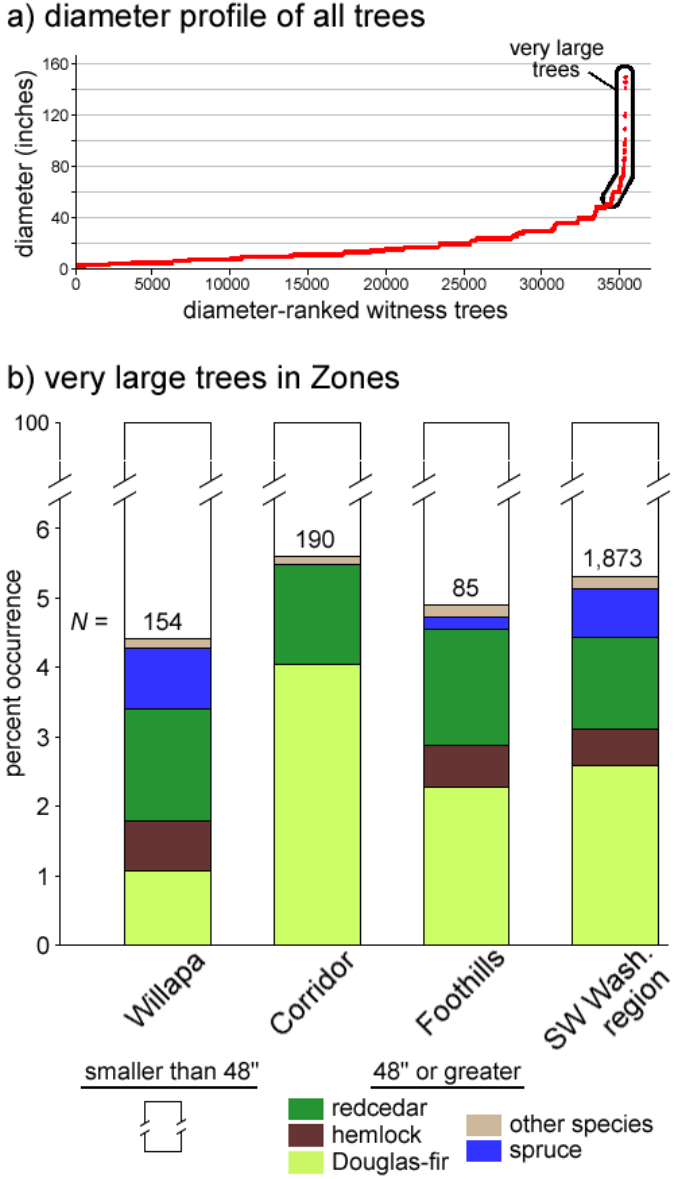
Very large witness trees: a) in the hockey-stick blade; b) by species and zone. N, tallies in databases (core townships for zones and all townships for region).

Tallies and species breakdowns of very large trees in circumscribed forest zones were determined from the separate subsets of data pooled from selected townships. Species proportions differed between zones, as shown in Figure 10b. For example, in Corridor Zone, where Douglas-fir was more strongly represented in the population, three-quarters of very large trees were of that species, and, unsurprisingly, none was spruce; by contrast, in Willapa Zone less than one-quarter of its very large trees were Douglas-fir and one-fifth were spruce. The proportion of redcedar among populations of very large trees was about one-third across the region as a whole.

At the rate of one out of nineteen (5.3%), these outstanding trees were so common that, on average, at least one very large tree should have been observable from every vantage point in SW Washington. According to their species breakdown (Figure 10b, right column), half of the time the observed tree would be a Douglas-fir. In reality, of course, very large trees were not distributed uniformly, yet every township’s database included at least one very large witness tree (Figure 11); some townships had many (e. g., T8/R2E) and others had few (T14/R4E).

**Figure 11.**
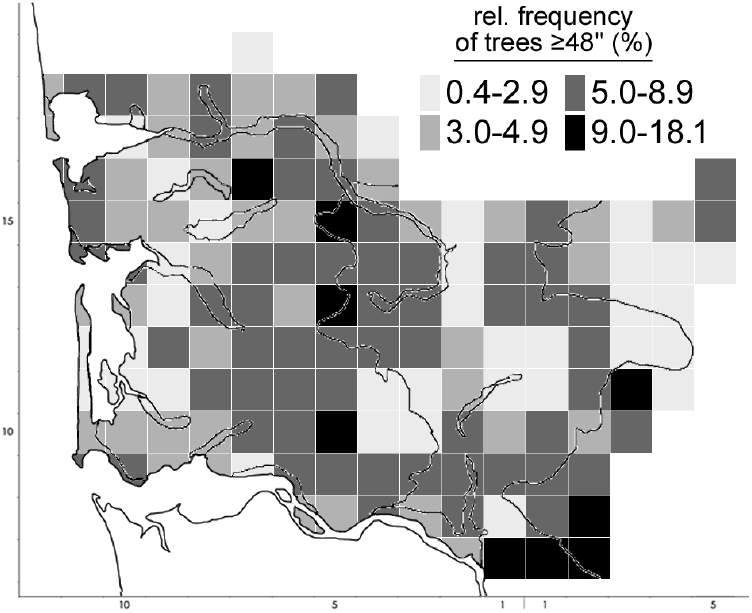
Every township had a very large tree, some had many. Zone boundaries are indicated.

To appreciate the impressive size of such trees requires some imagination: to embrace even the *smallest* member of the class requires two people with linked arms. The immense size attained by these specimens suggests that they had been growing for very long periods of time, probably hundreds of years. Being so old, their developmental relation to the generation of younger and smaller trees deserves some thought; very large trees in SW Washington presumably represented vestiges of one or more ancient generations of trees, rare individuals that had surreptitiously survived one or more stand-replacing disturbances in prehistoric times.

### Oldgrowthness

A rule-of-thumb in forest science asserts that, following a stand-replacing disturbance, a regenerating stand attains old growth status after about 200 years of orderly successional development, even if objective diagnostics of the exact transition point are elusive. According to orthodox views of ecological succession in most forests of the Western Hemlock Zone, sun-loving pioneer species like Douglas-fir first repopulate disturbance-cleared sites; only after 75-100 years of pioneer growth do shade-tolerant species such as hemlock and redcedar begin to appear (thanks to crown closure, shade, accumulated litter, and stabilized soil moisture), and thereafter they become dominant. That successional sequence suggests that at the 200-year mark pioneering Douglas-fir would be approximately one hundred years older than the oldest hemlock or redcedar specimens. Predicated on such age differentials, then, a stand’s old growth status could be objectively determined by counting annual growth rings in its trees; however, coring trees with an increment borer is too labor-intensive to be applied comprehensively across a landscape, so, for practicality, if the presence and onset of old growth needed to be ascertained in the field, a simpler age test – or a quick proxy – would be required.

In the 1980s, when conservation of old-growth was valued by society, a committee of forest stakeholders devised a field-worthy alternative to tree coring (Old-Growth Definition Task Group, 1986). The operational premise was that tree diameters can serve as species-specific proxies (stand-ins) for tree ages; accordingly, a diagnostic formula, entitled the Interim Definition, allowed forest managers to discriminate old growth stands (those to be conserved) from less developed stands (candidates to be logged) by measuring and counting the stand’s larger, dominant trees. Absent any preferable tests of oldgrowthness, the Interim Definition will be employed here as a kind of yardstick to hold against witness tree data for evidence of old growth in pre-settlement forests. Admittedly, the chosen criteria were not designed to accommodate the wide diversity of growing conditions found everywhere in SW Washington or the inflated spatial scales and vast landscapes sampled by relatively few witness trees.

For forest types in western Washington, the Interim Definition asserts that, on a per-acre basis, a stand qualifies as old growth if it contains either: a) eight or more shade-intolerant Douglas-fir trees at least 32 inches in diameter; or b) twelve or more shade-tolerant trees at least 16 inches in diameter (i. e., either hemlock or redcedar). In a spirit of exploratory enquiry, not scientific rigor, these threshold criteria will be applied as benchmarks against collections of witness trees to test whether the data sets contain the requisite numbers and sizes to qualify as old growth. As expressed in their original form, the criteria and the target data are incommensurate and thus unsuitable for comparison, like apples and oranges: the Interim Definition specifies *absolute numbers* of trees *per acre* (without specifying total tree density or population size), whereas classes of witness trees are expressed as *relative proportions* (percentages), because they are samples of a parent population whose size is also unspecified. To harmonize the disparate forms, and thus bring them into alignment, it is necessary to postulate concrete population sizes and/or to assume a common forest density (trees per acre), from which population sizes can be derived.

Although arbitrary, the operational expediency of asserting a *universal tree density* solves the incommensurability problem between Interim Definition criteria and class sizes of witness trees. Arguably, densities of most real forest stands fall within a range of 75 to 400 trees per acre (excepting young thickets, which are denser); therefore, for the purposes of the present exploration, a plausible blanket density of 150 trees per acre will be assumed for all stands (those addressed by the Interim Definition as well as pre-settlement stands sampled by witness trees). [N. B. At the chosen density, the average distance between trees would be 18 feet.]

According to the above postulated density, Interim Definition’s criteria for old growth translate as:

- at least 5.3% of Douglas-fir (eight per 150 trees) must be ≥ 32 inches in diameter, or
- at least 8.0% of hemlock or redcedar (twelve per 150 trees) must be ≥ 16 inches in diameter.

In their new, proportional forms (percentage values above), the benchmarks are now compatible with proportional classes of witness trees and therefore can be used screen databases for evidence of old growth, as the following three tests will illustrate.

TEST I – REGION-WIDE: To test for old growth in SW Washington’s pre-settlement forests *as a whole*, all witness trees in the database (35,431 in total) were filtered by species and diameter. Subsets of Douglas-fir ≥ 32 inches in diameter and of hemlock/redcedar ≥ 16 inches in diameter were then isolated and their members were counted to determine whether they were sufficient in number to meet or exceed the requisite Interim Definition percentages. Accordingly, the results of this test follow:

- 1,972 Douglas-fir (5.6%) were ≥ 32 inches in diameter (i. e., 1972 ÷ 354.31), and
- 6,117 hemlock & 2,345 redcedar (17.3% & 6.6%) were ≥ 16 inches in diameter (23.9% combined).

All the above percentages (excepting redcedar’s) exceed the required criteria, thus old growth was demonstrably abundant, and probably widespread, throughout the region’s forests. Indeed, the test confirms the presence of old growth *separately* by shade-intolerant Douglas-fir (though only barely) and the combination of shade tolerants (decisively by hemlock alone, though not quite by redcedar alone).

DISCLAIMER! Upon reflection, the foregoing exercise contains a potential flaw that may weaken or invalidate the conclusions: a previous discussion of so-called very large trees (≥ 48”) raised the possibility that such large-diameter specimens may have persisted into pre-settlement times as surviving vestiges of ancient populations. If so, it may be inappropriate to include such venerable “ringers” in the screening for old growth (imagine the unfair consequences if a professional wrestler was allowed to join a team of amateurs). Upon the assumption that the genuine objective of an old growth investigation is to evaluate only the principal generation of trees – those that developed since the most recent major disturbance – then inclusion of the cohort of legacy trees from earlier generations would necessarily skew the results toward a false indication of old growth that the principal population would not merit on its own.

#### TEST II – REGION-WIDE (REVISED)

To guard against a possible bias posed by large legacy trees, the exploratory exercise was rerun after discounting the size-effect of very large trees (only their size, not their presence, so the population basis of 35,431 was unaltered). Thus, in this revised test, the database was filtered to include Douglas-fir only between 32 and 47 inches in diameter and hemlock/redcedar only between 16 and 47 inches in diameter. Results of the revised are:

- 1,068 Douglas-fir (3.0%), and
- 5,929 hemlock & 1,864 redcedar, (16.7% & 5.3%, or 22.0% combined).

These results again clearly indicate that, even after very large trees are discounted, old growth was widespread, but only by the contribution of hemlock (alone or in combination with redcedar). By themselves, neither Douglas-fir nor redcedar had enough specimens of the required sizes to indicate old growth.

#### TEST III – ZONE-SPECIFIC

The presence of old growth in early forests of SW Washington as a region does not preclude the possibility that old growth occurred only in some zones, but not in all. The present tests explore old growth in each of the major forest zones delineated in Figures 6 and 7. As will be recalled, smaller witness tree databases for zones comprise only data pooled from representative townships, but the same Interim Definition criteria are employed and the role of very large trees was again discounted. Test results for zones, and interpretations (grey panel), are as follows:

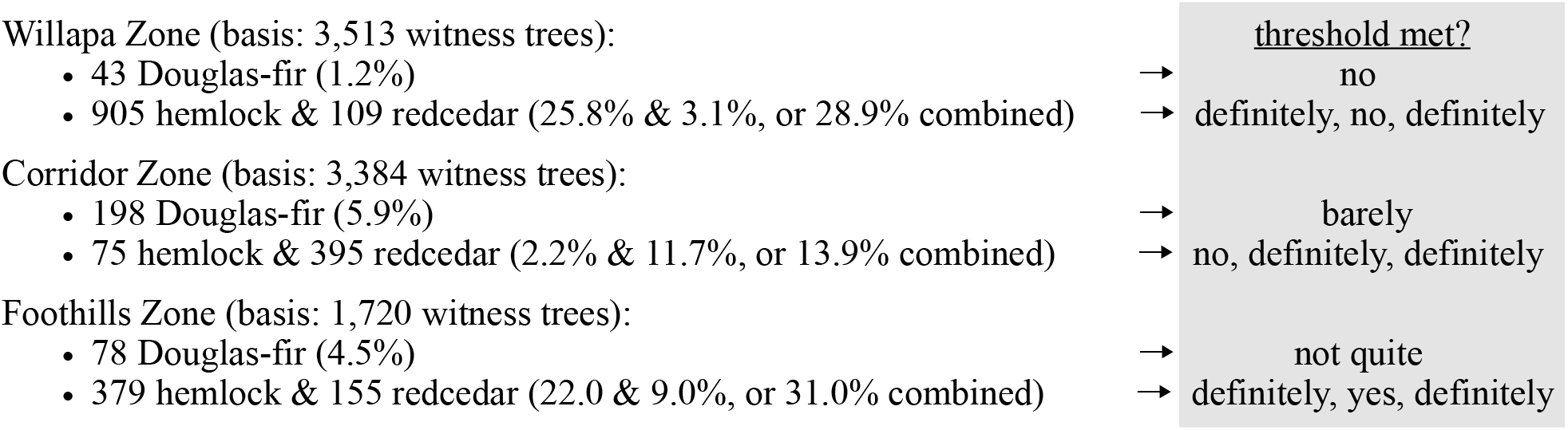

In keeping with previous test results, and contingent upon the initial assumptions, old growth was evidenced in the each of the three zones, but not by all species in each zone. In Willapa, only “hemlock old growth” occurred. In Corridor, “redcedar old growth” was fairly common, presumably confined to places where redcedar was abundant (Figure 3); some “Douglas-fir old growth” also occurred, but “hemlock old growth” was absent. In Foothills, “hemlock-redcedar old growth” was abundant, and some “Douglas-fir old growth” likely also occurred.

## Epilog

Big-picture quantification completes this examination of the early natural forests of SW Washington, but that topic inevitably segues to the post-settlement emphasis on *timber*, that is trees destined for the sawmill in the service of society: forest trees as a commercial commodity. Thus, this report ends with a brief history of how post-settlement societal events altered the pre-settlement landscape.

### Tree Tally

In a previous section that considered oldgrowthness, an average forest density of 150 trees per acre was postulated. That density value also allows the total number of trees across SW Washington to be estimated by simply multiplying tree density and the region’s land area in acres (150 × 640 acres per sq mile × 4,400 sq miles), which equals 422,400,000 trees. Four hundred million trees is an impressively large number, especially when one considers that they would be cut before the appearance of the portable gasoline chainsaw in the late 1950s; they would be felled by muscle power and the infamous “misery whip,” a two-man handsaw. Even if half the trees were ignored (trees under 12 inches in diameter, or 39.7%, being dismissed as too small and very large trees, or 5.3%, were too awkward to transport), the remaining number between 12 and 47 inches (55%) still amounts to a prodigious one-quarter billion trees cut by hand over the span of a century!

### Pragmatic Proxy

A common practice in forest management is to calculate a stand’s basal area as an approximation or proxy of its timber volume; the calculation estimates what the timber yield might be. The basal area *of a single tree* is its cross-sectional area near the ground (in square inches or square feet); *stand basal area* is the sum of all trees’ basal areas. Ideally, stem volume in a tree or in a stand requires knowledge of three dimensions: basal width, basal depth, and height; in practice, however, a single dimension – basal diameter – is sufficient for a serious beginning, because of two geometrical properties: a) tree diameter is a two-for-one dimension by simultaneously expressing both width and depth, because tree trunks are generally quasi-circular in cross section; thus its basal area is π *r*^2^, where *r* is one-half the diameter; and b) modest differences in tree height matter relatively little, so long as basal diameter is known, because the trunk’s cone-like shape contains negligible volume near its apex.

A stand’s basal area is determined correctly if its tree basal areas are worked out individually before being summed; an error occurs if the stand’s linear average diameter is simply multiplied by the number of trees in a group, because the process fails to account for the exponential effect of diameters on areas. A stand basal area the product of the number of trees in the stand and a novel “average” diameter called the *quadratic mean diameter*, or QMD (itself computed from all member’s individual basal areas). QMD is defined as *the diameter e*quivalent *of the average basal area of a group of trees*, that is, the diameter of the abstract circle that represents the average of all cross-sectional areas of the group. QMD is in a class of derived values known as “root-mean-squares” and is always somewhat larger than the comparable linear average diameter, because it incorporates the exponential effect of large-diameter specimens. The utility of QMD in forest mensuration is further explained by Curtis and Marshall (2000).

The QMD of all witness trees for SW Washington is 22.45 inches or 1.87 feet, so the total basal area of all witness trees in this study is 97,397 sq feet (π(1.87 ÷ 2)^2^ × 35,431). Of that total area, each species has a percentage share (the species’ basal area ÷ 973.97), also referred to as its relative dominance, which is illustrated as a segment of the large pie chart in Figure 12. The relative dominance of Douglas-fir, for example, is 37% of total basal area; that value exceeds the relative frequency of Douglas-fir stems (23.9% from Table 1), because Douglas-fir stems were larger in diameter than most other species. For further comparison, Figure 12 likewise illustrates relative dominances within the major forest zones, whose smaller pie charts are scaled in size according to their land areas. Without specifying wood volumes, as such, species shares shown in Figure 12 are understood to represent the approximate relative proportions of wood volume among the region’s pre-settlement forests. For example, the volume of timber in Corridor Zone, regardless of species, was larger than the volume of timber in Foothills Zone; on the other hand, the volume-yield of hemlock in Foothills was more than three times greater than the volume-yield of hemlock in Corridor (38% of 600 versus 4% of 1,500).

**Figure 12.**
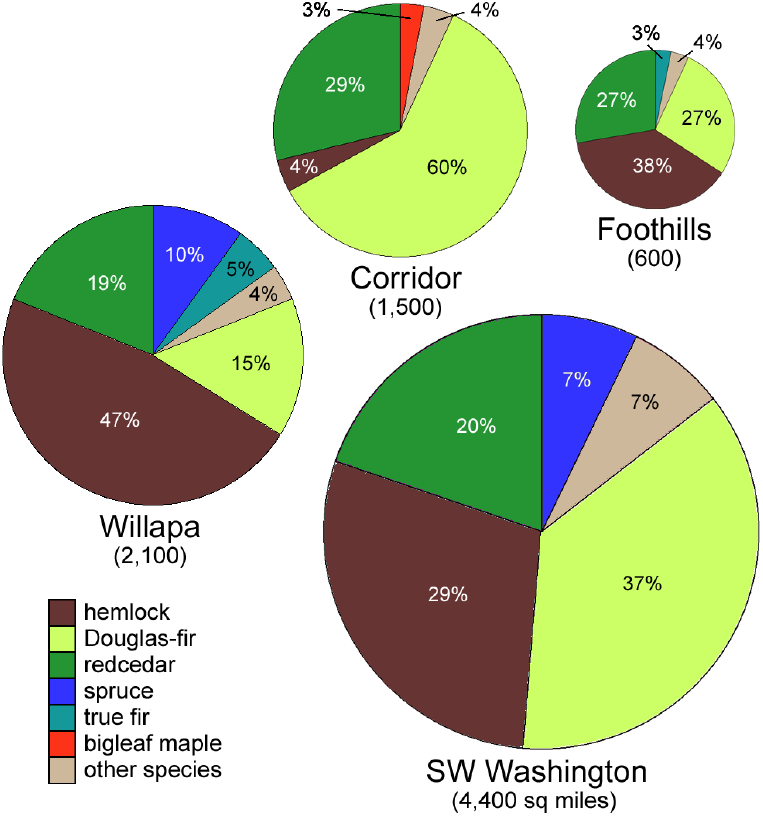
Shares of basal areas by species (also called relative dominance, a proxy for volume). Pie-chart areas are proportional to land areas.

### Lumber Numbers

Basal areas serve as proxies for wood volume only in a relative or proportional sense. The absolute measure of wood volume (or, at least, a closer approximation of such) requires knowledge of tree heights, the third dimension required for certainty. GLO records do not include tree heights, but another blanket estimate again provides the workaround: for present purposes, the average tree height for all species in all forests of SW Washington is assumed to have been 125 feet. [N. B. Like forest density, tree height functions *arithmetically* in volume computations, rather than exponentially as diameter does in basal area calculations; accordingly, if a different height value is preferred over 125 feet, then resultant volumes will scale accordingly.]

Given that the QMD of the average tree in SW Washington was 1.87 ft, its basal area would have been 2.75 sq feet (π × (1.87 ÷ 2)^2^) and its volume of trunk (as a cone) would have been 114.6 cubic feet (2.75 × 125 feet height ÷ 3). Assuming no wastage during harvesting or milling (which evidently was immense in practice and notwithstanding that paper pulp was also a major product), that average tree could have yielded 2063 lineal feet of 2×4s, or more than one-third of a mile of stud lumber. By extrapolation, therefore, the region’s full inventory of 422.4 million trees could have yielded a grand total of 165 million lineal miles of 2×4 lumber – the equivalent of 690 *lunae* of 2×4s, where one *luna* measures 238,900 miles, the distance between Earth and the Moon! Furthermore, at current retail prices of about $0.50 per lineal foot, the total commercial sale value of 2×4 lumber from SW Washington’s pre-settlement forests would be $435,600,000,000.

### Post-settlement

The notion that landed smallholders would beneficially spread socio-economic strength across America was a mainstay of pre-industrial, Jeffersonian ideology; it informed GLO’s principal operations: methodical surveys, land subdivision into equal-area plots, and public offerings of quarter-section plots (160 acres) to qualified individual applicants. The homestead process included five years of occupancy and proof of improvements (dwelling, well, access road, and some land clearing). However, the idea of homesteading did not really suit circumstances in much of SW Washington, where a settled life of farming was ostensibly not possible in the rugged uplands. Even where GLO surveyors hinted that lower forested areas might be “well adapted to agriculture when the fine timber is removed” (T19/R7W, p. 568, 1875), bitter experience soon revealed that stumpland could not be profitably managed by smallholders and that nutrient-poor forest soils were only marginally productive.

> “The most productive soil of the cutover uplands was less fertile than the least productive soil of the lowland prairies and dyked bottomlands and marshes.” (White, 1980, p. 126).

If prospects for farming were meager, the “inexhaustible” forests still held economic opportunities, just not for cash-strapped homesteaders. Commercially viable timbering required industrial-scale assets: upfront capital, access to novel technologies, partnerships with shippers, diverse knowledge and management skills; a solitary newcomer had only manual labor to bring to such an enterprise.

Early sawmills in western Washington were mostly established by absentee financiers from San Francisco, where wealth was accumulating after the Gold Rush and local demand for construction timber created a palpable market. The abrupt coastlines of California and Oregon prohibited easy access to nearby timber, so attention shifted northward to the calmer waters of Grays Harbor, Willapa Bay, and of course Puget Sound, from which products could be peacefully loaded onto ships for export. Through the 1880s, only muscle power and flotation brought logs from narrow coastal and riverbank strips to the harbor-side mills; the only mechanized component was the steam-powered saw itself. Expanding logging into the hilly hinterlands would require as-yet-unavailable mechanization of overland transport.

Savvy entrepreneurs with connections to industrial advances in eastern cities could envision industrial timbering SW Washington back country, but first they needed to acquire the land. Accordingly, in 1878 a band of industrialists and bankers convinced a naive federal legislature to pass the Timber and Mining Act: wherever land was declared unfit for agriculture and homesteading, the Act allowed a private party to acquire 160 acres at a discount price and with none of the inconvenient obligations of occupancy or improvements. The new law stimulated consortia of investors to precipitate a timber land rush – and to begin bending the laws. They found legal loopholes to exploit, land titles to be selectively ignored, and administrative oversight to be skirted. The Act’s stipulation that an applicant was limited to a single plot was readily circumvented; “dummy entrymen” were engaged to buy plots with borrowed capital and then to relinquish title back to the creditors, which often meant an invisible syndicate of finance capitalists (Beatty, 2008). Mill operators and partnering landowners avoided publicizing their identities and logging activities “because they did not want people to know anything about what they possessed or had done.” (Buchanan, 1936, p. 34).

Corporate acquisitions of largeholdings correctly anticipated innovations in mechanization. New devices for overland locomotion began to penetrate inland forests and to extract logs: rail trackage, steam donkeys, tractor crawlers, and other heavy equipment. Rising returns on investment allowed land ownership to consolidate into ever fewer hands, so by the 1890s, more than 500 sq miles of timber land in SW Washington were owned by five well-financed companies; monopolization was triumphing over distributive egalitarianism. Today, about 45% (2000 sq miles) is laced with ridge-top logging roads so that its forests can be managed as short-rotation commercial tree farms by a handful of private industrial corporate owners (Figure 13) and 15% (650 sq miles) is forest land owned by state and county agencies, whose units are also periodically logged by private contractors. The remaining 40%, mostly in Corridor Zone and valleys, is composed of small woodlots, individual farms, ranches, towns, and rural suburbs.

**Figure 13.**
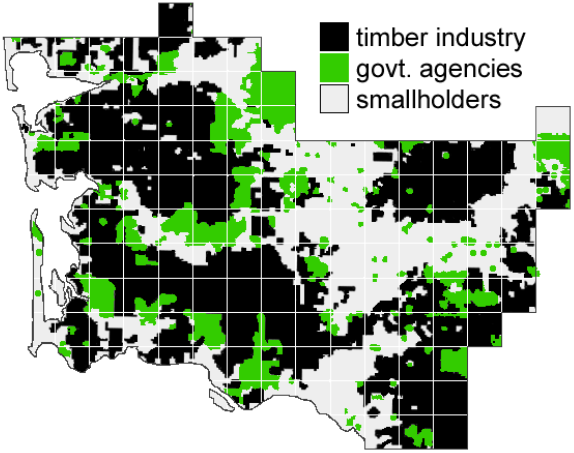
Timber production remains the region’s main land use. After Bolsinger et al. (1997), p. 8.

Incrementally, mechanized timber exploitation pushed deeper into the hinterland until, at some undated point around 1960, all the region’s original pre-settlement forests had been cutover and the natural ecosystem was set on a new trajectory. No meaningful remnant of the region’s early forests has persisted into present times, so reflections on that missing past seems quaintly plaintive:

> “What was lost to logging was not (just) forestland per se, but virgin forests. And with the passage of time even the loss was muted, for fewer and fewer Americans – increasingly urbanized creatures that they had become – could tell virgin forests from second growth or had any idea of what they were missing.” (Cox, 1983, p. 29).

[N. B. A separate downloadable file of data digests for SW Washington’s townships accompanies this article; activate ***Supplementary Material*** beneath **Download PDF** on the bioRxiv search-results page for the present title.]

## Appendix: Geophysical Context

**Figures a-e.**
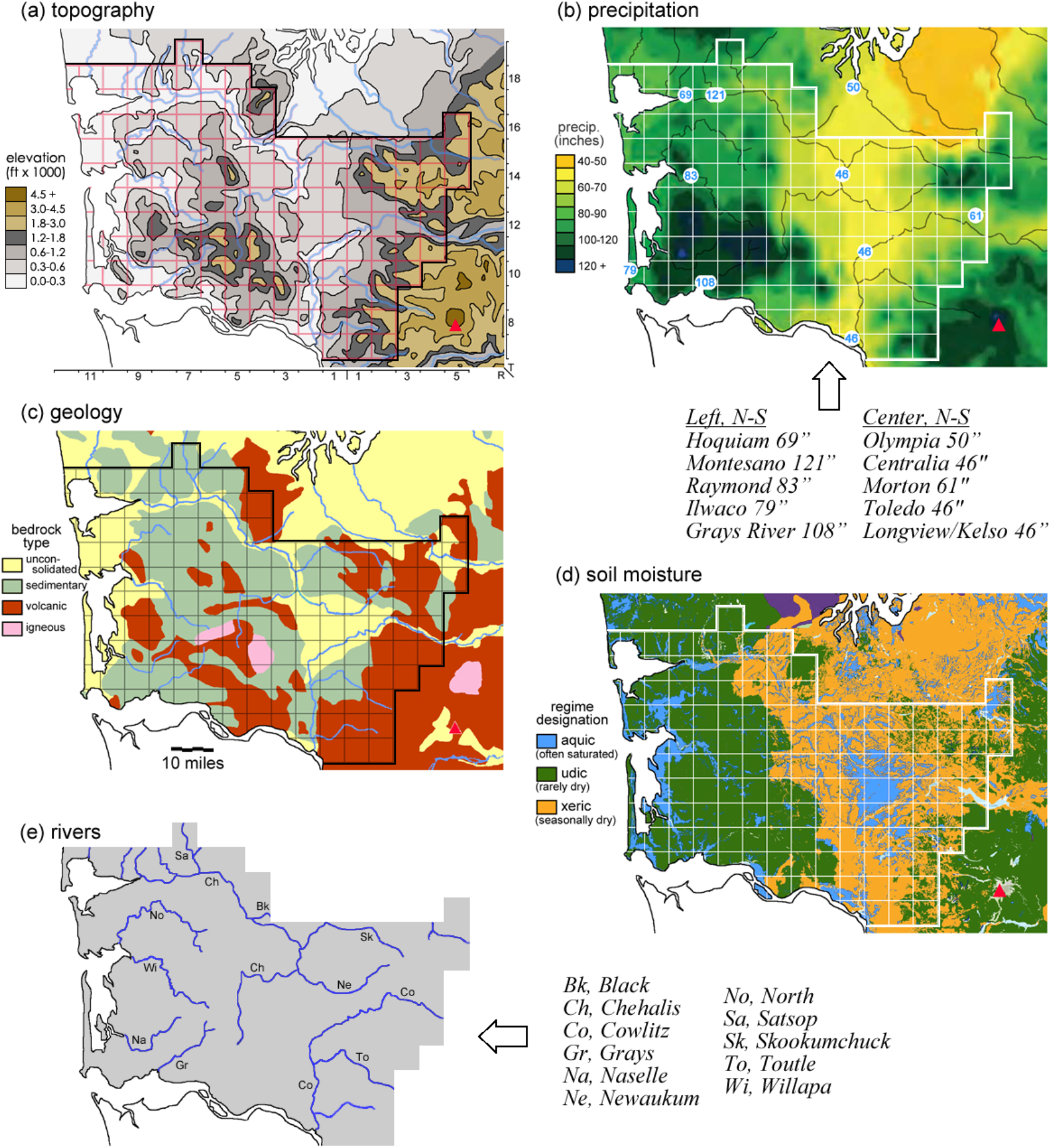
Geophysical and geo-climatic context for pre-settlement forests. Superposed township grids on T/R axes. Red triangles, Mount St. Helens. (a) after Yellowmaps (n.d.). (b) after PRISM (2000) with precipitation values for modern town sites from Wikipedia. (c) after WDNR (2010). (d) after Peterman (2012).

### Tectonics

It is well known that ancient tectonic forces sculpted major features in the Pacific Northwest, that colliding plates: a) up-thrust and buckled sea floor basalts and sedimentary layers to form the long Pacific Coast Range; b) depressed a subduction trough extending from British Columbia to Oregon; and c) elevated the Cascade Range through volcanic activity. It is less well known that the results of these geologic events in SW Washington differ considerably from their counterpart formations both to the north and the south (around Puget Sound and across the Columbia into Oregon, respectively):

- Compared to the massive Olympic Range and the imposing Oregon Coast Range, the segment of Pacific Coast Range in SW Washington is a discontinuous set of modest, eroded hills, named (with varying degrees of formal acceptance) the Willapa, Doty, Satsop, and Black Hills.
- SW Washington’s subduction trough, untouched by Ice-Age glaciers, was filled and leveled by unconsolidated layers of volcanic ejecta from nearby Mt. St. Helens (including pyroclastic ash, pumice, and lahar debris), unlike either Puget Trough that filled with seawater and glacial till or Oregon’s Willamette Valley that silted-in with remotely sourced basaltic debris during repeated Ice-Age outbursts of faraway Lake Missoula. Thus, soils of the three trough segments developed from unrelated parent materials.
- The Cascade Range adjacent SW Washington is the domain of Mts. Rainier and St. Helens, the chain’s youngest, tallest, and most active volcanoes; volcanic formations to the north and south tend to be older and more weathered.

### Geo-climatics

The hills of western SW Washington rarely rise to 2000 feet (Figure a), whereas the central lowlands are below 500 feet, and the Cascades ramparts ascend rapidly through sub-alpine to alpine areas above 4500 feet, although only their lower foothills are subjects in this study.

Precipitation follows elevation (Figure b). The western hills successfully intercept humid onshore airflow and stimulate more than 100 inches of precipitation throughout a 25-mile-wide band inland from the coast; in that band little precipitation falls as snow. Where the hills descend to the central lowlands, precipitation abruptly declines to match the rain shadow conditions of the Puget Lowlands. Further eastward, the rising Cascades again stimulate heavy precipitation, frequently as snow at higher elevations.

Onshore windstorms in SW Washington batter coastal areas and sometimes damage inland forests, especially in the western hills. The wide and deep Chehalis Gap allows moist marine air to stream into southern Puget Sound, where it contributes to the climatic peculiarity known as the Puget Sound Convergence Zone. Chehalis Gap is an Ice-Age relic resulting from a scouring slurry of outwash gravels and melt water emanating from the receding Puget Ice Lobe.

Surface runoff follows high precipitation. As illustrated in Figure (e), a few short drainage rivers in the western hills empty directly into the Pacific Ocean (Naselle, Willapa and North Rivers) or the lower Columbia River (Grays River). SW Washington’s two major watersheds generate separate stream systems that trace wide, counterclockwise paths before discharging into either Grays Harbor in the northwest (the Chehalis system) or the Columbia in the southeast (the Cowlitz system). In the first of these, the long Chehalis River aggregates streams from the western hills, meanders gently northward across the lowlands, is joined by the Newaukum and Skookumchuck Rivers from the east and the Black from the north, and finally veers westward as the main trunk to meander through the low, wide, pre-formed Chehalis Gap into Grays Harbor.

In the second large watershed, the headwaters of the Cowlitz River arise from an eponymous glacier on the eastern slopes of Mt. Rainier and the Toutle River drains the western slopes of Mt. Saint Helens. Each initially flows westward but then arcs southward; the two rivers merge to form the lower Cowlitz, which discharges into the Columbia at the modern cities of Kelso/Longview. Both rivers also experience seasonal pulses of melt water, with the result that flow in the lower Cowlitz fluctuates much more than the Chehalis.

The Cowlitz and Chehalis River systems nearly converge in the central lowlands (T12/R1W) before they abruptly pivot in opposite directions to empty 75 miles apart. The landform that prevents convergence and separates the watersheds is the ridge-like edge of a broad, plateau-like tongue of volcanic debris that declines gently from about 500 feet toward the northwest to about 150 feet. Some of its soils exhibit unusually high water-retention properties characteristic of clays (Figure d), a property it shares with silted coastal plains around Willapa Bay.

