## Supplementary Material for "Pre-Settlement Forests of Southwest Washington: Witness Statements"

Tom Schroeder

### Township Data Summaries

To augment the parent article cited above, this document provides summaries of witness tree data for the 138 townships involved in the study. Below, a blank, gridded LiDAR image of the SW Washington study area is provided as a project worksheet for transferring tree data from the following summaries.

<u>Abbreviations used in summaries</u>

**Twp/Range**: location coordinates by Township (North of Base Line in northern Oregon) and Range (East or West of Willamette Meridian)
**allWTs**: tally of all witness trees in the township
**all avDiam**: average linear diameter of all witness trees, regardless of species, in inches
**ttl QMD**: quadratic mean diameter of all witness trees in the township; (a QMD is the derived diameter of the abstract tree whose cross-sectional area is the average cross-sectional area of all trees in the set; multiple QMDs cannot be averaged, so for any new set its QMD must be computed anew from the raw linear diameters of members of the set)
**SrvyYr**: year date of the GLO survey (sometimes split over two years)
**#WTs**: number of witness trees of a given species
**DiamRange**: diameter range, from smallest to largest, within the species, in inches
**avDiam**: average linear diameter in inches
**MedianDiam**: median diameter, of which an equal number of witness was smaller and larger
**relFreq%**: relative frequency, meaning the fraction of a species among "allWTs"
**QMD**: quadratic mean diameter of witness trees of a species, in inches
**relDOM%**: relative dominance of the species, a proxy for the species' proportion of wood volume of the township's forest; relDOM% = relFreq% × ($QMD^2$ ÷ $ttlQMD^2$).

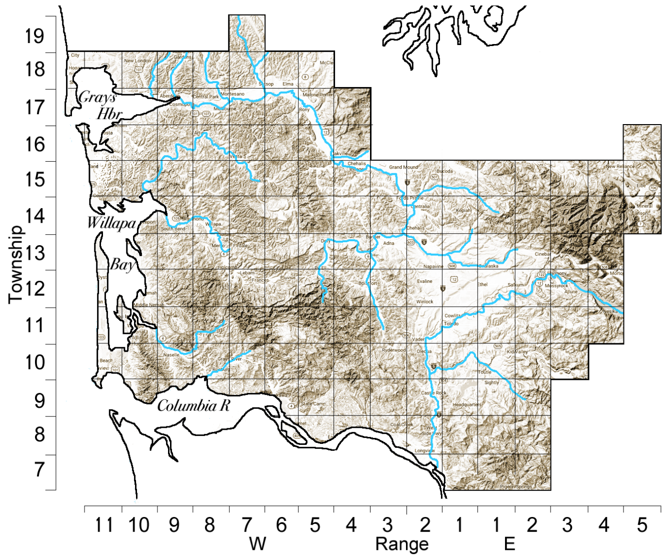

| Twp-Range | 19-7W | species | # WTs | Diam Range | av Diam | Median Diam | rel Freq% | QMD | rel DOM% |
|---|---|---|---|---|---|---|---|---|---|
| # all WTs | 288 | alder | 9 | 5to24 | 12.2 | 9 | 3.125 | 13.7 | 1.9 |
| all avDiam | 12.8 | barberry | 7 | 4to6 | 4.6 | 4 | 2.431 | 4.6 | 0.2 |
| ttl QMD | 17.7 | redcedar | 9 | 18to60 | 42.8 | 40 | 3.125 | 45.3 | 20.4 |
| SrvyYr | 1875 | Douglas-fir | 11 | 10to72 | 36.0 | 30 | 3.819 | 41.2 | 20.7 |
| | | hemlock | 145 | 4to40 | 14.4 | 10 | 50.35 | 17.2 | 47.6 |
| | | hazel | 1 | 4 | 4.0 | 4 | 0.347 | 4.0 | 0.0 |
| | | bl maple | 5 | 12to36 | 25.6 | 30 | 1.736 | 27.0 | 4.0 |
| | | spruce | 3 | 14to40 | 26.0 | 24 | 1.042 | 28.1 | 2.6 |
| | | cascara | 5 | 3to7 | 5.4 | 6 | 1.736 | 5.6 | 0.2 |
| | | v maple | 93 | 3to8 | 4.7 | 4 | 32.29 | 4.9 | 2.4 |

| Twp-Range | 18-12W | species | # WTs | Diam Range | av Diam | Median Diam | rel Freq% | QMD | rel DOM% |
|---|---|---|---|---|---|---|---|---|---|
| # all WTs | 93 | alder | 6 | 3to12 | 7.5 | 6.5 | 6.5 | 8.3 | 1.6 |
| all avDiam | 12.6 | redcedar | 3 | 48to72 | 56.0 | 48 | 3.2 | 57.1 | 37.8 |
| ttl QMD | 16.7 | crabapple | 2 | 3 | 3.0 | 3 | 2.2 | 3.0 | 0.1 |
| SrvyYr | 1858 | hemlock | 61 | 3to30 | 13.2 | 12 | 65.6 | 15.3 | 54.8 |
| | | spruce | 17 | 2to18 | 7.7 | 6 | 18.3 | 8.9 | 5.2 |
| | | willow | 4 | 3to6 | 4.3 | 4 | 4.3 | 4.4 | 0.3 |

| Twp-Range | 18-11W | species | # WTs | Diam Range | av Diam | Median Diam | rel Freq% | QMD | rel DOM% |
|---|---|---|---|---|---|---|---|---|---|
| # all WTs | 184 | alder | 11 | 6to20 | 10.6 | 9 | 6.0 | 11.5 | 1.3 |
| all avDiam | 18.5 | redcedar | 7 | 6to48 | 25.1 | 10 | 3.8 | 32.0 | 6.2 |
| ttl QMD | 25.2 | crabapple | 1 | 5 | 5.0 | 5 | 0.5 | 5.0 | 0.0 |
| SrvyYr | 1858 | hemlock | 114 | 3to48 | 16.0 | 12 | 62.0 | 19.0 | 35.3 |
| | | spruce | 37 | 4to120 | 32.7 | 24 | 20.1 | 42.4 | 56.9 |
| | | stinkwood | 1 | 4 | 4.0 | 4 | 0.5 | 4.0 | 0.0 |
| | | v maple | 9 | 4to10 | 5.3 | 5 | 4.9 | 5.6 | 0.2 |
| | | willow | 2 | 4to10 | 7.0 | 7 | 1.1 | 7.6 | 0.1 |
| | | yew | 2 | 3to4 | 3.5 | 3.5 | 1.1 | 3.5 | 0.0 |

| Twp-Range | 18-10W | species | # WTs | Diam Range | av Diam | Median Diam | rel Freq% | QMD | rel DOM% |
|---|---|---|---|---|---|---|---|---|---|
| # all WTs | 286 | alder | 6 | 3to14 | 8.3 | 1.5 | 2.1 | 9.1 | 0.4 |
| all avDiam | 16.4 | barberry | 9 | 3to10 | 5.8 | 5 | 3.1 | 6.4 | 0.3 |
| ttl QMD | 21.4 | redcedar | 6 | 4to36 | 13.7 | 10 | 2.1 | 17.3 | 1.4 |
| SrvyYr | 1858&84 | crabapple | 4 | 4to8 | 5.3 | 4.5 | 1.4 | 5.5 | 0.1 |
| | | cottonwood | 2 | 5to7 | 6.0 | 6 | 0.7 | 6.1 | 0.1 |
| | | Douglas-fir | 10 | 6to72 | 41.4 | 48 | 3.5 | 46.2 | 16.2 |
| | | hemlock | 202 | 3to48 | 15.1 | 12 | 70.6 | 17.7 | 48.0 |
| | | spruce | 33 | 3to84 | 29.0 | 24 | 11.5 | 36.4 | 33.3 |
| | | stinkwood | 1 | 14 | 14.0 | 14 | 0.3 | 14.0 | 0.1 |
| | | v maple | 13 | 2to6 | 4.2 | 4 | 4.5 | 4.4 | 0.2 |

| Twp-Range | 18-9W | species | # WTs | Diam Range | av Diam | Median Diam | rel Freq% | QMD | rel DOM% |
|---|---|---|---|---|---|---|---|---|---|
| # all WTs | 286 | alder | 18 | 4to24 | 11.6 | 9 | 6.3 | 13.2 | 2.5 |
| all avDiam | 15.8 | ash | 1 | 6 | 6.0 | 6 | 0.3 | 6.0 | 0.0 |
| ttl QMD | 20.8 | barberry | 6 | 4to6 | 5.0 | 5 | 2.1 | 5.1 | 0.1 |
| SrvyYr | 1859 | redcedar | 10 | 3to60 | 25.7 | 17 | 3.5 | 33.5 | 9.1 |
| | | crabapple | 1 | 5 | 5.0 | 5 | 0.3 | 5.0 | 0.0 |
| | | elder | 1 | 4 | 4.0 | 4 | 0.3 | 4.0 | 0.0 |
| | | hemlock | 195 | 3to60 | 16.2 | 13 | 68.2 | 19.5 | 60.2 |
| | | bl maple | 3 | 4to8 | 6.0 | 6 | 1.0 | 6.2 | 0.1 |
| | | spruce | 24 | 3to80 | 29.6 | 22 | 8.4 | 37.5 | 27.4 |
| | | v maple | 21 | 3to7 | 5.1 | 5 | 7.3 | 5.2 | 0.5 |
| | | willow | 5 | 3to5 | 3.8 | 4 | 1.7 | 3.9 | 0.1 |
| | | yellowwooc | 1 | 4 | 4.0 | 4 | 0.3 | 4.0 | 0.0 |

| Twp-Range | 18-8W | species | # WTs | Diam Range | av Diam | Median Diam | rel Freq% | QMD | rel DOM% |
|---|---|---|---|---|---|---|---|---|---|
| # all WTs | 284 | alder | 15 | 5to20 | 12.0 | 10 | 5.3 | 13.1 | 1.5 |
| all avDiam | 17.8 | barberry | 1 | 6 | 6.0 | 6 | 0.4 | 6.0 | 0.0 |
| ttl QMD | 24.3 | redcedar | 14 | 4to60 | 34.1 | 36 | 4.9 | 38.8 | 12.5 |
| SrvyYr | 1875 | cottonwood | 4 | 15to50 | 28.3 | 24 | 1.4 | 31.4 | 2.4 |
| | | Douglas-fir | 3 | 60to96 | 72.0 | 60 | 1.1 | 74.0 | 9.8 |
| | | hemlock | 161 | 3to48 | 18.1 | 16 | 56.7 | 21.4 | 44.0 |
| | | bl maple | 3 | 8to24 | 18.7 | 24 | 1.1 | 20.1 | 0.7 |
| | | spruce | 19 | 4to108 | 41.2 | 36 | 6.7 | 49.6 | 27.9 |
| | | cascara | 15 | 4to8 | 5.1 | 4 | 5.3 | 5.3 | 0.3 |
| | | v maple | 48 | 4to8 | 5.1 | 5 | 16.9 | 5.3 | 0.8 |
| | | willow | 1 | 6 | 6.0 | 6 | 0.4 | 6.0 | 0.0 |

| Twp-Range | 18-7W | species | # WTs | Diam Range | av Diam | Median Diam | rel Freq% | QMD | rel DOM% |
|---|---|---|---|---|---|---|---|---|---|
| # all WTs | 285 | alder | 20 | 5to36 | 14.7 | 14.5 | 7.0 | 16.3 | 5.6 |
| all avDiam | 14.2 | barberry | 10 | 4to7 | 5.2 | 5 | 3.5 | 5.3 | 0.3 |
| ttl QMD | 18.2 | boxwood | 1 | 4 | 4.0 | 4 | 0.4 | 4.0 | 0.0 |
| SrvyYr | 1859&75 | redcedar | 13 | 10to72 | 38.9 | 40 | 4.6 | 42.8 | 25.2 |
| | | crabapple | 1 | 3 | 3.0 | 3 | 0.4 | 3.0 | 0.0 |
| | | cottonwood | 3 | 36to50 | 40.7 | 36 | 1.1 | 41.2 | 5.4 |
| | | elder | 1 | 4 | 4.0 | 4 | 0.4 | 4.0 | 0.0 |
| | | Douglas-fir | 11 | 3to50 | 22.1 | 24 | 3.9 | 25.6 | 7.6 |
| | | hemlock | 179 | 3to50 | 13.7 | 12 | 62.8 | 16.4 | 50.7 |
| | | bl maple | 4 | 5to18 | 14.0 | 16.5 | 1.4 | 15.0 | 0.9 |
| | | oak | 3 | 8to18 | 13.3 | 14 | 1.1 | 14.0 | 0.6 |
| | | spruce | 5 | 6to36 | 20.8 | 24 | 1.8 | 23.2 | 2.8 |
| | | v maple | 33 | 3to7 | 4.4 | 4 | 11.6 | 4.5 | 0.7 |
| | | willow | 1 | 3 | 3.0 | 3 | 0.4 | 3.0 | 0.0 |

| Twp-Range | 18-6W | species | # WTs | Diam Range | av Diam | Median Diam | rel Freq% | QMD | rel DOM% |
|---|---|---|---|---|---|---|---|---|---|
| # all WTs | 282 | alder | 22 | 2to40 | 16.9 | 16.5 | 7.8 | 19.4 | 6.6 |
| all avDiam | 16.2 | ash | 3 | 15to20 | 14.3 | 18 | 1.1 | 17.8 | 0.8 |
| ttl QMD | 21.0 | barberry | 3 | 4to6 | 4.7 | 4 | 1.1 | 4.8 | 0.1 |
| SrvyYr | 1862&76 | redcedar | 28 | 5to96 | 28.0 | 27 | 9.9 | 34.4 | 26.6 |
| | | crabapple | 3 | 4to10 | 6.7 | 6 | 1.1 | 7.1 | 0.1 |
| | | cottonwood | 2 | 5to7 | 6.0 | 6 | 0.7 | 7.9 | 0.1 |
| | | dogwood | 2 | 6to8 | 7.0 | 7 | 0.7 | 7.1 | 0.1 |
| | | Douglas-fir | 27 | 6to48 | 23.6 | 19 | 9.6 | 26.9 | 15.7 |
| | | hemlock | 135 | 2to80 | 15.3 | 12 | 47.9 | 19.1 | 39.6 |
| | | hazel | 1 | 20 | 20.0 | 20 | 0.4 | 20.0 | 0.3 |
| | | bl maple | 7 | 7to48 | 20.7 | 14 | 2.5 | 25.1 | 3.5 |
| | | oak | 3 | 15to30 | 21.0 | 18 | 1.1 | 22.0 | 1.2 |
| | | spruce | 8 | 12to36 | 25.1 | 24 | 2.8 | 26.4 | 4.5 |
| | | cascara | 4 | 3to6 | 4.7 | 5 | 1.4 | 4.9 | 0.1 |
| | | v maple | 27 | 2to80 | 3.4 | 3 | 9.6 | 3.7 | 0.3 |
| | | willow | 4 | 2to10 | 4.5 | 4.5 | 1.4 | 5.7 | 0.1 |
| | | yew | 3 | 7to15 | 11.3 | 12 | 1.1 | 11.8 | 0.3 |

| Twp-Range | 18-5W | species | # WTs | Diam Range | av Diam | Median Diam | rel Freq% | QMD | rel DOM% |
|---|---|---|---|---|---|---|---|---|---|
| # all WTs | 284 | alder | 21 | 4to30 | 11.8 | 10 | 7.4 | 13.3 | 3.0 |
| all avDiam | 15.7 | ash | 5 | 8to24 | 15.2 | 12 | 1.8 | 16.3 | 1.1 |
| ttl QMD | 20.7 | redcedar | 33 | 1to70 | 28.2 | 24 | 11.6 | 33.4 | 30.3 |
| SrvyYr | 1872 | dogwood | 1 | 4 | 4.0 | 4 | 0.4 | 4.0 | 0.0 |
| | | Douglas-fir | 23 | 6to84 | 27.9 | 24 | 8.1 | 33.6 | 21.4 |
| | | hemlock | 137 | 4to48 | 15.0 | 12 | 48.2 | 18.2 | 37.3 |
| | | bl maple | 7 | 4to48 | 18.6 | 12 | 2.5 | 23.6 | 3.2 |
| | | spruce | 3 | 9to40 | 19.7 | 10 | 1.1 | 24.4 | 1.5 |
| | | cascara | 12 | 2to30 | 7.3 | 5.5 | 4.2 | 10.1 | 1.0 |
| | | v maple | 41 | 2to20 | 5.1 | 5 | 14.4 | 5.8 | 1.1 |
| | | willow | 1 | 8 | 8.0 | 8 | 0.4 | 8.0 | 0.1 |

| Twp-Range | 17-10W | species | # WTs | Diam Range | av Diam | Median Diam | rel Freq% | QMD | rel DOM% |
|---|---|---|---|---|---|---|---|---|---|
| # all WTs | 126 | alder | 11 | 5to144 | 21.8 | 6 | 8.7 | 45.4 | 32.6 |
| all avDiam | 17.8 | crabapple | 2 | 4to5 | 4.5 | 4.5 | 1.6 | 4.5 | 0.1 |
| ttl QMD | 23.5 | hemlock | 89 | 4to36 | 17.3 | 16 | 70.6 | 19.4 | 48.1 |
| SrvyYr | 1858 | spruce | 23 | 3to60 | 19.7 | 12 | 18.3 | 24.1 | 19.2 |
| | | yellowwooc | 1 | 7 | 7.0 | 7 | 0.8 | 7.0 | 0.1 |

| Twp-Range | 17-9W | species | # WTs | Diam Range | av Diam | Median Diam | rel Freq% | QMD | rel DOM% |
|---|---|---|---|---|---|---|---|---|---|
| # all WTs | 267 | alder | 26 | 3to20 | 9.4 | 8 | 9.7 | 10.4 | 2.8 |
| all avDiam | 15.2 | barberry | 3 | 6to7 | 6.3 | 6 | 1.1 | 6.4 | 0.1 |
| ttl QMD | 19.4 | redcedar | 11 | 8to80 | 38.4 | 30 | 4.1 | 44.1 | 21.3 |
| SrvyYr | 1859&82 | crabapple | 8 | 3to50 | 10.9 | 5.5 | 3.0 | 18.4 | 2.7 |
| | | elder | 1 | 5 | 5.0 | 5 | 0.4 | 5.0 | 0.0 |
| | | hemlock | 172 | 3to50 | 14.3 | 12 | 64.4 | 17.0 | 49.6 |
| | | bl maple | 1 | 22 | 22.0 | 22 | 0.4 | 22.0 | 0.5 |
| | | spruce | 38 | 4to60 | 19.9 | 15 | 14.2 | 24.3 | 22.3 |
| | | v maple | 3 | 3to4 | 3.3 | 3 | 1.1 | 3.4 | 0.0 |
| | | willow | 3 | 3to6 | 4.7 | 5 | 1.1 | 4.8 | 0.1 |
| | | yew | 1 | 24 | 24.0 | 24 | 0.4 | 24.0 | 0.6 |

| Twp-Range | 17-8W | species | # WTs | Diam Range | av Diam | Median Diam | rel Freq% | QMD | rel DOM% |
|---|---|---|---|---|---|---|---|---|---|
| # all WTs | 286 | alder | 19 | 4to30 | 10.1 | 8 | 6.6 | 11.6 | 1.7 |
| all avDiam | 17.1 | barberry | 7 | 4to10 | 6.7 | 6 | 2.4 | 6.8 | 0.2 |
| ttl QMD | 22.9 | redcedar | 9 | 6to48 | 19.8 | 14 | 3.1 | 24.7 | 3.7 |
| SrvyYr | 1859&82 | crabapple | 11 | 3to60 | 16.5 | 6 | 3.8 | 25.3 | 4.7 |
| | | cottonwood | 1 | 18 | 18.0 | 18 | 0.3 | 18.0 | 0.2 |
| | | elder | 2 | 3to7 | 5.0 | 5 | 0.7 | 5.4 | 0.0 |
| | | Douglas-fir | 2 | 40to60 | 50.0 | 50 | 0.7 | 51.0 | 3.5 |
| | | hemlock | 154 | 3to60 | 15.7 | 12 | 53.8 | 18.4 | 34.8 |
| | | bl maple | 5 | 4to24 | 12.8 | 14 | 1.7 | 15.0 | 0.8 |
| | | spruce | 54 | 6to96 | 29.0 | 16.5 | 18.9 | 37.2 | 49.9 |
| | | v maple | 10 | 3to60 | 4.3 | 4.5 | 3.5 | 4.4 | 0.1 |
| | | willow | 11 | 3to15 | 6.6 | 4 | 3.8 | 7.7 | 0.4 |
| | | yellowwooc | 1 | 6 | 6.0 | 6 | 0.3 | 6.0 | 0.0 |

| | | | | Diam | av | Median | rel | | rel |
|---|---|---|---|---|---|---|---|---|---|
| **Twp-Range** | **17-7W** | **species** | **# WTs** | **Range** | **Diam** | **Diam** | **Freq%** | **QMD** | **DOM%** |
| **# all WTs** | 285 | alder | 40 | 5to40 | 16.3 | 12 | 14.0 | 19.0 | 10.3 |
| **all avDiam** | 17.9 | ash | 4 | 10to24 | 17.0 | 17 | 1.4 | 17.8 | 0.9 |
| **ttl QMD** | 22.1 | barberry | 9 | 4to10 | 6.6 | 6 | 3.2 | 6.9 | 0.3 |
| **SrvyYr** | 1859&82 | redcedar | 15 | 6to72 | 28.5 | 28 | 5.3 | 34.4 | 12.7 |
| | | crabapple | 4 | 6to30 | 12.5 | 7 | 1.4 | 16.1 | 0.7 |
| | | cottonwood | 10 | 24to60 | 38.6 | 38 | 3.5 | 39.9 | 11.4 |
| | | elder | 2 | 3to5 | 4.0 | 4 | 0.7 | 4.1 | 0.0 |
| | | Douglas-fir | 27 | 6to60 | 24.2 | 20 | 9.5 | 27.5 | 14.7 |
| | | hemlock | 116 | 5to50 | 16.8 | 14 | 40.7 | 19.3 | 31.1 |
| | | bl maple | 9 | 4to40 | 14.6 | 8 | 3.2 | 19.4 | 2.4 |
| | | spruce | 15 | 16to60 | 34.1 | 30 | 5.3 | 36.3 | 14.2 |
| | | v maple | 14 | 4to10 | 5.9 | 6 | 4.9 | 6.2 | 0.4 |
| | | willow | 15 | 3to18 | 6.7 | 5 | 5.3 | 7.7 | 0.6 |
| | | yellowwooc | 5 | 3to8 | 5.4 | 5 | 1.8 | 5.7 | 0.1 |

| | | | | Diam | av | Median | rel | | rel |
|---|---|---|---|---|---|---|---|---|---|
| **Twp-Range** | **17-6W** | **species** | **# WTs** | **Range** | **Diam** | **Diam** | **Freq%** | **QMD** | **DOM%** |
| **# all WTs** | 282 | alder | 21 | 2to22 | 14.1 | 12 | 7.4 | 17.4 | 4.8 |
| **all avDiam** | 16.3 | ash | 5 | 2to15 | 7.8 | 6 | 1.8 | 9.2 | 0.3 |
| **ttl QMD** | 21.6 | barberry | 8 | 5to10 | 7.0 | 7 | 2.8 | 7.1 | 0.3 |
| **SrvyYr** | 1862&82 | redcedar | 35 | 6to60 | 32.9 | 40 | 12.4 | 36.8 | 35.8 |
| | | cherry | 1 | 2 | 2.0 | 2 | 0.4 | 2.0 | 0.0 |
| | | crabapple | 5 | 2to10 | 5.2 | 5 | 1.8 | 5.9 | 0.1 |
| | | cottonwood | 1 | 24 | 24.0 | 24 | 0.4 | 24.0 | 0.4 |
| | | dogwood | 3 | 6to9 | 7.7 | 8 | 1.1 | 7.8 | 0.1 |
| | | Douglas-fir | 17 | 6to40 | 22.7 | 24 | 6.0 | 24.6 | 7.8 |
| | | hemlock | 112 | 2to48 | 15.3 | 12 | 39.7 | 17.9 | 27.1 |
| | | bl maple | 14 | 6to48 | 20.7 | 15 | 5.0 | 24.4 | 6.3 |
| | | spruce | 8 | 10to96 | 43.3 | 36 | 2.8 | 51.2 | 15.9 |
| | | cascara | 7 | 3to12 | 6.0 | 4 | 2.5 | 6.8 | 0.2 |
| | | v maple | 29 | 1to8 | 4.4 | 5 | 10.3 | 4.7 | 0.5 |
| | | willow | 15 | 1to10 | 4.0 | 2 | 5.3 | 4.8 | 0.3 |
| | | yew | 1 | 8 | 8.0 | 8 | 0.4 | 8.0 | 0.0 |

| | | | | Diam | av | Median | rel | | rel |
|---|---|---|---|---|---|---|---|---|---|
| **Twp-Range** | **17-5W** | **species** | **# WTs** | **Range** | **Diam** | **Diam** | **Freq%** | **QMD** | **DOM%** |
| **# all WTs** | 280 | alder | 16 | 7to30 | 13.8 | 12 | 5.7 | 15.0 | 3.0 |
| **all avDiam** | 16.1 | ash | 1 | 12 | 12.0 | 12 | 0.4 | 12.0 | 0.1 |
| **ttl QMD** | 20.5 | barberry | 5 | 4to9 | 6.8 | 8 | 1.8 | 7.1 | 0.2 |
| **SrvyYr** | 1868&75 | redcedar | 24 | 4to72 | 29.8 | 19 | 8.6 | 36.6 | 27.3 |
| | | crabapple | 2 | 4 | 4.0 | 4 | 0.7 | 4.0 | 0.0 |
| | | cottonwood | 5 | 4to40 | 13.6 | 8 | 1.8 | 19.2 | 1.6 |
| | | dogwood | 4 | 5to9 | 6.5 | 6 | 1.4 | 6.7 | 0.2 |
| | | Douglas-fir | 44 | 5to60 | 25.9 | 24 | 15.7 | 28.0 | 29.3 |
| | | hemlock | 121 | 4to60 | 15.3 | 12 | 43.2 | 18.2 | 33.9 |
| | | bl maple | 13 | 3to30 | 12.8 | 9 | 4.6 | 15.6 | 2.7 |
| | | oak | 3 | 8to18 | 12.7 | 12 | 1.1 | 13.3 | 0.5 |
| | | spruce | 3 | 8to16 | 12.0 | 12 | 1.1 | 12.4 | 0.4 |
| | | cascara | 1 | 3 | 3.0 | 3 | 0.4 | 3.0 | 0.0 |
| | | v maple | 30 | 3to8 | 4.7 | 4 | 10.7 | 4.8 | 0.6 |
| | | willow | 8 | 4to12 | 5.9 | 5 | 2.9 | 6.4 | 0.3 |

| | | | | Diam | av | Median | rel | | rel |
|---|---|---|---|---|---|---|---|---|---|
| **Twp-Range** | **17-4W** | **species** | **# WTs** | **Range** | **Diam** | **Diam** | **Freq%** | **QMD** | **DOM%** |
| **# all WTs** | 289 | alder | 3 | 8to16 | 12.7 | 14 | 1.0 | 13.1 | 0.5 |
| **all avDiam** | 16.4 | barberry | 1 | 5 | 5.0 | 5 | 0.3 | 5.0 | 0.0 |
| **ttl QMD** | 19.6 | redcedar | 28 | 6to60 | 22.5 | 15 | 9.7 | 27.1 | 18.5 |
| **SrvyYr** | 1875 | Douglas-fir | 45 | 5to48 | 20.3 | 20 | 15.6 | 22.3 | 20.1 |
| | | true fir | 9 | 4to40 | 18.9 | 16 | 3.1 | 21.5 | 3.8 |
| | | hemlock | 181 | 3to60 | 14.8 | 12 | 62.6 | 17.6 | 50.7 |
| | | bl maple | 2 | 14to24 | 19.0 | 19 | 0.7 | 19.7 | 0.7 |
| | | spruce | 12 | 7to48 | 19.2 | 16.5 | 4.2 | 22.5 | 5.5 |
| | | v maple | 7 | 3to60 | 4.4 | 4 | 2.4 | 4.7 | 0.1 |
| | | yew | 1 | 10 | 10.0 | 10 | 0.3 | 10.0 | 0.1 |

| | | | | Diam | av | Median | rel | | rel |
|---|---|---|---|---|---|---|---|---|---|
| **Twp-Range** | **16-11W** | **species** | **# WTs** | **Range** | **Diam** | **Diam** | **Freq%** | **QMD** | **DOM%** |
| **# all WTs** | 160 | alder | 5 | 7to12 | 10.0 | 10 | 3.1 | 10.2 | 0.6 |
| **all avDiam** | 16.8 | redcedar | 11 | 8to84 | 46.4 | 60 | 6.9 | 53.4 | 38.8 |
| **ttl QMD** | 22.5 | crabapple | 1 | 4 | 4.0 | 4 | 0.6 | 4.0 | 0.0 |
| **SrvyYr** | 1858 | hemlock | 121 | 4to70 | 15.3 | 12 | 75.6 | 18.9 | 53.8 |
| | | spruce | 17 | 4to48 | 14.7 | 14 | 10.6 | 17.7 | 6.6 |
| | | cascara | 3 | 6to7 | 6.3 | 6 | 1.9 | 6.4 | 0.2 |
| | | willow | 2 | 3to4 | 3.5 | 3.5 | 1.3 | 3.5 | 0.0 |

| | | | | Diam | av | Median | rel | | rel |
|---|---|---|---|---|---|---|---|---|---|
| **Twp-Range** | **16-10W** | **species** | **# WTs** | **Range** | **Diam** | **Diam** | **Freq%** | **QMD** | **DOM%** |
| **# all WTs** | 293 | alder | 8 | 8to20 | 12.5 | 10 | 2.7 | 13.3 | 1.0 |
| **all avDiam** | 17.1 | redcedar | 34 | 5to60 | 30.3 | 30 | 11.6 | 35.0 | 30.6 |
| **ttl QMD** | 21.5 | crabapple | 1 | 6 | 6.0 | 6 | 0.3 | 6.0 | 0.0 |
| **SrvyYr** | 1858 | hemlock | 239 | 3to48 | 15.1 | 12 | 81.6 | 17.7 | 55.0 |
| | | spruce | 11 | 4to96 | 30.6 | 24 | 3.8 | 40.7 | 13.4 |

| | | | | Diam | av | Median | rel | | rel |
|---|---|---|---|---|---|---|---|---|---|
| **Twp-Range** | **16-9W** | **species** | **# WTs** | **Range** | **Diam** | **Diam** | **Freq%** | **QMD** | **DOM%** |
| **# all WTs** | 274 | alder | 20 | 8to34 | 15.6 | 14 | 7.3 | 17.0 | 5.0 |
| **all avDiam** | 15.8 | barberry | 15 | 4to9 | 6.1 | 6 | 5.5 | 6.2 | 0.5 |
| **ttl QMD** | 20.5 | redcedar | 7 | 6to120 | 46.0 | 40 | 2.6 | 60.6 | 22.4 |
| **SrvyYr** | 1881&93 | Douglas-fir | 2 | 36to60 | 48.0 | 48 | 0.7 | 49.5 | 4.3 |
| | | hemlock | 195 | 4to48 | 14.9 | 12 | 71.2 | 17.0 | 48.9 |
| | | spruce | 24 | 5to78 | 23.4 | 17.5 | 8.8 | 30.2 | 19.1 |
| | | cascara | 1 | 8 | 8.0 | 8 | 0.4 | 8.0 | 0.1 |
| | | v maple | 10 | 3to5 | 4.0 | 4 | 3.6 | 4.1 | 0.1 |

| | | | | Diam | av | Median | rel | | rel |
|---|---|---|---|---|---|---|---|---|---|
| **Twp-Range** | **16-8W** | **species** | **# WTs** | **Range** | **Diam** | **Diam** | **Freq%** | **QMD** | **DOM%** |
| **# all WTs** | 288 | alder | 14 | 5to22 | 14.9 | 15.5 | 4.9 | 15.7 | 3.6 |
| **all avDiam** | 14.3 | barberry | 21 | 4to18 | 7.2 | 6 | 7.3 | 7.9 | 1.4 |
| **ttl QMD** | 18.3 | redcedar | 4 | 5to40 | 10.9 | 6 | 1.4 | 20.6 | 1.8 |
| **SrvyYr** | 1891 | crabapple | 1 | 4 | 4.0 | 4 | 0.3 | 4.0 | 0.0 |
| | | Douglas-fir | 9 | 5to6 | 5.7 | 6 | 3.1 | 5.7 | 0.3 |
| | | hemlock | 155 | 6to60 | 16.0 | 14 | 53.8 | 18.0 | 52.1 |
| | | bl maple | 1 | 12 | 12.0 | 12 | 0.3 | 12.0 | 0.2 |
| | | spruce | 42 | 5to84 | 22.0 | 13 | 14.6 | 30.0 | 39.4 |
| | | v maple | 38 | 3to8 | 5.2 | 5 | 13.2 | 5.3 | 1.1 |
| | | willow | 3 | 4to10 | 6.3 | 5 | 1.0 | 6.9 | 0.1 |

| | | | | Diam | av | Median | rel | | rel |
|---|---|---|---|---|---|---|---|---|---|
| **Twp-Range** | **16-7W** | **species** | **# WTs** | **Range** | **Diam** | **Diam** | **Freq%** | **QMD** | **DOM%** |
| **# all WTs** | 274 | alder | 10 | 14to24 | 18.6 | 20 | 3.6 | 18.8 | 1.5 |
| **all avDiam** | 25.8 | barberry | 3 | 6to14 | 9.3 | 8 | 1.1 | 9.9 | 0.1 |
| **ttl QMD** | 29.6 | redcedar | 45 | 5to70 | 31.5 | 24 | 16.4 | 36.1 | 24.4 |
| **SrvyYr** | 1895 | Douglas-fir | 36 | 18to70 | 39.6 | 36 | 13.1 | 42.4 | 27.0 |
| | | hemlock | 142 | 6to48 | 20.8 | 20 | 51.8 | 22.0 | 28.7 |
| | | bl maple | 5 | 14to24 | 18.4 | 18 | 1.8 | 18.7 | 0.7 |
| | | spruce | 30 | 6to60 | 33.8 | 36 | 10.9 | 37.5 | 17.6 |
| | | v maple | 3 | 5to6 | 5.7 | 6 | 1.1 | 5.7 | 0.0 |

| | | | | Diam | av | Median | rel | | rel |
|---|---|---|---|---|---|---|---|---|---|
| **Twp-Range** | **16-6W** | **species** | **# WTs** | **Range** | **Diam** | **Diam** | **Freq%** | **QMD** | **DOM%** |
| **# all WTs** | 288 | alder | 2 | 15to16 | 15.5 | 15.5 | 0.7 | 15.5 | 0.2 |
| **all avDiam** | 20.4 | barberry | 3 | 4to6 | 5.3 | 6 | 1.0 | 5.4 | 0.0 |
| **ttl QMD** | 26.4 | redcedar | 19 | 5to60 | 25.7 | 15 | 6.6 | 31.6 | 9.4 |
| **SrvyYr** | 1893 | Douglas-fir | 29 | 5to95 | 42.6 | 40 | 10.1 | 46.9 | 31.8 |
| | | true fir | 5 | 10to30 | 14.4 | 10 | 1.7 | 16.4 | 0.7 |
| | | hemlock | 197 | 4to50 | 17.2 | 14 | 68.4 | 19.7 | 37.9 |
| | | bl maple | 7 | 5to24 | 11.9 | 12 | 2.4 | 13.1 | 0.6 |
| | | spruce | 11 | 4to136 | 44.7 | 30 | 3.8 | 59.1 | 19.1 |
| | | v maple | 15 | 3to8 | 4.8 | 5 | 5.2 | 5.0 | 0.2 |

| | | | | Diam | av | Median | rel | | rel |
|---|---|---|---|---|---|---|---|---|---|
| **Twp-Range** | **16-5W** | **species** | **# WTs** | **Range** | **Diam** | **Diam** | **Freq%** | **QMD** | **DOM%** |
| **# all WTs** | 262 | alder | 10 | 5to15 | 9.9 | 11 | 3.8 | 10.5 | 0.9 |
| **all avDiam** | 17.9 | ash | 7 | 6to30 | 16.6 | 12 | 2.7 | 19.2 | 2.0 |
| **ttl QMD** | 22.2 | barberry | 1 | 8 | 8.0 | 8 | 0.4 | 8.0 | 0.0 |
| **SrvyYr** | 1882 | redcedar | 42 | 5to60 | 26.5 | 25 | 16.0 | 30.4 | 30.1 |
| | | cherry | 2 | 8to9 | 8.5 | 8.5 | 0.8 | 8.5 | 0.1 |
| | | crabapple | 3 | 4to6 | 5.0 | 5 | 1.1 | 5.1 | 0.1 |
| | | cottonwood | 4 | 8to48 | 21.3 | 14.4 | 1.5 | 26.7 | 2.2 |
| | | dogwood | 1 | 9 | 9.0 | 9 | 0.4 | 9.0 | 0.1 |
| | | Douglas-fir | 54 | 1to70 | 24.5 | 20 | 20.6 | 29.0 | 35.1 |
| | | hemlock | 59 | 4to48 | 17.5 | 15 | 22.5 | 19.7 | 17.7 |
| | | bl maple | 41 | 3to36 | 13.7 | 12 | 15.6 | 15.9 | 8.0 |
| | | oak | 2 | 40 | 40.0 | 40 | 0.8 | 40.0 | 2.5 |
| | | spruce | 1 | 16 | 16.0 | 16 | 0.4 | 16.0 | 0.2 |
| | | cascara | 9 | 4to12 | 7.7 | 8 | 3.4 | 7.9 | 0.4 |
| | | v maple | 23 | 3to7 | 4.8 | 4 | 8.8 | 5.0 | 0.4 |
| | | willow | 3 | 5to12 | 8.7 | 9 | 1.1 | 9.1 | 0.2 |

| | | | | Diam | av | Median | rel | | rel |
|---|---|---|---|---|---|---|---|---|---|
| **Twp-Range** | **16-4W** | **species** | **# WTs** | **Range** | **Diam** | **Diam** | **Freq%** | **QMD** | **DOM%** |
| **# all WTs** | 284 | alder | 12 | 4to19 | 10.1 | 9.5 | 4.2 | 11.0 | 1.3 |
| **all avDiam** | 16.4 | ash | 6 | 6to20 | 12.3 | 11 | 2.1 | 13.3 | 0.9 |
| **ttl QMD** | 20.0 | barberry | 2 | 6to9 | 7.5 | 7.7 | 0.7 | 7.7 | 0.1 |
| **SrvyYr** | 1867&75 | redcedar | 34 | 6to60 | 27.0 | 24 | 12.0 | 30.5 | 27.7 |
| | | cherry | 3 | 6to8 | 7.0 | 7 | 1.1 | 7.1 | 0.1 |
| | | crabapple | 1 | 12 | 12.0 | 12 | 0.4 | 12.0 | 0.1 |
| | | cottonwood | 2 | 9to10 | 9.5 | 9.5 | 0.7 | 9.5 | 0.2 |
| | | dogwood | 5 | 4to14 | 9.0 | 10 | 1.8 | 9.6 | 0.4 |
| | | Douglas-fir | 73 | 1to56 | 21.6 | 20 | 25.7 | 25.2 | 40.8 |
| | | hemlock | 109 | 3to38 | 13.7 | 12 | 38.4 | 15.7 | 23.7 |
| | | bl maple | 10 | 4to30 | 10.3 | 6.5 | 3.5 | 12.8 | 1.4 |
| | | oak | 7 | 12to30 | 20.0 | 18 | 2.5 | 21.2 | 2.8 |
| | | v maple | 18 | 4to7 | 5.3 | 5 | 6.3 | 5.4 | 0.5 |
| | | willow | 2 | 9to10 | 9.5 | 9.5 | 0.7 | 9.5 | 0.2 |

| | | | | Diam | av | Median | rel | | rel |
|---|---|---|---|---|---|---|---|---|---|
| **Twp-Range** | **16-5E** | **species** | **# WTs** | **Range** | **Diam** | **Diam** | **Freq%** | **QMD** | **DOM%** |
| **# all WTs** | 287 | alder | 3 | 8to14 | 10.3 | 12 | 1.0 | 11.6 | 0.2 |
| **all avDiam** | 19.2 | redcedar | 45 | 5to70 | 25.9 | 20 | 15.7 | 30.8 | 26.5 |
| **ttl QMD** | 23.7 | cherry | 1 | 10 | 10.0 | 10 | 0.3 | 10.0 | 0.1 |
| **SrvyYr** | 1893&97 | cottonwood | 1 | 24 | 24.0 | 24 | 0.3 | 24.0 | 0.4 |
| | | Douglas-fir | 42 | 3to64 | 30.3 | 30 | 14.6 | 34.4 | 30.8 |
| | | true fir | 2 | 8to28 | 18.0 | 18 | 0.7 | 20.6 | 0.5 |
| | | hemlock | 176 | 3to72 | 15.4 | 12 | 61.3 | 18.6 | 37.6 |
| | | spruce | 9 | 9to60 | 21.4 | 16 | 3.1 | 26.1 | 3.8 |
| | | v maple | 3 | 3to64 | 4.4 | 4 | 1.0 | 4.5 | 0.0 |
| | | yew | 5 | 5to12 | 8.0 | 8 | 1.7 | 8.5 | 0.2 |

| | | | | Diam | av | Median | rel | | rel |
|---|---|---|---|---|---|---|---|---|---|
| **Twp-Range** | **15-11W** | **species** | **# WTs** | **Range** | **Diam** | **Diam** | **Freq%** | **QMD** | **DOM%** |
| **# all WTs** | 254 | alder | 9 | 5to16 | 8.9 | 6 | 3.5 | 9.8 | 0.6 |
| **all avDiam** | 18.1 | redcedar | 26 | 7to96 | 39.6 | 36 | 10.2 | 45.7 | 40.2 |
| **ttl QMD** | 23.1 | hemlock | 184 | 3to48 | 16.6 | 13 | 72.4 | 19.3 | 50.8 |
| **SrvyYr** | 1858&90 | pine | 4 | 6to10 | 7.5 | 7 | 1.6 | 7.7 | 0.2 |
| | | spruce | 31 | 3to60 | 13.5 | 8 | 12.2 | 19.0 | 8.2 |

| Twp-Range | 15-10W | species | # WTs | Diam Range | av Diam | Median Diam | rel Freq% | QMD | rel DOM% |
|---|---|---|---|---|---|---|---|---|---|
| # all WTs | 262 | alder | 16 | 4to20 | 10.9 | 8.5 | 6.1 | 12.0 | 2.6 |
| all avDiam | 14.4 | redcedar | 19 | 4to60 | 33.2 | 40 | 7.3 | 39.0 | 33.1 |
| ttl QMD | 18.3 | crabapple | 9 | 3to14 | 5.6 | 4 | 3.4 | 6.5 | 0.4 |
| SrvyYr | 1858 | hemlock | 185 | 3to40 | 13.3 | 10 | 70.6 | 15.5 | 50.9 |
| | | spruce | 24 | 4to60 | 16.5 | 12 | 9.2 | 21.5 | 12.7 |
| | | cascara | 1 | 6 | 6.0 | 6 | 0.4 | 6.0 | 0.0 |
| | | v maple | 2 | 4 | 4.0 | 4 | 0.8 | 4.0 | 0.0 |
| | | willow | 2 | 3to5 | 4.0 | 4 | 0.8 | 4.6 | 0.0 |
| | | yew | 4 | 4to60 | 4.8 | 5 | 1.5 | 4.6 | 0.1 |

| Twp-Range | 15-9W | species | # WTs | Diam Range | av Diam | Median Diam | rel Freq% | QMD | rel DOM% |
|---|---|---|---|---|---|---|---|---|---|
| # all WTs | 285 | alder | 10 | 4to24 | 10.4 | 8.5 | 3.5 | 11.8 | 1.2 |
| all avDiam | 16.1 | barb | 9 | 4to12 | 8.1 | 8 | 3.2 | 8.5 | 0.5 |
| ttl QMD | 20.5 | redcedar | 15 | 5to120 | 32.2 | 20 | 5.3 | 45.0 | 25.3 |
| SrvyYr | 1893 | cascara | 1 | 3 | 3.0 | 3 | 0.4 | 3.0 | 0.0 |
| | | Douglas-fir | 8 | 12to60 | 29.3 | 17 | 2.8 | 34.4 | 7.9 |
| | | true fir | 2 | 14to16 | 15.0 | 15 | 0.7 | 15.0 | 0.4 |
| | | hemlock | 208 | 3to40 | 15.2 | 12 | 73.0 | 17.4 | 52.7 |
| | | spruce | 26 | 5to72 | 18.1 | 14 | 9.1 | 23.4 | 11.9 |
| | | vine maple | 6 | 4to8 | 5.2 | 14 | 2.1 | 5.4 | 0.1 |

| Twp-Range | 15-8W | species | # WTs | Diam Range | av Diam | Median Diam | rel Freq% | QMD | rel DOM% |
|---|---|---|---|---|---|---|---|---|---|
| # all WTs | 280 | alder | 18 | 6to24 | 13.2 | 12.5 | 6.4 | 14.0 | 3.4 |
| all avDiam | 15.5 | barberry | 2 | 8to12 | 10.0 | 10 | 0.7 | 10.2 | 0.2 |
| ttl QMD | 19.3 | redcedar | 17 | 4to120 | 25.7 | 14 | 6.1 | 38.5 | 24.3 |
| SrvyYr | 1891 | Douglas-fir | 66 | 5to60 | 17.3 | 15 | 23.6 | 19.4 | 24.0 |
| | | hemlock | 118 | 4to40 | 15.0 | 13.5 | 42.1 | 16.6 | 31.3 |
| | | bl maple | 2 | 10 | 10.0 | 10 | 0.7 | 10.0 | 0.2 |
| | | spruce | 35 | 5to70 | 16.9 | 12 | 12.5 | 21.6 | 15.7 |
| | | cascara | 14 | 4to120 | 7.0 | 6 | 5.0 | 7.5 | 0.8 |
| | | v maple | 8 | 3to8 | 5.1 | 5 | 2.9 | 5.4 | 0.2 |

| Twp-Range | 15-7W | species | # WTs | Diam Range | av Diam | Median Diam | rel Freq% | QMD | rel DOM% |
|---|---|---|---|---|---|---|---|---|---|
| # all WTs | 284 | alder | 8 | 4to26 | 12.8 | 11.5 | 2.8 | 14.5 | 1.0 |
| all avDiam | 19.9 | barberry | 2 | 7to8 | 7.5 | 7.5 | 0.7 | 7.5 | 0.1 |
| ttl QMD | 24.1 | redcedar | 36 | 6to96 | 29.8 | 23 | 12.7 | 36.6 | 29.2 |
| SrvyYr | 1891 | Douglas-fir | 65 | 9to57 | 22.1 | 20 | 22.9 | 24.2 | 23.0 |
| | | hemlock | 152 | 4to42 | 17.0 | 15 | 53.5 | 18.9 | 32.8 |
| | | bl maple | 1 | 20 | 20.0 | 20 | 0.4 | 20.0 | 0.2 |
| | | spruce | 11 | 7to8 | 34.9 | 22 | 3.9 | 44.7 | 13.3 |
| | | cascara | 4 | 6to7 | 6.3 | 6 | 1.4 | 6.3 | 0.1 |
| | | v maple | 3 | 6to8 | 6.7 | 6 | 1.1 | 6.7 | 0.1 |
| | | willow | 1 | 7 | 7.0 | 7 | 0.4 | 7.0 | 0.0 |
| | | yew | 1 | 6 | 6.0 | 6 | 0.4 | 6.0 | 0.0 |

| Twp-Range | 15-6W | species | # WTs | Diam Range | av Diam | Median Diam | rel Freq% | QMD | rel DOM% |
|---|---|---|---|---|---|---|---|---|---|
| # all WTs | 264 | alder | 7 | 6to20 | 12.3 | 12 | 2.7 | 13.4 | 0.9 |
| all avDiam | 19 | redcedar | 19 | 7to72 | 29.2 | 26 | 7.2 | 33.8 | 15.1 |
| ttl QMD | 23.3 | cherry | 3 | 4to5 | 4.3 | 4 | 1.1 | 4.4 | 0.0 |
| SrvyYr | 1901 | dogwood | 1 | 4 | 4.0 | 4 | 0.4 | 4.0 | 0.0 |
| | | Douglas-fir | 41 | 8to98 | 31.2 | 28 | 15.5 | 35.6 | 36.1 |
| | | hemlock | 179 | 4to96 | 15.5 | 13 | 67.8 | 18.1 | 40.9 |
| | | bl maple | 5 | 4to17 | 10.8 | 10 | 1.9 | 11.9 | 0.5 |
| | | spruce | 8 | 8to60 | 31.1 | 30 | 3.0 | 34.1 | 6.5 |
| | | cascara | 1 | 8 | 8.0 | 8 | 0.4 | 8.0 | 0.0 |

| Twp-Range | 15-5W | species | # WTs | Diam Range | av Diam | Median Diam | rel Freq% | QMD | rel DOM% |
|---|---|---|---|---|---|---|---|---|---|
| # all WTs | 267 | alder | 14 | 4to48 | 12.5 | 10.5 | 5.2 | 16.4 | 2.0 |
| all avDiam | 21.1 | ash | 1 | 10 | 10.0 | 10 | 0.4 | 10.0 | 0.1 |
| ttl QMD | 26.7 | barberry | 3 | 4to8 | 6.3 | 7 | 1.1 | 6.6 | 0.1 |
| SrvyYr | 1876 | redcedar | 29 | 5to60 | 34.6 | 36 | 10.9 | 38.7 | 22.9 |
| | | cherry | 8 | 3to8 | 5.0 | 4 | 3.0 | 5.3 | 0.1 |
| | | crabapple | 1 | 4 | 4.0 | 4 | 0.4 | 4.0 | 0.0 |
| | | dogwood | 2 | 5to9 | 7.0 | 7 | 0.7 | 7.3 | 0.1 |
| | | Douglas-fir | 121 | 3to72 | 25.1 | 24 | 45.3 | 30.4 | 58.8 |
| | | true fir | 1 | 20 | 20.0 | 20 | 0.4 | 20.0 | 0.2 |
| | | hemlock | 49 | 4to44 | 19.4 | 18 | 18.4 | 21.7 | 12.2 |
| | | bl maple | 18 | 4to30 | 12.7 | 9 | 6.7 | 16.8 | 2.7 |
| | | spruce | 1 | 48 | 48.0 | 48 | 0.4 | 48.0 | 1.2 |
| | | cascara | 3 | 3to72 | 5.0 | 5 | 1.1 | 5.3 | 0.0 |
| | | v maple | 8 | 4to10 | 5.5 | 5 | 3.0 | 5.8 | 0.1 |
| | | willow | 8 | 3to10 | 5.9 | 5 | 3.0 | 6.4 | 0.2 |

| Twp-Range | 15-4W | species | # WTs | Diam Range | av Diam | Median Diam | rel Freq% | QMD | rel DOM% |
|---|---|---|---|---|---|---|---|---|---|
| # all WTs | 255 | alder | 23 | 1to18 | 10.6 | 10 | 9.0 | 11.5 | 1.9 |
| all avDiam | 18.0 | ash | 11 | 5to15 | 9.7 | 10 | 4.3 | 10.2 | 0.7 |
| ttl QMD | 25.0 | redcedar | 23 | 4to118 | 30.4 | 24 | 9.0 | 37.8 | 20.7 |
| SrvyYr | 1855&74 | cottonwood | 5 | 7to18 | 11.6 | 9 | 2.0 | 12.4 | 0.5 |
| | | dogwood | 7 | 4to10 | 7.4 | 7 | 2.7 | 7.8 | 0.3 |
| | | Douglas-fir | 102 | 4to72 | 23.4 | 18 | 40.0 | 28.6 | 52.5 |
| | | hemlock | 15 | 5to148 | 21.3 | 12 | 5.9 | 40.4 | 15.4 |
| | | bl maple | 50 | 3to40 | 12.1 | 9 | 19.6 | 15.3 | 7.3 |
| | | oak | 2 | 12to20 | 16.0 | 16 | 0.8 | 16.5 | 0.3 |
| | | cascara | 5 | 4to72 | 5.6 | 6 | 2.0 | 5.8 | 0.1 |
| | | v maple | 10 | 4to8 | 5.4 | 5 | 3.9 | 5.6 | 0.2 |
| | | willow | 2 | 4 | 4.0 | 4 | 0.8 | 4.0 | 0.0 |

| Twp-Range | 15-3W | species | # WTs | Diam Range | av Diam | Median Diam | rel Freq% | QMD | rel DOM% |
|---|---|---|---|---|---|---|---|---|---|
| # all WTs | 218 | alder | 16 | 5to33 | 14.9 | 12 | 7.3 | 17.0 | 4.7 |
| all avDiam | 17.4 | ash | 22 | 3to30 | 12.2 | 10 | 10.1 | 14.4 | 4.6 |
| ttl QMD | 21.3 | redcedar | 27 | 5to50 | 22.9 | 20 | 12.4 | 26.0 | 18.4 |
| SrvyYr | 1854 | cherry | 3 | 8to14 | 11.3 | 12 | 1.4 | 11.6 | 0.4 |
| | | cottonwood | 4 | 10to48 | 20.5 | 12 | 1.8 | 25.9 | 2.7 |
| | | dogwood | 3 | 4to10 | 6.7 | 6 | 1.4 | 7.1 | 0.2 |
| | | Douglas-fir | 67 | 5to60 | 23.0 | 18 | 30.7 | 27.5 | 51.3 |
| | | hemlock | 15 | 6to48 | 19.1 | 18 | 6.9 | 21.8 | 7.2 |
| | | bl maple | 46 | 5to36 | 12.2 | 10 | 21.1 | 13.8 | 8.8 |
| | | oak | 5 | 5to24 | 12.0 | 10 | 2.3 | 19.2 | 1.9 |
| | | v maple | 5 | 4to6 | 5.2 | 5 | 2.3 | 5.3 | 0.1 |
| | | willow | 3 | 10to14 | 12.0 | 12 | 1.4 | 12.1 | 0.4 |
| | | yellowwooc | 2 | 5to8 | 6.5 | 6.5 | 0.9 | 6.7 | 0.1 |

| Twp-Range | 15-2W | species | # WTs | Diam Range | av Diam | Median Diam | rel Freq% | QMD | rel DOM% |
|---|---|---|---|---|---|---|---|---|---|
| # all WTs | 270 | alder | 20 | 3to20 | 8.6 | 8.5 | 7.4 | 9.6 | 1.6 |
| all avDiam | 17.0 | ash | 8 | 7to24 | 16.5 | 17.5 | 3.0 | 17.3 | 2.1 |
| ttl QMD | 20.6 | redcedar | 80 | 5to60 | 23.7 | 24 | 29.6 | 26.6 | 49.2 |
| SrvyYr | 1855&69 | crabapple | 2 | 3to6 | 4.5 | 4.5 | 0.7 | 4.7 | 0.0 |
| | | cottonwood | 4 | 3to12 | 9.0 | 4 | 1.5 | 7.7 | 0.2 |
| | | dogwood | 1 | 7 | 7.0 | 7 | 0.4 | 7.0 | 0.0 |
| | | Douglas-fir | 98 | 4to60 | 18.4 | 15 | 36.3 | 21.6 | 39.8 |
| | | hemlock | 6 | 6to9 | 7.2 | 7 | 2.2 | 7.3 | 0.3 |
| | | bl maple | 21 | 3to18 | 9.4 | 10 | 7.8 | 10.3 | 1.9 |
| | | oak | 11 | 8to36 | 19.0 | 15 | 4.1 | 21.2 | 4.3 |
| | | hawthorn | 2 | 4 | 4.0 | 4 | 0.7 | 4.0 | 0.0 |
| | | v maple | 3 | 4to60 | 5.3 | 4 | 1.1 | 5.4 | 0.1 |
| | | willow | 14 | 3to6 | 4.7 | 4.5 | 5.2 | 4.9 | 0.3 |

| Twp-Range | 15-1W | species | # WTs | Diam Range | av Diam | Median Diam | rel Freq% | QMD | rel DOM% |
|---|---|---|---|---|---|---|---|---|---|
| # all WTs | 280 | alder | 20 | 4to20 | 8.6 | 8 | 7.1 | 9.5 | 1.5 |
| all avDiam | 15.9 | ash | 5 | 6to20 | 10.8 | 8 | 1.8 | 11.9 | 0.6 |
| ttl QMD | 20.6 | redcedar | 59 | 2to48 | 17.4 | 16 | 21.1 | 20.3 | 20.6 |
| SrvyYr | 1868 | cherry | 4 | 4to8 | 6.0 | 6 | 1.4 | 6.2 | 0.1 |
| | | crabapple | 1 | 8 | 8.0 | 8 | 0.4 | 8.0 | 0.1 |
| | | dogwood | 1 | 5 | 5.0 | 5 | 0.4 | 5.0 | 0.0 |
| | | Douglas-fir | 105 | 4to76 | 22.7 | 20 | 37.5 | 27.8 | 68.7 |
| | | true fir | 10 | 5to22 | 10.8 | 8 | 3.6 | 12.4 | 1.3 |
| | | hemlock | 31 | 2to20 | 8.0 | 8 | 11.1 | 9.0 | 2.1 |
| | | hazel | 1 | 3 | 3.0 | 3 | 0.4 | 3.0 | 0.0 |
| | | bl maple | 21 | 4to22 | 11.5 | 10 | 7.5 | 12.9 | 2.9 |
| | | oak | 2 | 15to24 | 19.5 | 19.5 | 0.7 | 20.0 | 0.7 |
| | | spruce | 1 | 20 | 20.0 | 20 | 0.4 | 20.0 | 0.3 |
| | | cascara | 3 | 3to26 | 11.3 | 15 | 1.1 | 15.4 | 0.6 |
| | | v maple | 9 | 2to8 | 3.8 | 4 | 3.2 | 4.1 | 0.1 |
| | | willow | 7 | 3to12 | 6.0 | 5 | 2.5 | 6.7 | 0.3 |

| Twp-Range | 15-1E | species | # WTs | Diam Range | av Diam | Median Diam | rel Freq% | QMD | rel DOM% |
|---|---|---|---|---|---|---|---|---|---|
| # all WTs | 288 | alder | 10 | 6to20 | 13.3 | 12 | 3.5 | 14.1 | 1.1 |
| all avDiam | 20.5 | redcedar | 43 | 6to60 | 19.8 | 15 | 14.9 | 24.3 | 14.3 |
| ttl QMD | 24.8 | cherry | 1 | 6 | 6.0 | 6 | 0.3 | 6.0 | 0.0 |
| SrvyYr | 1873 | Douglas-fir | 129 | 5to60 | 26.3 | 26 | 44.8 | 30.2 | 66.1 |
| | | true fir | 1 | 10 | 10.0 | 10 | 0.3 | 10.0 | 0.1 |
| | | hemlock | 72 | 4to44 | 16.6 | 12 | 25.0 | 19.9 | 16.1 |
| | | bl maple | 23 | 5to30 | 11.7 | 10 | 8.0 | 13.0 | 2.2 |
| | | v maple | 9 | 2to6 | 4.4 | 5 | 3.1 | 4.6 | 0.1 |

| Twp-Range | 15-2E | species | # WTs | Diam Range | av Diam | Median Diam | rel Freq% | QMD | rel DOM% |
|---|---|---|---|---|---|---|---|---|---|
| # all WTs | 286 | alder | 5 | 4to20 | 9.8 | 8 | 1.7 | 11.4 | 0.5 |
| all avDiam | 16.6 | redcedar | 46 | 3to72 | 21.9 | 18 | 16.1 | 26.1 | 26.0 |
| ttl QMD | 20.5 | cherry | 2 | 8to10 | 9.0 | 9 | 0.7 | 9.1 | 0.1 |
| SrvyYr | 1874 | Douglas-fir | 63 | 5to85 | 24.9 | 20 | 22.0 | 29.3 | 45.0 |
| | | true fir | 9 | 7to40 | 18.7 | 15 | 3.1 | 21.3 | 3.4 |
| | | hemlock | 138 | 3to40 | 12.7 | 12 | 48.3 | 14.2 | 23.2 |
| | | bl maple | 13 | 4to18 | 10.4 | 8 | 4.5 | 11.6 | 1.5 |
| | | cascara | 2 | 2to3 | 2.5 | 2.5 | 0.7 | 2.6 | 0.0 |
| | | v maple | 6 | 5to85 | 6.0 | 6 | 2.1 | 6.1 | 0.2 |
| | | willow | 1 | 4 | 4.0 | 4 | 0.3 | 4.0 | 0.0 |
| | | yew | 1 | 6 | 6.0 | 6 | 0.3 | 6.0 | 0.0 |

| Twp-Range | 15-3E | species | # WTs | Diam Range | av Diam | Median Diam | rel Freq% | QMD | rel DOM% |
|---|---|---|---|---|---|---|---|---|---|
| # all WTs | 283 | alder | 10 | 5to18 | 10.4 | 8.5 | 3.5 | 11.2 | 1.6 |
| all avDiam | 13.8 | ash | 1 | 5 | 5.0 | 5 | 0.4 | 5.0 | 0.0 |
| ttl QMD | 16.5 | redcedar | 36 | 5to40 | 17.8 | 15 | 12.7 | 20.4 | 19.3 |
| SrvyYr | 1897 | cherry | 3 | 6to8 | 6.7 | 6 | 1.1 | 6.7 | 0.2 |
| | | Douglas-fir | 128 | 3to60 | 14.9 | 12 | 45.2 | 18.1 | 53.9 |
| | | hemlock | 80 | 4to40 | 12.5 | 10 | 28.3 | 14.5 | 21.7 |
| | | bl maple | 17 | 4to16 | 9.2 | 9 | 6.0 | 9.6 | 2.0 |
| | | oak | 4 | 10to18 | 13.5 | 13 | 1.4 | 13.8 | 1.0 |
| | | pine | 2 | 5to7 | 6.0 | 6 | 0.7 | 6.1 | 0.1 |
| | | v maple | 1 | 6 | 6.0 | 6 | 0.4 | 6.0 | 0.0 |
| | | willow | 1 | 4 | 4.0 | 4 | 0.4 | 4.0 | 0.0 |

| Twp-Range | 15-4E | species | # WTs | Diam Range | av Diam | Median Diam | rel Freq% | QMD | rel DOM% |
|---|---|---|---|---|---|---|---|---|---|
| # all WTs | 285 | alder | 15 | 12to24 | 15.1 | 14 | 5.3 | 15.5 | 3.1 |
| all avDiam | 16.7 | redcedar | 25 | 6to60 | 22.6 | 20 | 8.8 | 27.4 | 16.3 |
| ttl QMD | 20.1 | cherry | 2 | 6to8 | 7.0 | 7 | 0.7 | 7.1 | 0.1 |
| SrvyYr | 1893&96 | crabapple | 1 | 10 | 10.0 | 10 | 0.4 | 10.0 | 0.1 |
| | | cottonwood | 1 | 9 | 9.0 | 9 | 0.4 | 9.0 | 0.1 |
| | | Douglas-fir | 172 | 3to60 | 16.5 | 14 | 60.4 | 19.8 | 58.5 |
| | | true fir | 3 | 8to30 | 22.7 | 30 | 1.1 | 24.9 | 1.6 |
| | | hemlock | 61 | 4to44 | 15.9 | 12 | 21.4 | 19.0 | 19.1 |
| | | bl maple | 2 | 10to30 | 20.0 | 20 | 0.7 | 22.4 | 0.9 |
| | | pine | 1 | 7 | 7.0 | 7 | 0.4 | 7.0 | 0.0 |
| | | v maple | 1 | 4 | 4.0 | 4 | 0.4 | 4.0 | 0.0 |
| | | willow | 1 | 10 | 10.0 | 10 | 0.4 | 10.0 | 0.1 |

| Twp-Range | 15-5E | species | # WTs | Diam Range | av Diam | Median Diam | rel Freq% | QMD | rel DOM% |
|---|---|---|---|---|---|---|---|---|---|
| # all WTs | 288 | alder | 24 | 5to14 | 11.2 | 12 | 8.3 | 11.5 | 2.4 |
| all avDiam | 18.3 | redcedar | 41 | 5to50 | 22.3 | 18 | 14.2 | 25.7 | 20.4 |
| ttl QMD | 21.5 | crabapple | 1 | 9 | 9.0 | 9 | 0.3 | 9.0 | 0.1 |
| SrvyYr | 1891 | cottonwood | 4 | 7to50 | 19.3 | 10 | 1.4 | 26.3 | 2.1 |
| | | Douglas-fir | 41 | 3to60 | 28.2 | 30 | 14.2 | 32.6 | 32.7 |
| | | hemlock | 163 | 4to50 | 16.2 | 14 | 56.6 | 18.0 | 39.5 |
| | | bl maple | 6 | 8to18 | 11.3 | 10 | 2.1 | 11.9 | 0.6 |
| | | spruce | 6 | 10to30 | 19.2 | 20 | 2.1 | 20.6 | 1.9 |
| | | v maple | 1 | 5 | 5.0 | 5 | 0.3 | 5.0 | 0.0 |
| | | yew | 1 | 5 | 5.0 | 5 | 0.3 | 5.0 | 0.0 |

| Twp-Range | 14-10W | species | # WTs | Diam Range | av Diam | Median Diam | rel Freq% | QMD | rel DOM% |
|---|---|---|---|---|---|---|---|---|---|
| # all WTs | 79 | alder | 4 | 6to20 | 11.8 | 10.5 | 5.1 | 13.1 | 1.3 |
| all avDiam | 17.8 | redcedar | 4 | 6to70 | 33.0 | 28 | 5.1 | 40.7 | 13.0 |
| ttl QMD | 25.4 | Douglas-fir | 3 | 6to9 | 8.0 | 9 | 3.8 | 8.1 | 0.4 |
| SrvyYr | 1855 | hemlock | 9 | 5to40 | 17.3 | 14 | 11.4 | 20.4 | 7.3 |
| | | spruce | 59 | 4to140 | 17.3 | 13 | 74.7 | 26.0 | 78.0 |

| Twp-Range | 14-9W | species | # WTs | Diam Range | av Diam | Median Diam | rel Freq% | QMD | rel DOM% |
|---|---|---|---|---|---|---|---|---|---|
| # all WTs | 272 | alder | 11 | 4to16 | 8.8 | 8 | 4.0 | 9.5 | 1.1 |
| all avDiam | 12.9 | redcedar | 7 | 6to72 | 38.1 | 30 | 2.6 | 46.5 | 16.8 |
| ttl QMD | 18.2 | crabapple | 4 | 4 | 4.0 | 4 | 1.5 | 4.0 | 0.1 |
| SrvyYr | 1858&95 | Douglas-fir | 8 | 4to20 | 8.6 | 7 | 2.9 | 9.9 | 0.9 |
| | | hemlock | 169 | 3to48 | 12.4 | 8 | 62.1 | 15.4 | 44.7 |
| | | spruce | 38 | 4to90 | 20.8 | 14.5 | 14.0 | 28.9 | 35.3 |
| | | cascara | 14 | 3to10 | 5.3 | 5 | 5.1 | 5.6 | 0.5 |
| | | v maple | 19 | 2to14 | 4.5 | 4 | 7.0 | 5.1 | 0.6 |
| | | willow | 1 | 4 | 4.0 | 4 | 0.4 | 4.0 | 0.0 |
| | | yew | 1 | 7 | 7.0 | 7 | 0.4 | 7.0 | 0.1 |

| Twp-Range | 14-8W | species | # WTs | Diam Range | av Diam | Median Diam | rel Freq% | QMD | rel DOM% |
|---|---|---|---|---|---|---|---|---|---|
| # all WTs | 290 | alder | 33 | 3to30 | 12.5 | 12 | 11.4 | 14.1 | 9.0 |
| all avDiam | 11.5 | redcedar | 8 | 4to60 | 21.4 | 19 | 2.8 | 27.8 | 8.4 |
| ttl QMD | 15.9 | cherry | 1 | 8 | 8.0 | 8 | 0.3 | 8.0 | 0.1 |
| SrvyYr | 1869&91 | crabapple | 16 | 2to7 | 4.3 | 4 | 5.5 | 4.4 | 0.4 |
| | | Douglas-fir | 29 | 5to72 | 18.7 | 14 | 10.0 | 25.6 | 25.9 |
| | | hemlock | 96 | 3to30 | 11.2 | 9.5 | 33.1 | 12.7 | 21.2 |
| | | bl maple | 2 | 4to14 | 7.0 | 7 | 0.7 | 10.3 | 0.3 |
| | | spruce | 40 | 3to72 | 18.7 | 13 | 13.8 | 24.4 | 32.5 |
| | | cascara | 26 | 3to9 | 5.0 | 5 | 9.0 | 5.2 | 1.0 |
| | | v maple | 37 | 3to8 | 4.2 | 4 | 12.8 | 4.4 | 1.0 |
| | | willow | 2 | 5to10 | 7.5 | 7.5 | 0.7 | 7.9 | 0.2 |

| Twp-Range | 14-7W | species | # WTs | Diam Range | av Diam | Median Diam | rel Freq% | QMD | rel DOM% |
|---|---|---|---|---|---|---|---|---|---|
| # all WTs | 285 | alder | 12 | 4to20 | 13.4 | 15 | 4.2 | 14.5 | 1.9 |
| all avDiam | 16.2 | barberry | 11 | 4to10 | 6.0 | 5 | 3.9 | 6.3 | 0.3 |
| ttl QMD | 21.7 | redcedar | 47 | 5to120 | 28.6 | 18 | 16.5 | 38.0 | 50.6 |
| SrvyYr | 1878 | Douglas-fir | 21 | 6to60 | 27.0 | 24 | 7.4 | 30.9 | 15.0 |
| | | true fir | 4 | 8to24 | 14.0 | 12 | 1.4 | 15.4 | 0.7 |
| | | hemlock | 167 | 4to36 | 12.9 | 12 | 58.6 | 14.4 | 25.8 |
| | | spruce | 6 | 14to60 | 29.7 | 24 | 2.1 | 33.9 | 5.1 |
| | | cascara | 5 | 4to5 | 4.4 | 4 | 1.8 | 4.4 | 0.1 |
| | | v maple | 10 | 3to8 | 4.6 | 4 | 3.5 | 4.8 | 0.2 |
| | | yew | 2 | 5to16 | 10.5 | 10.5 | 0.7 | 11.8 | 0.2 |

| Twp-Range | 14-6W | species | # WTs | Diam Range | av Diam | Median Diam | rel Freq% | QMD | rel DOM% |
|---|---|---|---|---|---|---|---|---|---|
| # all WTs | 286 | alder | 4 | 10to24 | 18.0 | 19 | 1.4 | 18.7 | 0.8 |
| all avDiam | 19.4 | barberry | 2 | 4to7 | 5.5 | 5.5 | 0.7 | 5.7 | 0.0 |
| ttl QMD | 24.1 | redcedar | 24 | 4to72 | 35.1 | 36 | 8.4 | 38.9 | 21.9 |
| SrvyYr | 1895 | Douglas-fir | 26 | 10to62 | 36.4 | 36 | 9.1 | 41.0 | 26.4 |
| | | hemlock | 219 | 4to54 | 15.3 | 12 | 76.6 | 18.0 | 42.8 |
| | | spruce | 9 | 14to60 | 35.8 | 30 | 3.1 | 38.2 | 7.9 |
| | | v maple | 1 | 4 | 4.0 | 4 | 0.3 | 4.0 | 0.0 |
| | | yew | 1 | 8 | 8.0 | 8 | 0.3 | 8.0 | 0.0 |

| Twp-Range | 14-5W | species | # WTs | Diam Range | av Diam | Median Diam | rel Freq% | QMD | rel DOM% |
|---|---|---|---|---|---|---|---|---|---|
| # all WTs | 289 | alder | 2 | 15to16 | 15.5 | 15.5 | 0.7 | 15.5 | 0.2 |
| all avDiam | 25.3 | barberry | 1 | 8 | 8.0 | 8 | 0.3 | 8.0 | 0.0 |
| ttl QMD | 29.3 | redcedar | 56 | 10to72 | 36.0 | 36 | 19.4 | 39.1 | 34.5 |
| SrvyYr | 1901 | dogwood | 3 | 8to10 | 9.3 | 10 | 1.0 | 9.4 | 0.1 |
| | | Douglas-fir | 57 | 10to76 | 32.8 | 30 | 19.7 | 35.2 | 28.6 |
| | | true fir | 1 | 8 | 8.0 | 8 | 0.3 | 8.0 | 0.0 |
| | | hemlock | 153 | 5to53 | 20.1 | 18 | 52.9 | 22.9 | 32.4 |
| | | larch | 5 | 8to44 | 20.0 | 12 | 1.7 | 24.5 | 1.2 |
| | | bl maple | 6 | 3to18 | 7.7 | 5 | 2.1 | 9.5 | 0.2 |
| | | spruce | 3 | 33to64 | 44.3 | 36 | 1.0 | 46.5 | 2.6 |
| | | v maple | 2 | 6to8 | 7.0 | 7 | 0.7 | 7.1 | 0.0 |

| Twp-Range | 14-4W | species | # WTs | Diam Range | av Diam | Median Diam | rel Freq% | QMD | rel DOM% |
|---|---|---|---|---|---|---|---|---|---|
| # all WTs | 288 | alder | 21 | 1to36 | 9.9 | 8 | 7.3 | 12.3 | 1.7 |
| all avDiam | 21.2 | barberry | 2 | 5to8 | 6.5 | 6.5 | 0.7 | 6.7 | 0.0 |
| ttl QMD | 25.7 | redcedar | 67 | 6to70 | 24.9 | 24 | 23.3 | 28.2 | 27.9 |
| SrvyYr | 1872 | cherry | 1 | 6 | 6.0 | 6 | 0.3 | 6.0 | 0.0 |
| | | dogwood | 1 | 6 | 6.0 | 6 | 0.3 | 6.0 | 0.0 |
| | | Douglas-fir | 130 | 4to70 | 25.8 | 24 | 45.1 | 30.2 | 62.2 |
| | | true fir | 1 | 40 | 40.0 | 40 | 0.3 | 40.0 | 0.8 |
| | | hemlock | 39 | 3to36 | 13.1 | 10 | 13.5 | 15.3 | 4.8 |
| | | bl maple | 24 | 4to36 | 11.4 | 10 | 8.3 | 14.3 | 2.6 |
| | | v maple | 2 | 4to8 | 6.0 | 6 | 0.7 | 6.3 | 0.0 |

| Twp-Range | 14-3W | species | # WTs | Diam Range | av Diam | Median Diam | rel Freq% | QMD | rel DOM% |
|---|---|---|---|---|---|---|---|---|---|
| # all WTs | 277 | alder | 11 | 4to12 | 14.2 | 12 | 4.0 | 15.8 | 1.7 |
| all avDiam | 19.3 | ash | 4 | 10to15 | 12.3 | 12 | 1.4 | 12.4 | 0.4 |
| ttl QMD | 24.2 | redcedar | 85 | 4to96 | 27.0 | 24 | 30.7 | 31.4 | 51.5 |
| SrvyYr | 1857 | cherry | 1 | 14 | 14.0 | 14 | 0.4 | 14.0 | 0.1 |
| | | Douglas-fir | 97 | 4to72 | 19.0 | 14 | 35.0 | 24.4 | 35.5 |
| | | true fir | 5 | 7to40 | 16.2 | 10 | 1.8 | 20.3 | 1.3 |
| | | hemlock | 33 | 3to36 | 13.3 | 10 | 11.9 | 15.8 | 5.1 |
| | | bl maple | 27 | 4to30 | 12.9 | 12 | 9.7 | 14.4 | 3.4 |
| | | oak | 2 | 12to30 | 21.0 | 21 | 0.7 | 22.9 | 0.6 |
| | | v maple | 7 | 5to12 | 7.0 | 6 | 2.5 | 7.3 | 0.2 |
| | | willow | 2 | 6to10 | 8.0 | 8 | 0.7 | 8.3 | 0.1 |
| | | yellowwooc | 2 | 4to6 | 5.0 | 5 | 0.7 | 5.1 | 0.0 |
| | | yew | 1 | 8 | 8.0 | 8 | 0.4 | 8.0 | 0.0 |

| Twp-Range | 14-2W | species | # WTs | Diam Range | av Diam | Median Diam | rel Freq% | QMD | rel DOM% |
|---|---|---|---|---|---|---|---|---|---|
| # all WTs | 277 | alder | 20 | 3to18 | 9.5 | 7 | 7.2 | 10.4 | 2.2 |
| all avDiam | 16.0 | ash | 19 | 6to24 | 12.6 | 14 | 6.9 | 13.5 | 3.5 |
| ttl QMD | 18.9 | redcedar | 93 | 6to46 | 18.7 | 16 | 33.6 | 20.8 | 40.9 |
| SrvyYr | 1855 | cherry | 1 | 6 | 6.0 | 6 | 0.4 | 6.0 | 0.0 |
| | | crabapple | 4 | 3to7 | 5.0 | 5 | 1.4 | 5.2 | 0.1 |
| | | cottonwood | 2 | 3to24 | 13.5 | 13.5 | 0.7 | 17.1 | 0.6 |
| | | Douglas-fir | 83 | 3to50 | 19.1 | 16 | 30.0 | 22.2 | 41.7 |
| | | hemlock | 16 | 4to36 | 9.6 | 8 | 5.8 | 12.0 | 2.3 |
| | | bl maple | 21 | 3to28 | 12.2 | 12 | 7.6 | 13.6 | 4.0 |
| | | oak | 10 | 9to36 | 18.2 | 12 | 3.6 | 21.0 | 4.5 |
| | | v maple | 5 | 3to6 | 4.6 | 4 | 1.8 | 4.8 | 0.1 |
| | | willow | 3 | 3to9 | 5.0 | 3 | 1.1 | 5.7 | 0.1 |

| Twp-Range | 14-1W | species | # WTs | Diam Range | av Diam | Median Diam | rel Freq% | QMD | rel DOM% |
|---|---|---|---|---|---|---|---|---|---|
| # all WTs | 288 | alder | 16 | 4to36 | 13.0 | 10 | 5.6 | 15.6 | 2.4 |
| all avDiam | 16.9 | ash | 4 | 8to24 | 13.5 | 11 | 1.4 | 14.9 | 0.6 |
| ttl QMD | 23.5 | redcedar | 64 | 3to48 | 16.9 | 15.5 | 22.2 | 19.5 | 15.3 |
| SrvyYr | 1868 | cherry | 8 | 2to5 | 3.6 | 4 | 2.8 | 3.7 | 0.1 |
| | | crabapple | 2 | 4to6 | 5.0 | 5 | 0.7 | 5.1 | 0.0 |
| | | dogwood | 1 | 8 | 8.0 | 8 | 0.3 | 8.0 | 0.0 |
| | | Douglas-fir | 112 | 2to120 | 24.0 | 20 | 38.9 | 31.9 | 71.6 |
| | | hemlock | 31 | 2to28 | 11.2 | 8 | 10.8 | 13.7 | 3.6 |
| | | bl maple | 30 | 2to80 | 11.3 | 7.5 | 10.4 | 18.1 | 6.1 |
| | | cascara | 2 | 3to8 | 5.5 | 5.5 | 0.7 | 6.0 | 0.0 |
| | | v maple | 12 | 2to6 | 4.1 | 4 | 4.2 | 4.3 | 0.1 |
| | | willow | 4 | 2to3 | 2.8 | 3 | 1.4 | 2.8 | 0.0 |
| | | yew | 2 | 5to6 | 5.5 | 5.5 | 0.694 | 5.52 | 0.0382 |

| Twp-Range | 14-1E | species | # WTs | Diam Range | av Diam | Median Diam | rel Freq% | QMD | rel DOM% |
|---|---|---|---|---|---|---|---|---|---|
| # all WTs | 272 | alder | 2 | 7to14 | 10.5 | 10.5 | 0.7 | 11.1 | 0.2 |
| all avDiam | 18.3 | ash | 1 | 24 | 24.0 | 24 | 0.4 | 24.0 | 0.4 |
| ttl QMD | 23.4 | barberry | 1 | 5 | 5.0 | 5 | 0.4 | 5.0 | 0.0 |
| SrvyYr | 1873 | redcedar | 33 | 3to70 | 23.9 | 14 | 12.1 | 31.3 | 21.7 |
| | | cherry | 4 | 6 | 6.0 | 6 | 1.5 | 6.0 | 0.1 |
| | | Douglas-fir | 61 | 4to72 | 29.9 | 28 | 22.4 | 34.0 | 47.5 |
| | | true fir | 3 | 10to18 | 13.7 | 13 | 1.1 | 15.2 | 0.5 |
| | | hemlock | 132 | 4to40 | 15.3 | 12 | 48.5 | 17.8 | 28.2 |
| | | bl maple | 14 | 3to20 | 9.1 | 6.5 | 5.1 | 10.8 | 1.1 |
| | | cascara | 1 | 4 | 4.0 | 4 | 0.4 | 4.0 | 0.0 |
| | | v maple | 20 | 3to6 | 4.7 | 5 | 7.4 | 4.8 | 0.3 |

| Twp-Range | 14-2E | species | # WTs | Diam Range | av Diam | Median Diam | rel Freq% | QMD | rel DOM% |
|---|---|---|---|---|---|---|---|---|---|
| # all WTs | 287 | alder | 3 | 4to14 | 8.0 | 6 | 1.0 | 9.1 | 0.2 |
| all avDiam | 16.4 | redcedar | 7 | 3to50 | 28.4 | 24 | 2.4 | 34.7 | 6.6 |
| ttl QMD | 21.2 | Douglas-fir | 39 | 4to92 | 17.5 | 10 | 13.6 | 27.0 | 22.1 |
| SrvyYr | 1896 | true fir | 62 | 4to70 | 16.7 | 12 | 21.6 | 21.0 | 21.3 |
| | | hemlock | 173 | 4to78 | 15.8 | 12 | 60.3 | 19.2 | 49.7 |
| | | v maple | 3 | 3to6 | 4.7 | 5 | 1.0 | 4.8 | 0.1 |

| Twp-Range | 14-3E | species | # WTs | Diam Range | av Diam | Median Diam | rel Freq% | QMD | rel DOM% |
|---|---|---|---|---|---|---|---|---|---|
| # all WTs | 290 | alder | 1 | 18 | 18.0 | 18 | 0.3 | 18.0 | 0.6 |
| all avDiam | 11.1 | redcedar | 4 | 6to8 | 7.3 | 6.5 | 1.4 | 7.3 | 0.4 |
| ttl QMD | 13.2 | Douglas-fir | 215 | 4to48 | 10.9 | 10 | 74.1 | 12.9 | 71.0 |
| SrvyYr | 1910 | true fir | 4 | 5to16 | 10.3 | 10 | 1.4 | 11.0 | 0.9 |
| | | hemlock | 57 | 4to50 | 12.3 | 10 | 19.7 | 15.0 | 25.4 |
| | | larch | 5 | 6to10 | 8.0 | 8 | 1.7 | 8.2 | 0.7 |
| | | pine | 4 | 6to18 | 9.8 | 7.5 | 1.4 | 10.9 | 0.9 |

| Twp-Range | 14-4E | species | # WTs | Diam Range | av Diam | Median Diam | rel Freq% | QMD | rel DOM% |
|---|---|---|---|---|---|---|---|---|---|
| # all WTs | 281 | alder | 1 | 18 | 18.0 | 18 | 0.4 | 18.0 | 0.3 |
| all avDiam | 18.3 | redcedar | 9 | 10to38 | 19.6 | 20 | 3.2 | 21.2 | 3.8 |
| ttl QMD | 19.6 | crabapple | 1 | 10 | 10.0 | 10 | 0.4 | 10.0 | 0.1 |
| SrvyYr | 1906 | Douglas-fir | 186 | 6to32 | 16.8 | 18 | 66.2 | 17.6 | 53.5 |
| | | true fir | 10 | 8to60 | 24.3 | 19.5 | 3.6 | 29.1 | 7.9 |
| | | hemlock | 65 | 10to36 | 22.2 | 22 | 23.1 | 23.1 | 32.2 |
| | | pine | 6 | 12to20 | 16.5 | 17 | 2.1 | 16.7 | 1.6 |
| | | v maple | 1 | 5 | 5.0 | 5 | 0.4 | 5.0 | 0.0 |
| | | yew | 2 | 20 | 20.0 | 20 | 0.7 | 20.0 | 0.7 |

| Twp-Range | 14-5E | species | # WTs | Diam Range | av Diam | Median Diam | rel Freq% | QMD | rel DOM% |
|---|---|---|---|---|---|---|---|---|---|
| # all WTs | 286 | alder | 19 | 4to20 | 11.2 | 12 | 6.6 | 11.8 | 3.8 |
| all avDiam | 13.2 | barberry | 1 | 6 | 6.0 | 6 | 0.3 | 6.0 | 0.1 |
| ttl QMD | 15.6 | redcedar | 25 | 5to60 | 18.6 | 12 | 8.7 | 23.0 | 19.1 |
| SrvyYr | 1894 | crabapple | 1 | 10 | 10.0 | 10 | 0.3 | 10.0 | 0.1 |
| | | cottonwood | 1 | 20 | 20.0 | 20 | 0.3 | 20.0 | 0.6 |
| | | Douglas-fir | 100 | 4to48 | 13.5 | 12 | 35.0 | 15.5 | 34.8 |
| | | true fir | 11 | 4to14 | 9.3 | 8 | 3.8 | 9.6 | 1.5 |
| | | hemlock | 115 | 4to70 | 13.1 | 12 | 40.2 | 15.3 | 38.8 |
| | | bl maple | 5 | 8to14 | 10.4 | 10 | 1.7 | 10.6 | 0.8 |
| | | pine | 1 | 6 | 6.0 | 6 | 0.3 | 6.0 | 0.1 |
| | | v maple | 4 | 4to6 | 5.3 | 5.5 | 1.4 | 5.3 | 0.2 |
| | | willow | 2 | 4 | 4.0 | 4 | 0.7 | 4.0 | 0.0 |
| | | yew | 1 | 10 | 10.0 | 10 | 0.3 | 10.0 | 0.1 |

| Twp-Range | 13-10W | species | # WTs | Diam Range | av Diam | Median Diam | rel Freq% | QMD | rel DOM% |
|---|---|---|---|---|---|---|---|---|---|
| # all WTs | 202 | alder | 8 | 4to15 | 8.3 | 7 | 4.0 | 9.2 | 0.6 |
| all avDiam | 19.2 | redcedar | 10 | 6to75 | 36.7 | 4 | 5.0 | 41.7 | 15.2 |
| ttl QMD | 23.8 | crabapple | 5 | 4to8 | 5.2 | 4 | 2.5 | 5.4 | 0.1 |
| SrvyYr | 1855&58 | hemlock | 85 | 6to86 | 23.8 | 24 | 42.1 | 27.5 | 56.3 |
| | | pine | 4 | 4to60 | 4.8 | 4.5 | 2.0 | 4.8 | 0.1 |
| | | spruce | 88 | 4to60 | 15.6 | 12 | 43.6 | 18.9 | 27.6 |
| | | willow | 2 | 4to5 | 4.5 | 4.5 | 1.0 | 4.5 | 0.0 |

| Twp-Range | 13-9W | species | # WTs | Diam Range | av Diam | Median Diam | rel Freq% | QMD | rel DOM% |
|---|---|---|---|---|---|---|---|---|---|
| # all WTs | 289 | alder | 1 | 18 | 18.0 | 18 | 0.3 | 18.0 | 0.3 |
| all avDiam | 16.5 | redcedar | 9 | 12to72 | 38.4 | 36 | 3.1 | 44.4 | 14.5 |
| ttl QMD | 20.6 | Douglas-fir | 9 | 6to40 | 16.0 | 12 | 3.1 | 19.3 | 2.7 |
| SrvyYr | 1891 | true fir | 1 | 10 | 10.0 | 10 | 0.3 | 10.0 | 0.1 |
| | | hemlock | 227 | 3to44 | 14.9 | 12 | 78.5 | 17.7 | 57.8 |
| | | spruce | 38 | 4to84 | 22.7 | 20 | 13.1 | 28.2 | 24.6 |
| | | cascara | 1 | 5 | 5.0 | 5 | 0.3 | 5.0 | 0.0 |
| | | v maple | 3 | 4to84 | 5.3 | 4 | 1.0 | 5.7 | 0.1 |

| Twp-Range | 13-8W | species | # WTs | Diam Range | av Diam | Median Diam | rel Freq% | QMD | rel DOM% |
|---|---|---|---|---|---|---|---|---|---|
| # all WTs | 287 | alder | 16 | 4to24 | 10.7 | 7 | 5.6 | 15.2 | 3.4 |
| all avDiam | 13.0 | redcedar | 5 | 12to36 | 22.4 | 20 | 1.7 | 24.2 | 2.7 |
| ttl QMD | 19.5 | crabapple | 3 | 3to6 | 4.4 | 4 | 1.0 | 4.5 | 0.1 |
| SrvyYr | 1869&81 | dogwood | 1 | 4 | 4.0 | 4 | 0.3 | 4.0 | 0.0 |
| | | Douglas-fir | 46 | 3to72 | 22.1 | 6 | 16.0 | 31.9 | 42.8 |
| | | true fir | 4 | 7to24 | 13.5 | 11.5 | 1.4 | 15.1 | 0.8 |
| | | hemlock | 108 | 3to50 | 13.3 | 9.5 | 37.6 | 16.5 | 26.8 |
| | | bl maple | 1 | 14 | 14.0 | 14 | 0.3 | 14.0 | 0.2 |
| | | spruce | 21 | 4to72 | 25.1 | 14 | 7.3 | 33.5 | 21.5 |
| | | cascara | 6 | 3to10 | 5.8 | 4 | 2.1 | 6.6 | 0.2 |
| | | v maple | 67 | 3to10 | 4.6 | 4 | 23.3 | 4.7 | 1.4 |
| | | willow | 9 | 3to8 | 5.0 | 4 | 3.1 | 5.2 | 0.2 |

| Twp-Range | 13-7W | species | # WTs | Diam Range | av Diam | Median Diam | rel Freq% | QMD | rel DOM% |
|---|---|---|---|---|---|---|---|---|---|
| # all WTs | 290 | alder | 5 | 8to30 | 16.4 | 15 | 1.7 | 17.9 | 1.0 |
| all avDiam | 16.7 | barberry | 8 | 4to10 | 7.1 | 6.5 | 2.8 | 7.4 | 0.3 |
| ttl QMD | 23.0 | redcedar | 30 | 8to96 | 35.5 | 33 | 10.3 | 40.6 | 32.2 |
| SrvyYr | 1867 | cherry | 1 | 10 | 10.0 | 10 | 0.3 | 10.0 | 0.1 |
| | | crabapple | 1 | 4 | 4.0 | 4 | 0.3 | 4.0 | 0.0 |
| | | elder | 1 | 6 | 6.0 | 6 | 0.3 | 6.0 | 0.0 |
| | | Douglas-fir | 22 | 10to70 | 38.4 | 36 | 7.6 | 40.6 | 23.6 |
| | | hemlock | 121 | 3to48 | 13.3 | 10 | 41.7 | 15.7 | 19.3 |
| | | bl maple | 11 | 4to24 | 14.6 | 14 | 3.8 | 16.3 | 1.9 |
| | | spruce | 20 | 10to72 | 33.7 | 24 | 6.9 | 39.5 | 20.4 |
| | | v maple | 68 | 3to10 | 4.6 | 4 | 23.4 | 4.7 | 1.0 |
| | | willow | 2 | 5to7 | 6.0 | 6 | 0.7 | 6.1 | 0.0 |

| Twp-Range | 13-6W | species | # WTs | Diam Range | av Diam | Median Diam | rel Freq% | QMD | rel DOM% |
|---|---|---|---|---|---|---|---|---|---|
| # all WTs | 286 | alder | 5 | 6to24 | 10.8 | 6 | 1.7 | 12.9 | 0.7 |
| all avDiam | 15.9 | bearberry | 2 | 7 | 7.0 | 7 | 0.7 | 7.0 | 0.1 |
| ttl QMD | 20.2 | redcedar | 19 | 13to84 | 30.1 | 30 | 6.6 | 33.8 | 18.6 |
| SrvyYr | 1890 | Douglas-fir | 17 | 5to76 | 35.3 | 31 | 5.9 | 41.6 | 25.3 |
| | | hemlock | 202 | 3to48 | 14.4 | 12 | 70.6 | 16.7 | 48.4 |
| | | bl maple | 2 | 9to20 | 14.5 | 14.5 | 0.7 | 15.5 | 0.4 |
| | | spruce | 6 | 10to48 | 30.7 | 30 | 2.1 | 33.0 | 5.6 |
| | | v maple | 33 | 3to12 | 5.2 | 5 | 11.5 | 5.5 | 0.9 |

| Twp-Range | 13-5W | species | # WTs | Diam Range | av Diam | Median Diam | rel Freq% | QMD | rel DOM% |
|---|---|---|---|---|---|---|---|---|---|
| # all WTs | 293 | alder | 8 | 7to28 | 16.1 | 14 | 2.7 | 17.6 | 1.2 |
| all avDiam | 21.4 | ash | 10 | 4to24 | 11.2 | 10 | 3.4 | 12.5 | 0.7 |
| ttl QMD | 27.1 | redcedar | 34 | 3to84 | 31.2 | 30 | 11.6 | 36.2 | 20.7 |
| SrvyYr | 1855&75 | cherry | 4 | 6to10 | 8.0 | 8 | 1.4 | 8.1 | 0.1 |
| | | crabapple | 2 | 6to9 | 7.5 | 7.5 | 0.7 | 7.7 | 0.1 |
| | | cottonwood | 5 | 4to16 | 12.0 | 13 | 1.7 | 12.7 | 0.4 |
| | | dogwood | 3 | 8to12 | 10.7 | 8 | 1.0 | 9.5 | 0.1 |
| | | Douglas-fir | 103 | 4to80 | 28.2 | 20 | 35.2 | 34.7 | 57.6 |
| | | true fir | 1 | 18 | 18.0 | 18 | 0.3 | 18.0 | 0.2 |
| | | hemlock | 88 | 6to48 | 17.2 | 14 | 30.0 | 19.7 | 15.9 |
| | | bl maple | 22 | 4to40 | 14.7 | 13 | 7.5 | 16.6 | 2.8 |
| | | spruce | 1 | 5 | 5.0 | 5 | 0.3 | 5.0 | 0.0 |
| | | cascara | 4 | 6to10 | 8.0 | 8 | 1.4 | 8.1 | 0.1 |
| | | v maple | 7 | 4to10 | 5.5 | 5 | 2.4 | 6.1 | 0.1 |
| | | willow | 1 | 6 | 6.0 | 6 | 0.3 | 6.0 | 0.0 |

| Twp-Range | 13-4W | species | # WTs | Diam Range | av Diam | Median Diam | rel Freq% | QMD | rel DOM% |
|---|---|---|---|---|---|---|---|---|---|
| # all WTs | 283 | alder | 26 | 5to27 | 9.3 | 9 | 9.2 | 9.9 | 1.6 |
| all avDiam | 20.1 | ash | 3 | 4to24 | 11.3 | 6 | 1.1 | 14.5 | 0.4 |
| ttl QMD | 24.1 | redcedar | 56 | 6to70 | 27.0 | 24 | 19.8 | 30.5 | 31.7 |
| SrvyYr | 1855 | cherry | 7 | 4to10 | 6.3 | 6 | 2.5 | 6.6 | 0.2 |
| | | cottonwood | 8 | 7to48 | 25.1 | 25 | 2.8 | 28.2 | 3.9 |
| | | dogwood | 3 | 8to18 | 12.0 | 10 | 1.1 | 12.8 | 0.3 |
| | | Douglas-fir | 161 | 4to65 | 21.1 | 18 | 56.9 | 24.8 | 60.2 |
| | | bl maple | 17 | 4to30 | 11.5 | 12 | 6.0 | 12.9 | 1.7 |
| | | willow | 2 | 8to10 | 9.0 | 9 | 0.7 | 9.1 | 0.1 |

| Twp-Range | 13-3W | species | # WTs | Diam Range | av Diam | Median Diam | rel Freq% | QMD | rel DOM% |
|---|---|---|---|---|---|---|---|---|---|
| # all WTs | 283 | alder | 13 | 3to14 | 5.5 | 4 | 4.6 | 6.3 | 0.2 |
| all avDiam | 23.1 | ash | 16 | 4to24 | 12.9 | 13 | 5.7 | 14.5 | 1.6 |
| ttl QMD | 27.4 | redcedar | 45 | 6to60 | 23.6 | 20 | 15.9 | 26.7 | 15.1 |
| SrvyYr | 1857 | cherry | 1 | 3 | 3.0 | 3 | 0.4 | 3.0 | 0.0 |
| | | crabapple | 1 | 10 | 10.0 | 10 | 0.4 | 10.0 | 0.0 |
| | | cottonwood | 3 | 5to8 | 6.3 | 6 | 1.1 | 6.5 | 0.1 |
| | | dogwood | 1 | 6 | 6.0 | 6 | 0.4 | 6.0 | 0.0 |
| | | Douglas-fir | 171 | 5to70 | 28.6 | 29 | 60.4 | 31.8 | 81.6 |
| | | hemlock | 2 | 12to20 | 16.0 | 16 | 0.7 | 16.5 | 0.3 |
| | | bl maple | 18 | 4to18 | 8.7 | 7.5 | 6.4 | 9.6 | 0.8 |
| | | oak | 2 | 8 | 8.0 | 8 | 0.7 | 8.0 | 0.1 |
| | | willow | 5 | 4to10 | 7.6 | 9 | 1.8 | 8.0 | 0.1 |
| | | yellowwoo | 5 | 4to8 | 6.0 | 6 | 1.8 | 6.3 | 0.1 |

| Twp-Range | 13-2W | species | # WTs | Diam Range | av Diam | Median Diam | rel Freq% | QMD | rel DOM% |
|---|---|---|---|---|---|---|---|---|---|
| # all WTs | 274 | alder | 20 | 4to28 | 9.3 | 7 | 7.3 | 11.0 | 2.4 |
| all avDiam | 16.4 | ash | 16 | 5to16 | 9.6 | 8 | 5.8 | 10.2 | 1.6 |
| ttl QMD | 19.3 | redcedar | 68 | 6to44 | 18.4 | 15.5 | 24.8 | 20.5 | 28.3 |
| SrvyYr | 1855 | cherry | 3 | 5to18 | 11.7 | 12 | 1.1 | 12.8 | 0.5 |
| | | dogwood | 2 | 3to8 | 5.5 | 5.5 | 0.7 | 6.0 | 0.1 |
| | | Douglas-fir | 120 | 5to50 | 20.3 | 19 | 43.8 | 22.7 | 60.7 |
| | | hemlock | 9 | 6to14 | 9.3 | 9 | 3.3 | 9.7 | 0.8 |
| | | bl maple | 13 | 4to36 | 13.5 | 10 | 4.7 | 17.4 | 3.9 |
| | | oak | 9 | 5to24 | 10.2 | 9 | 3.3 | 11.7 | 1.2 |
| | | v maple | 4 | 4to5 | 4.5 | 4.5 | 1.5 | 4.5 | 0.1 |
| | | willow | 10 | 3to14 | 6.1 | 4 | 3.6 | 7.2 | 0.5 |

| Twp-Range | 13-1W | species | # WTs | Diam Range | av Diam | Median Diam | rel Freq% | QMD | rel DOM% |
|---|---|---|---|---|---|---|---|---|---|
| # all WTs | 283 | alder | 23 | 5to24 | 11.5 | 12 | 8.1 | 12.6 | 1.9 |
| all avDiam | 20.1 | ash | 19 | 4to36 | 13.5 | 12 | 6.7 | 15.5 | 2.4 |
| ttl QMD | 26.1 | redcedar | 78 | 6to120 | 29.4 | 24 | 27.6 | 37.3 | 56.4 |
| SrvyYr | 1857 | cherry | 1 | 6 | 6.0 | 6 | 0.4 | 6.0 | 0.0 |
| | | crabapple | 2 | 6to8 | 7.0 | 7 | 0.7 | 7.1 | 0.1 |
| | | cottonwood | 2 | 5to12 | 8.5 | 8.5 | 0.7 | 9.2 | 0.1 |
| | | dogwood | 1 | 8 | 8.0 | 8 | 0.4 | 8.0 | 0.0 |
| | | Douglas-fir | 115 | 6to70 | 19.3 | 16 | 40.6 | 23.4 | 32.6 |
| | | hemlock | 13 | 5to40 | 14.6 | 12 | 4.6 | 17.5 | 2.1 |
| | | bl maple | 22 | 4to36 | 13.5 | 12 | 7.8 | 15.1 | 2.6 |
| | | oak | 4 | 5to36 | 21.3 | 22 | 1.4 | 24.0 | 1.2 |
| | | spruce | 1 | 36 | 36.0 | 36 | 0.4 | 36.0 | 0.7 |
| | | v maple | 1 | 4 | 4.0 | 4 | 0.4 | 4.0 | 0.0 |
| | | willow | 1 | 8 | 8.0 | 8 | 0.4 | 8.0 | 0.0 |

| Twp-Range | 13-1E | species | # VTs | Diam Range | av Diam | Median Diam | rel Freq% | QMD | rel DOM% |
|---|---|---|---|---|---|---|---|---|---|
| # all VTs | 250 | alder | 14 | 4to24 | 11.4 | 10 | 5.6 | 12.6 | 1.4 |
| all avDiam | 20.6 | ash | 12 | 4to18 | 9.8 | 9 | 4.8 | 10.5 | 0.8 |
| ttl QMD | 25.2 | barberry | 1 | 6 | 6.0 | 6 | 0.4 | 6.0 | 0.0 |
| SrvyYr | 1873 | redcedar | 32 | 4to72 | 28.2 | 22 | 12.8 | 34.2 | 23.5 |
| | | cherry | 4 | 6to30 | 12.0 | 6 | 1.6 | 16.0 | 0.6 |
| | | crabapple | 1 | 4 | 4.0 | 4 | 0.4 | 4.0 | 0.0 |
| | | cottonwood | 3 | 7to24 | 17.0 | 20 | 1.2 | 18.5 | 0.6 |
| | | dogwood | 1 | 10 | 10.0 | 10 | 0.4 | 10.0 | 0.1 |
| | | Douglas-fir | 85 | 4to70 | 26.1 | 24 | 34.0 | 30.6 | 50.0 |
| | | true fir | 2 | 12to24 | 18.0 | 18 | 0.8 | 19.0 | 0.5 |
| | | hemlock | 74 | 4to48 | 17.5 | 14 | 29.6 | 20.0 | 18.5 |
| | | bl maple | 18 | 7to40 | 15.8 | 12 | 7.2 | 18.1 | 3.7 |
| | | spruce | 1 | 10 | 10.0 | 10 | 0.4 | 10.0 | 0.1 |
| | | serviceberi | 1 | 5 | 5.0 | 5 | 0.4 | 5.0 | 0.0 |
| | | v maple | 1 | 4 | 4.0 | 4 | 0.4 | 4.0 | 0.0 |

| Twp-Range | 13-2E | species | # VTs | Diam Range | av Diam | Median Diam | rel Freq% | QMD | rel DOM% |
|---|---|---|---|---|---|---|---|---|---|
| # all VTs | 283 | alder | 41 | 2to20 | 10.0 | 10 | 14.5 | 11.0 | 3.2 |
| all avDiam | 17.5 | ash | 3 | 6to16 | 10.0 | 8 | 1.1 | 10.9 | 0.2 |
| ttl QMD | 23.4 | barberry | 4 | 3to6 | 4.8 | 5 | 1.4 | 4.9 | 0.1 |
| SrvyYr | 1873 | redcedar | 5 | 8to40 | 27.6 | 30 | 1.8 | 30.2 | 2.9 |
| | | cherry | 7 | 3to12 | 6.7 | 5 | 2.5 | 7.6 | 0.3 |
| | | crabapple | 2 | 8to20 | 14.0 | 14 | 0.7 | 15.2 | 0.3 |
| | | Douglas-fir | 176 | 4to84 | 20.7 | 13.5 | 62.2 | 26.8 | 81.7 |
| | | hemlock | 28 | 5to70 | 17.1 | 12 | 9.9 | 23.3 | 9.8 |
| | | bl maple | 7 | 6to30 | 15.1 | 12 | 2.5 | 17.4 | 1.4 |
| | | v maple | 7 | 4to7 | 4.7 | 4 | 2.5 | 4.8 | 0.1 |
| | | willow | 3 | 4to6 | 5.7 | 6 | 1.1 | 5.4 | 0.1 |

| Twp-Range | 13-3E | species | # VTs | Diam Range | av Diam | Median Diam | rel Freq% | QMD | rel DOM% |
|---|---|---|---|---|---|---|---|---|---|
| # all VTs | 288 | alder | 50 | 4to24 | 9.2 | 8 | 17.4 | 10.3 | 9.3 |
| all avDiam | 11.3 | redcedar | 11 | 6to20 | 11.9 | 10 | 3.8 | 12.7 | 3.1 |
| ttl QMD | 14.1 | cherry | 4 | 4to6 | 5.0 | 5 | 1.4 | 5.1 | 0.2 |
| SrvyYr | 1890 | Douglas-fir | 166 | 4to72 | 12.8 | 10 | 57.6 | 16.2 | 76.4 |
| | | hemlock | 28 | 4to30 | 11.6 | 9.5 | 9.7 | 13.0 | 8.3 |
| | | larch | 8 | 5to16 | 9.5 | 9 | 2.8 | 10.2 | 1.4 |
| | | bl maple | 16 | 4to12 | 5.3 | 4 | 5.6 | 5.8 | 0.9 |
| | | spruce | 2 | 6to10 | 8.0 | 8 | 0.7 | 8.3 | 0.2 |
| | | hawthorn | 1 | 5 | 5.0 | 5 | 0.3 | 5.0 | 0.0 |
| | | willow | 2 | 4to5 | 4.5 | 4.5 | 0.7 | 4.5 | 0.1 |

| Twp-Range | 13-4E | species | # VTs | Diam Range | av Diam | Median Diam | rel Freq% | QMD | rel DOM% |
|---|---|---|---|---|---|---|---|---|---|
| # all VTs | 291 | alder | 12 | 6to16 | 10.2 | 9 | 4.1 | 10.8 | 1.3 |
| all avDiam | 16.0 | barberry | 2 | 7to8 | 7.5 | 7.5 | 0.7 | 7.5 | 0.1 |
| ttl QMD | 19.2 | redcedar | 13 | 6to40 | 26.5 | 30 | 4.5 | 29.0 | 10.2 |
| SrvyYr | 1891&1910 | Douglas-fir | 98 | 4to60 | 12.8 | 10 | 33.7 | 15.2 | 21.2 |
| | | true fir | 1 | 60 | 60.0 | 60 | 0.3 | 60.0 | 3.4 |
| | | hemlock | 151 | 4to48 | 18.1 | 17 | 51.9 | 20.9 | 61.6 |
| | | larch | 1 | 6 | 6.0 | 6 | 0.3 | 6.0 | 0.0 |
| | | bl maple | 5 | 6to40 | 15.2 | 10 | 1.7 | 19.7 | 1.8 |
| | | spruce | 1 | 10 | 10.0 | 10 | 0.3 | 10.0 | 0.1 |
| | | v maple | 6 | 5to10 | 6.5 | 6 | 2.1 | 6.7 | 0.3 |
| | | yew | 1 | 6 | 6.0 | 6 | 0.3 | 6.0 | 0.0 |

| Twp-Range | 12-11W | species | # VTs | Diam Range | av Diam | Median Diam | rel Freq% | QMD | rel DOM% |
|---|---|---|---|---|---|---|---|---|---|
| # all VTs | 80 | alder | 16 | 4to18 | 7.5 | 6 | 20.0 | 8.5 | 4.9 |
| all avDiam | 12.3 | crabapple | 6 | 4to8 | 5.2 | 4.5 | 7.5 | 5.4 | 0.7 |
| ttl QMD | 17.2 | Douglas-fir | 1 | 20 | 20.0 | 20 | 1.3 | 20.0 | 1.7 |
| SrvyYr | 1858 | hemlock | 13 | 5to30 | 9.9 | 8 | 16.3 | 11.6 | 7.4 |
| | | pine | 26 | 4to42 | 9.8 | 8 | 32.5 | 13.7 | 20.6 |
| | | spruce | 17 | 5to70 | 24.7 | 24 | 21.3 | 29.9 | 64.5 |
| | | willow | 1 | 7 | 7.0 | 7 | 1.3 | 7.0 | 0.2 |

| Twp-Range | 12-10W | species | # VTs | Diam Range | av Diam | Median Diam | rel Freq% | QMD | rel DOM% |
|---|---|---|---|---|---|---|---|---|---|
| # all VTs | 162 | alder | 4 | 5to12 | 9.8 | 11 | 2.5 | 10.2 | 0.8 |
| all avDiam | 14.9 | redcedar | 2 | 13to56 | 34.5 | 34.5 | 1.2 | 39.7 | 5.8 |
| ttl QMD | 18.3 | cherry | 2 | 6to12 | 9.0 | 9 | 1.2 | 9.5 | 0.3 |
| SrvyYr | 1875 | crabapple | 1 | 6 | 6.0 | 6 | 0.6 | 6.0 | 0.1 |
| | | dogwood | 2 | 4to5 | 4.5 | 4.5 | 1.2 | 4.5 | 0.1 |
| | | hemlock | 138 | 4to48 | 14.8 | 12 | 85.2 | 17.4 | 77.6 |
| | | spruce | 13 | 5to72 | 17.7 | 10 | 8.0 | 24.6 | 14.5 |

| Twp-Range | 12-9W | species | # VTs | Diam Range | av Diam | Median Diam | rel Freq% | QMD | rel DOM% |
|---|---|---|---|---|---|---|---|---|---|
| # all VTs | 284 | alder | 2 | 10 | 10.0 | 10 | 0.7 | 10.0 | 0.1 |
| all avDiam | 20.3 | redcedar | 22 | 9to90 | 31.5 | 24 | 7.7 | 38.8 | 19.5 |
| ttl QMD | 24.4 | Douglas-fir | 7 | 8to28 | 14.3 | 12 | 2.5 | 15.8 | 1.0 |
| SrvyYr | 1894 | true fir | 23 | 10to50 | 20.0 | 14 | 8.1 | 23.3 | 7.4 |
| | | hemlock | 214 | 5to60 | 18.5 | 14.5 | 75.4 | 21.4 | 57.9 |
| | | spruce | 16 | 8to60 | 34.4 | 30 | 5.6 | 38.5 | 14.0 |

| Twp-Range | 12-8W | species | # VTs | Diam Range | av Diam | Median Diam | rel Freq% | QMD | rel DOM% |
|---|---|---|---|---|---|---|---|---|---|
| # all VTs | 291 | alder | 7 | 8to24 | 13.7 | 12 | 2.4 | 14.5 | 1.0 |
| all avDiam | 18.5 | redcedar | 12 | 13to62 | 39.8 | 40 | 4.1 | 42.3 | 15.3 |
| ttl QMD | 22.0 | Douglas-fir | 19 | 5to72 | 25.1 | 20 | 6.5 | 31.9 | 13.8 |
| SrvyYr | 1884&96 | grand fir | 45 | 6to52 | 17.2 | 14 | 15.5 | 19.7 | 12.4 |
| | | hemlock | 189 | 4to53 | 17.9 | 14 | 64.9 | 20.3 | 55.4 |
| | | bl maple | 2 | 12to18 | 15.0 | 15 | 0.7 | 15.3 | 0.3 |
| | | spruce | 8 | 6to30 | 14.6 | 12.5 | 2.7 | 16.7 | 1.6 |
| | | v maple | 6 | 3to5 | 3.5 | 3 | 2.1 | 3.6 | 0.1 |
| | | yew | 3 | 3to11 | 6.0 | 4 | 1.0 | 7.0 | 0.1 |

| Twp-Range | 12-7W | species | # VTs | Diam Range | av Diam | Median Diam | rel Freq% | QMD | rel DOM% |
|---|---|---|---|---|---|---|---|---|---|
| # all VTs | 297 | alder | 9 | 10to24 | 16.3 | 15 | 3.0 | 17.0 | 1.8 |
| all avDiam | 18.1 | barberry | 2 | 4to5 | 4.5 | 4.5 | 0.7 | 4.5 | 0.0 |
| ttl QMD | 22.0 | redcedar | 34 | 8to60 | 31.1 | 36 | 11.4 | 33.6 | 26.7 |
| SrvyYr | 1882 | crabapple | 3 | 4to6 | 4.7 | 4 | 1.0 | 4.8 | 0.0 |
| | | Douglas-fir | 15 | 10to60 | 33.6 | 36 | 5.1 | 36.4 | 13.9 |
| | | grand fir | 32 | 5to48 | 20.5 | 19 | 10.8 | 25.1 | 14.1 |
| | | hemlock | 158 | 4to48 | 15.9 | 14 | 53.2 | 18.3 | 36.7 |
| | | bl maple | 10 | 4to36 | 12.6 | 10.5 | 3.4 | 15.5 | 1.7 |
| | | spruce | 10 | 8to48 | 20.2 | 17 | 3.4 | 23.2 | 3.8 |
| | | v maple | 20 | 3to36 | 6.4 | 4 | 6.7 | 9.4 | 1.2 |
| | | willow | 3 | 5to7 | 6.3 | 7 | 1.0 | 6.4 | 0.1 |
| | | yew | 1 | 10 | 10.0 | 10 | 0.3 | 10.0 | 0.1 |

| Twp-Range | 12-6W | species | # VTs | Diam Range | av Diam | Median Diam | rel Freq% | QMD | rel DOM% |
|---|---|---|---|---|---|---|---|---|---|
| # all VTs | 288 | alder | 5 | 6to15 | 11.8 | 14 | 1.7 | 12.3 | 0.4 |
| all avDiam | 19.9 | barberry | 1 | 6 | 6.0 | 6 | 0.3 | 6.0 | 0.0 |
| ttl QMD | 24.3 | redcedar | 23 | 5to70 | 30.5 | 30 | 8.0 | 34.2 | 15.8 |
| SrvyYr | 1891-02 | dogwood | 1 | 5 | 5.0 | 5 | 0.3 | 5.0 | 0.0 |
| | | Douglas-fir | 32 | 6to72 | 32.9 | 33 | 11.1 | 38.7 | 28.2 |
| | | grand fir | 19 | 5to36 | 13.1 | 10 | 6.6 | 15.5 | 2.7 |
| | | hemlock | 183 | 4to60 | 18.1 | 18 | 63.5 | 20.7 | 46.3 |
| | | larch | 3 | 12to30 | 20.0 | 18 | 1.0 | 21.4 | 0.8 |
| | | bl maple | 2 | 16to24 | 20.0 | 20 | 0.7 | 20.4 | 0.5 |
| | | spruce | 6 | 6to72 | 30.0 | 26 | 2.1 | 37.6 | 5.0 |
| | | v maple | 13 | 4to8 | 5.2 | 5 | 4.5 | 5.4 | 0.2 |

| Twp-Range | 12-5W | species | # VTs | Diam Range | av Diam | Median Diam | rel Freq% | QMD | rel DOM% |
|---|---|---|---|---|---|---|---|---|---|
| # all VTs | 283 | alder | 6 | 3to28 | 18.0 | 21 | 2.1 | 20.0 | 1.4 |
| all avDiam | 20.5 | redcedar | 28 | 6to60 | 25.3 | 24 | 9.9 | 28.2 | 12.8 |
| ttl QMD | 24.8 | crabapple | 1 | 6 | 6.0 | 6 | 0.4 | 6.0 | 0.0 |
| SrvyYr | 1891 | Douglas-fir | 85 | 4to72 | 29.1 | 30 | 30.0 | 33.5 | 54.7 |
| | | grand fir | 5 | 4to72 | 8.6 | 7 | 1.8 | 9.6 | 0.3 |
| | | hemlock | 137 | 4to54 | 16.3 | 12 | 48.4 | 19.2 | 29.1 |
| | | larch | 1 | 15 | 15.0 | 15 | 0.4 | 15.0 | 0.1 |
| | | bl maple | 12 | 6to27 | 12.7 | 11 | 4.2 | 13.9 | 1.3 |
| | | pine | 1 | 18 | 18.0 | 18 | 0.4 | 18.0 | 0.2 |
| | | vine maple | 5 | 3to6 | 4.4 | 5 | 1.8 | 4.6 | 0.1 |
| | | yew | 2 | 5 | 5.0 | 5 | 0.7 | 5.0 | 0.0 |

| Twp-Range | 12-4W | species | # VTs | Diam Range | av Diam | Median Diam | rel Freq% | QMD | rel DOM% |
|---|---|---|---|---|---|---|---|---|---|
| # all VTs | 269 | alder | 8 | 6to16 | 11.6 | 11 | 3.0 | 12.0 | 0.6 |
| all avDiam | 21.7 | ash | 4 | 3to24 | 11.3 | 9 | 1.5 | 13.7 | 0.4 |
| ttl QMD | 26.3 | barberry | 1 | 8 | 8.0 | 8 | 0.4 | 8.0 | 0.0 |
| SrvyYr | 1874 | redcedar | 51 | 5to90 | 21.9 | 20 | 19.0 | 26.5 | 19.2 |
| | | cherry | 2 | 5 | 5.0 | 5 | 0.7 | 5.0 | 0.0 |
| | | crabapple | 1 | 9 | 9.0 | 9 | 0.4 | 9.0 | 0.0 |
| | | dogwood | 1 | 6 | 6.0 | 6 | 0.4 | 6.0 | 0.0 |
| | | Douglas-fir | 109 | 3to96 | 28.8 | 28 | 40.5 | 33.3 | 64.8 |
| | | hemlock | 67 | 6to36 | 17.1 | 14 | 24.9 | 19.0 | 12.9 |
| | | bl maple | 17 | 2to24 | 11.8 | 12 | 6.3 | 12.7 | 1.5 |
| | | pine | 1 | 18 | 18.0 | 18 | 0.4 | 18.0 | 0.2 |
| | | spruce | 1 | 20 | 20.0 | 20 | 0.4 | 20.0 | 0.2 |
| | | v maple | 3 | 4to7 | 5.7 | 6 | 1.1 | 5.8 | 0.1 |
| | | willow | 2 | 4 | 4.0 | 4 | 0.7 | 4.0 | 0.0 |
| | | yew | 1 | 8 | 8.0 | 8 | 0.4 | 8.0 | 0.0 |

| Twp-Range | 12-3W | species | # VTs | Diam Range | av Diam | Median Diam | rel Freq% | QMD | rel DOM% |
|---|---|---|---|---|---|---|---|---|---|
| # all VTs | 283 | alder | 12 | 6to24 | 12.1 | 10 | 4.2 | 13.4 | 1.3 |
| all avDiam | 18.4 | ash | 5 | 3to14 | 8.4 | 8 | 1.8 | 9.1 | 0.3 |
| ttl QMD | 23.7 | barberry | 4 | 6to10 | 7.0 | 6 | 1.4 | 7.2 | 0.1 |
| SrvyYr | 1874 | redcedar | 11 | 7to66 | 33.0 | 36 | 3.9 | 36.9 | 9.4 |
| | | cherry | 13 | 5to36 | 10.2 | 6 | 4.6 | 14.1 | 1.6 |
| | | dogwood | 8 | 4to10 | 6.9 | 6 | 2.8 | 7.3 | 0.3 |
| | | Douglas-fir | 162 | 4to72 | 23.7 | 20 | 57.2 | 28.3 | 81.6 |
| | | hemlock | 13 | 6to48 | 15.2 | 12 | 4.6 | 18.9 | 2.9 |
| | | hazel | 1 | 4 | 4.0 | 4 | 0.4 | 4.0 | 0.0 |
| | | bl maple | 39 | 3to24 | 7.8 | 6 | 13.8 | 9.0 | 2.0 |
| | | spruce | 1 | 20 | 20.0 | 20 | 0.4 | 20.0 | 0.3 |
| | | serviceberi | 1 | 4 | 4.0 | 4 | 0.4 | 4.0 | 0.0 |
| | | v maple | 7 | 4to72 | 5.7 | 6 | 2.5 | 5.8 | 0.1 |
| | | willow | 6 | 4to8 | 5.5 | 5.5 | 2.1 | 5.7 | 0.1 |

| Twp-Range | 12-2W | species | # WTs | Diam Range | av Diam | Median Diam | rel Freq% | QMD | rel DOM% |
|---|---|---|---|---|---|---|---|---|---|
| # all WTs | 293 | alder | 10 | 6to14 | 9.5 | 8.5 | 3.4 | 10.0 | 0.6 |
| all avDiam | 20.0 | ash | 18 | 3to24 | 13.2 | 14 | 6.1 | 14.1 | 2.2 |
| ttl QMD | 23.6 | ctwd/asper | 1 | 3 | 3.0 | 3 | 0.3 | 3.0 | 0.0 |
| SrvyYr | 1853 | redcedar | 36 | 5to72 | 21.8 | 18 | 12.3 | 26.4 | 15.3 |
| | | cherry | 2 | 4to12 | 8.0 | 8 | 0.7 | 8.9 | 0.1 |
| | | crabapple | 1 | 12 | 12.0 | 12 | 0.3 | 12.0 | 0.1 |
| | | dogwood | 2 | 7to12 | 9.5 | 9.5 | 0.7 | 9.8 | 0.1 |
| | | Douglas-fir | 194 | 5to80 | 22.3 | 18 | 66.2 | 25.7 | 78.1 |
| | | hemlock | 6 | 8to24 | 16.3 | 17 | 2.0 | 17.0 | 1.1 |
| | | bl maple | 12 | 4to20 | 11.8 | 13 | 4.1 | 13.0 | 1.2 |
| | | oak | 7 | 5to24 | 15.1 | 16 | 2.4 | 16.2 | 1.1 |
| | | cascara | 2 | 6to8 | 7.0 | 7 | 0.7 | 7.1 | 0.1 |
| | | v maple | 2 | 6 | 6.0 | 6 | 0.7 | 6.0 | 0.0 |

| Twp-Range | 12-1W | species | # WTs | Diam Range | av Diam | Median Diam | rel Freq% | QMD | rel DOM% |
|---|---|---|---|---|---|---|---|---|---|
| # all WTs | 265 | alder | 5 | 6to15 | 8.6 | 6 | 1.9 | 9.3 | 0.5 |
| all avDiam | 15.3 | ash | 23 | 3to30 | 9.9 | 10 | 8.7 | 11.3 | 3.4 |
| ttl QMD | 18.0 | ctwd/asper | 9 | 3to24 | 11.8 | 10 | 3.4 | 13.8 | 2.0 |
| SrvyYr | 1853 | redcedar | 18 | 8to50 | 17.0 | 14.5 | 6.8 | 19.6 | 8.0 |
| | | cherry | 6 | 8to10 | 9.3 | 10 | 2.3 | 9.4 | 0.6 |
| | | crabapple | 2 | 6 | 6.0 | 6 | 0.8 | 6.0 | 0.1 |
| | | Douglas-fir | 143 | 5to50 | 19.1 | 16 | 54.0 | 21.5 | 76.7 |
| | | hemlock | 7 | 3to18 | 11.9 | 12 | 2.6 | 12.7 | 1.3 |
| | | bl maple | 13 | 5to18 | 9.3 | 8 | 4.9 | 10.0 | 1.5 |
| | | oak | 7 | 4to24 | 12.6 | 10 | 2.6 | 17.6 | 2.5 |
| | | pine | 2 | 4to10 | 7.0 | 7 | 0.8 | 7.6 | 0.1 |
| | | spruce | 16 | 2to16 | 10.1 | 11 | 6.0 | 10.8 | 2.2 |
| | | cascara | 1 | 10 | 10.0 | 10 | 0.4 | 10.0 | 0.1 |
| | | v maple | 1 | 5 | 5.0 | 5 | 0.4 | 5.0 | 0.0 |
| | | willow | 12 | 3to12 | 5.2 | 6 | 4.5 | 7.5 | 0.8 |

| Twp-Range | 12-1E | species | # WTs | Diam Range | av Diam | Median Diam | rel Freq% | QMD | rel DOM% |
|---|---|---|---|---|---|---|---|---|---|
| # all WTs | 287 | alder | 29 | 6to24 | 11.6 | 12 | 10.1 | 12.5 | 4.0 |
| all avDiam | 16.5 | ash | 16 | 3to30 | 11.8 | 11 | 5.6 | 13.5 | 2.6 |
| ttl QMD | 20.0 | barberry | 1 | 7 | 7.0 | 7 | 0.3 | 7.0 | 0.0 |
| SrvyYr | 1855 | redcedar | 23 | 6to40 | 19.7 | 18 | 8.0 | 22.3 | 10.0 |
| | | cherry | 6 | 6to12 | 8.3 | 8 | 2.1 | 8.6 | 0.4 |
| | | cottonwood | 3 | 13to24 | 20.3 | 24 | 1.0 | 21.0 | 1.2 |
| | | Douglas-fir | 130 | 5to72 | 21.5 | 20 | 45.3 | 25.0 | 70.8 |
| | | hemlock | 16 | 8to36 | 16.9 | 13.5 | 5.6 | 19.2 | 5.1 |
| | | bl maple | 31 | 4to24 | 11.8 | 10 | 10.8 | 12.8 | 4.4 |
| | | oak | 2 | 6to12 | 9.0 | 9 | 0.7 | 9.5 | 0.2 |
| | | pigeonwood | 3 | 5to24 | 13.0 | 10 | 1.0 | 15.3 | 0.6 |
| | | v maple | 26 | 3to12 | 5.6 | 6 | 9.1 | 5.8 | 0.8 |
| | | willow | 1 | 8 | 8.0 | 8 | 0.3 | 8.0 | 0.1 |

| Twp-Range | 12-2E | species | # WTs | Diam Range | av Diam | Median Diam | rel Freq% | QMD | rel DOM% |
|---|---|---|---|---|---|---|---|---|---|
| # all WTs | 261 | alder | 44 | 6to35 | 12.6 | 12 | 16.9 | 13.7 | 6.6 |
| all avDiam | 17.4 | ash | 13 | 6to29 | 11.9 | 12 | 5.0 | 13.3 | 1.8 |
| ttl QMD | 21.9 | barberry | 4 | 8to12 | 10.0 | 10 | 1.5 | 10.2 | 0.3 |
| SrvyYr | 1872 | redcedar | 18 | 6to60 | 27.7 | 21 | 6.9 | 32.9 | 15.5 |
| | | cherry | 2 | 6to10 | 8.0 | 8 | 0.8 | 8.3 | 0.1 |
| | | crabapple | 3 | 5to12 | 7.7 | 6 | 1.1 | 8.3 | 0.2 |
| | | cottonwood | 2 | 12to20 | 16.0 | 16 | 0.8 | 16.5 | 0.4 |
| | | Douglas-fir | 106 | 4to72 | 22.0 | 16 | 40.6 | 27.4 | 63.2 |
| | | hemlock | 30 | 6to35 | 15.0 | 12.5 | 11.5 | 16.5 | 6.5 |
| | | bl maple | 33 | 5to30 | 12.5 | 12 | 12.6 | 14.0 | 5.1 |
| | | v maple | 4 | 5to6 | 5.8 | 6 | 1.5 | 6.5 | 0.1 |
| | | willow | 2 | 6 | 6.0 | 6 | 0.8 | 6.0 | 0.1 |

| Twp-Range | 12-3E | species | # WTs | Diam Range | av Diam | Median Diam | rel Freq% | QMD | rel DOM% |
|---|---|---|---|---|---|---|---|---|---|
| # all WTs | 276 | alder | 63 | 3to24 | 10.4 | 10 | 22.8 | 11.2 | 13.9 |
| all avDiam | 11.5 | ash | 5 | 6to10 | 8.0 | 8 | 1.8 | 8.2 | 0.6 |
| ttl QMD | 14.4 | barberry | 12 | 4to10 | 7.2 | 7 | 4.3 | 7.4 | 1.1 |
| SrvyYr | 1873 | redcedar | 5 | 6to48 | 19.6 | 12 | 1.8 | 25.0 | 5.5 |
| | | cherry | 2 | 8to10 | 9.0 | 9 | 0.7 | 9.1 | 0.3 |
| | | crabapple | 1 | 6 | 6.0 | 6 | 0.4 | 6.0 | 0.1 |
| | | cottonwood | 2 | 4to10 | 7.0 | 7 | 0.7 | 7.6 | 0.2 |
| | | Douglas-fir | 126 | 4to50 | 13.9 | 10 | 45.7 | 17.7 | 68.7 |
| | | hemlock | 31 | 4to20 | 10.9 | 10 | 11.2 | 11.8 | 7.5 |
| | | bl maple | 16 | 5to11 | 7.2 | 6 | 5.8 | 7.4 | 1.5 |
| | | v maple | 10 | 4to6 | 4.9 | 5 | 3.6 | 5.0 | 0.4 |
| | | willow | 3 | 4to8 | 5.3 | 4 | 1.1 | 5.7 | 0.2 |

| Twp-Range | 12-4E | species | # WTs | Diam Range | av Diam | Median Diam | rel Freq% | QMD | rel DOM% |
|---|---|---|---|---|---|---|---|---|---|
| # all WTs | 292 | alder | 50 | 4to24 | 11.2 | 10 | 17.1 | 12.3 | 7.9 |
| all avDiam | 14.3 | ash | 1 | 5 | 5.0 | 5 | 0.3 | 5.0 | 0.0 |
| ttl QMD | 18.1 | barberry | 6 | 6to10 | 7.5 | 7 | 2.1 | 7.6 | 0.4 |
| SrvyYr | 1880&02 | redcedar | 15 | 5to60 | 24.7 | 18 | 5.1 | 29.3 | 13.3 |
| | | cherry | 2 | 7to20 | 13.3 | 13.5 | 0.7 | 15.0 | 0.5 |
| | | crabapple | 4 | 6to40 | 20.0 | 17 | 1.4 | 23.5 | 2.3 |
| | | cottonwood | 6 | 4to40 | 14.2 | 10 | 2.1 | 18.8 | 2.2 |
| | | dogwood | 1 | 6 | 6.0 | 6 | 0.3 | 6.0 | 0.0 |
| | | Douglas-fir | 115 | 5to84 | 15.1 | 10 | 39.4 | 20.1 | 48.3 |
| | | hemlock | 54 | 5to48 | 15.0 | 12 | 18.5 | 17.3 | 16.7 |
| | | hazel | 1 | 5 | 5.0 | 5 | 0.3 | 5.0 | 0.0 |
| | | bl maple | 24 | 5to40 | 13.7 | 12 | 8.2 | 15.8 | 6.3 |
| | | spruce | 2 | 4to5 | 4.5 | 4.5 | 0.7 | 4.5 | 0.0 |
| | | v maple | 9 | 6to30 | 11.3 | 7 | 3.1 | 14.2 | 1.9 |
| | | willow | 2 | 6 | 6.0 | 6 | 0.7 | 6.0 | 0.1 |

| Twp-Range | 11-11W | species | # WTs | Diam Range | av Diam | Median Diam | rel Freq% | QMD | rel DOM% |
|---|---|---|---|---|---|---|---|---|---|
| # all WTs | 84 | alder | 23 | 4to10 | 6.7 | 6 | 27.4 | 6.9 | 6.3 |
| all avDiam | 9.9 | crabapple | 10 | 4to10 | 5.7 | 5 | 11.9 | 5.9 | 2.0 |
| ttl QMD | 14.4 | dogwood | 1 | 5 | 5.0 | 5 | 1.2 | 5.0 | 0.1 |
| SrvyYr | 1858 | hemlock | 21 | 4to40 | 14.1 | 10 | 25.0 | 17.0 | 35.0 |
| | | pine | 10 | 4to9 | 5.7 | 4.5 | 11.9 | 6.1 | 2.1 |
| | | spruce | 11 | 5to70 | 20.8 | 12 | 13.1 | 29.0 | 53.4 |
| | | willow | 8 | 4to5 | 4.5 | 4.5 | 9.5 | 4.5 | 0.9 |

| Twp-Range | 11-10W | species | # WTs | Diam Range | av Diam | Median Diam | rel Freq% | QMD | rel DOM% |
|---|---|---|---|---|---|---|---|---|---|
| # all WTs | 206 | alder | 8 | 4to14 | 9.4 | 9.5 | 3.9 | 9.9 | 0.9 |
| all avDiam | 17.0 | barberry | 1 | 5 | 5.0 | 5 | 0.5 | 5.0 | 0.0 |
| ttl QMD | 21.0 | redcedar | 2 | 40to84 | 62.0 | 62 | 1.0 | 65.8 | 9.5 |
| SrvyYr | 1873 | cherry | 2 | 5to18 | 11.5 | 11.5 | 1.0 | 13.2 | 0.4 |
| | | crabapple | 3 | 6to16 | 10.7 | 10 | 1.5 | 11.4 | 0.4 |
| | | dogwood | 4 | 4to9 | 6.0 | 5.5 | 1.9 | 6.3 | 0.2 |
| | | hemlock | 172 | 4to60 | 16.9 | 14 | 83.5 | 20.2 | 77.2 |
| | | spruce | 14 | 5to60 | 22.1 | 22 | 6.8 | 27.2 | 11.4 |

| Twp-Range | 11-9W | species | # WTs | Diam Range | av Diam | Median Diam | rel Freq% | QMD | rel DOM% |
|---|---|---|---|---|---|---|---|---|---|
| # all WTs | 289 | alder | 4 | 10to15 | 11.8 | 11 | 1.4 | 12.0 | 0.6 |
| all avDiam | 15.3 | barberry | 1 | 5 | 5.0 | 5 | 0.3 | 5.0 | 0.0 |
| ttl QMD | 18.6 | redcedar | 6 | 11to24 | 15.5 | 14 | 2.1 | 16.1 | 1.6 |
| SrvyYr | 1876&91 | crabapple | 1 | 6 | 6.0 | 6 | 0.3 | 6.0 | 0.0 |
| | | Douglas-fir | 7 | 5to50 | 23.1 | 30 | 2.4 | 28.5 | 5.7 |
| | | hemlock | 232 | 4to46 | 15.0 | 12 | 80.3 | 17.4 | 70.4 |
| | | larch | 18 | 6to53 | 17.5 | 11 | 6.2 | 22.7 | 9.3 |
| | | spruce | 12 | 7to72 | 17.9 | 14 | 4.2 | 31.6 | 12.0 |
| | | v maple | 8 | 4to6 | 4.6 | 4.5 | 2.8 | 4.7 | 0.2 |

| Twp-Range | 11-8W | species | # WTs | Diam Range | av Diam | Median Diam | rel Freq% | QMD | rel DOM% |
|---|---|---|---|---|---|---|---|---|---|
| # all WTs | 284 | alder | 8 | 7to24 | 15.1 | 14 | 2.8 | 16.5 | 1.8 |
| all avDiam | 15.2 | barberry | 8 | 5to10 | 6.3 | 6 | 2.8 | 6.4 | 0.3 |
| ttl QMD | 20.3 | redcedar | 12 | 6to60 | 36.3 | 40 | 4.2 | 39.7 | 16.1 |
| SrvyYr | 1884&94 | Douglas-fir | 10 | 28to70 | 50.1 | 55 | 3.5 | 52.3 | 23.2 |
| | | hemlock | 192 | 3to48 | 13.9 | 12 | 67.6 | 16.1 | 42.3 |
| | | pigeonwood | 1 | 5 | 5.0 | 5 | 0.4 | 5.0 | 0.0 |
| | | spruce | 11 | 5to72 | 29.9 | 10 | 3.9 | 40.7 | 15.5 |
| | | v maple | 42 | 3to7 | 4.5 | 4 | 14.8 | 4.7 | 0.8 |

| Twp-Range | 11-7W | species | # WTs | Diam Range | av Diam | Median Diam | rel Freq% | QMD | rel DOM% |
|---|---|---|---|---|---|---|---|---|---|
| # all WTs | 286 | alder | 15 | 6to30 | 13.4 | 10 | 5.2 | 15.2 | 2.2 |
| all avDiam | 19.5 | redcedar | 29 | 5to80 | 37.1 | 40 | 10.1 | 40.9 | 31.5 |
| ttl QMD | 23.2 | Douglas-fir | 5 | 8to60 | 32.0 | 30 | 1.7 | 38.0 | 4.7 |
| SrvyYr | 1882-02 | true fir | 27 | 5to48 | 20.3 | 15 | 9.4 | 23.7 | 9.8 |
| | | hemlock | 198 | 6to48 | 17.1 | 15 | 69.2 | 19.1 | 46.7 |
| | | spruce | 3 | 48yo50 | 49.3 | 50 | 1.0 | 49.3 | 4.7 |
| | | cascara | 1 | 10 | 10.0 | 10 | 0.3 | 10.0 | 0.1 |
| | | v maple | 8 | 3to9 | 6.0 | 5.5 | 2.8 | 6.3 | 0.2 |

| Twp-Range | 11-6W | species | # WTs | Diam Range | av Diam | Median Diam | rel Freq% | QMD | rel DOM% |
|---|---|---|---|---|---|---|---|---|---|
| # all WTs | 288 | alder | 13 | 6to24 | 13.3 | 12 | 4.5 | 14.3 | 1.4 |
| all avDiam | 21.0 | barberry | 1 | 7 | 7.0 | 7 | 0.3 | 7.0 | 0.0 |
| ttl QMD | 25.4 | redcedar | 17 | 14to60 | 38.0 | 36 | 5.9 | 41.4 | 15.7 |
| SrvyYr | 1896 | Douglas-fir | 7 | 14to70 | 44.0 | 50 | 2.4 | 48.2 | 8.8 |
| | | true fir | 9 | 3to40 | 15.2 | 12 | 3.1 | 19.8 | 1.9 |
| | | hemlock | 185 | 3to60 | 18.3 | 16 | 64.2 | 21.0 | 44.1 |
| | | larch | 31 | 4to50 | 21.1 | 14 | 10.8 | 25.4 | 10.8 |
| | | bl maple | 4 | 24to50 | 36.0 | 35 | 1.4 | 37.3 | 3.0 |
| | | spruce | 19 | 6to90 | 30.5 | 24 | 6.6 | 37.4 | 14.3 |
| | | yew | 2 | 8 | 8.0 | 8 | 0.7 | 8.0 | 0.1 |

| Twp-Range | 11-5W | species | # WTs | Diam Range | av Diam | Median Diam | rel Freq% | QMD | rel DOM% |
|---|---|---|---|---|---|---|---|---|---|
| # all WTs | 288 | alder | 5 | 5to24 | 13.0 | 10 | 1.7 | 14.5 | 0.7 |
| all avDiam | 18.9 | redcedar | 25 | 4to60 | 21.0 | 14 | 8.7 | 26.8 | 11.9 |
| ttl QMD | 22.9 | Douglas-fir | 43 | 7to72 | 23.1 | 15 | 14.9 | 29.1 | 24.1 |
| SrvyYr | 1899 | true fir | 6 | 3to24 | 9.8 | 7 | 2.1 | 12.2 | 0.6 |
| | | hemlock | 180 | 3to60 | 18.2 | 16 | 62.5 | 21.0 | 52.2 |
| | | larch | 22 | 6to50 | 19.1 | 15 | 7.6 | 23.1 | 7.7 |
| | | pine | 1 | 18 | 18.0 | 18 | 0.3 | 18.0 | 0.2 |
| | | spruce | 1 | 60 | 60.0 | 60 | 0.3 | 60.0 | 2.4 |
| | | v maple | 1 | 4 | 4.0 | 4 | 0.3 | 4.0 | 0.0 |
| | | yew | 4 | 6to9 | 7.3 | 7 | 1.4 | 7.3 | 0.1 |

| Twp-Range | 11-4W | species | # WTs | Diam Range | av Diam | Median Diam | rel Freq% | QMD | rel DOM% |
|---|---|---|---|---|---|---|---|---|---|
| # all WTs | 288 | alder | 6 | 10to24 | 15.3 | 14 | 2.1 | 16.0 | 1.4 |
| all avDiam | 17.2 | barberry | 1 | 8 | 8.0 | 8 | 0.3 | 8.0 | 0.1 |
| ttl QMD | 19.8 | redcedar | 16 | 5to50 | 26.9 | 28 | 5.6 | 30.5 | 13.2 |
| SrvyYr | 1892 | dogwood | 1 | 7 | 7.0 | 7 | 0.3 | 7.0 | 0.0 |
| | | Douglas-fir | 25 | 8to60 | 31.8 | 30 | 8.7 | 34.0 | 25.5 |
| | | hemlock | 215 | 2to35 | 15.9 | 14 | 74.7 | 17.4 | 57.7 |
| | | bl maple | 6 | 10to24 | 14.4 | 12 | 2.1 | 15.1 | 1.2 |
| | | pine | 1 | 12 | 12.0 | 12 | 0.3 | 12.0 | 0.1 |
| | | v maple | 13 | 3to8 | 5.9 | 6 | 4.5 | 6.2 | 0.4 |
| | | yew | 4 | 5to12 | 9.3 | 10 | 1.4 | 9.6 | 0.3 |

| | | | | Diam | av | Median | rel | | rel |
|---|---|---|---|---|---|---|---|---|---|
| **Twp-Range** | **11-3W** | **species** | **# WTs** | **Range** | **Diam** | **Diam** | **Freq%** | **QMD** | **DOM%** |
| **# all WTs** | 289 | alder | 17 | 6to20 | 12.3 | 12 | 5.9 | 13.0 | 2.4 |
| **all avDiam** | 17.1 | ash | 2 | 4to15 | 9.5 | 9.5 | 0.7 | 11.0 | 0.2 |
| **ttl QMD** | 20.2 | barberry | 3 | 4to8 | 5.3 | 4 | 1.0 | 5.7 | 0.1 |
| **SrvyYr** | 1873 | redcedar | 23 | 6to40 | 14.8 | 12 | 8.0 | 17.0 | 5.6 |
| | | cherry | 8 | 3to10 | 5.0 | 4.5 | 2.8 | 5.4 | 0.2 |
| | | dogwood | 7 | 5to10 | 7.9 | 7 | 2.4 | 8.1 | 0.4 |
| | | Douglas-fir | 137 | 5to72 | 22.5 | 20 | 47.4 | 25.2 | 73.7 |
| | | hemlock | 47 | 6to36 | 16.9 | 15 | 16.3 | 18.3 | 13.4 |
| | | bl maple | 30 | 4to36 | 9.5 | 6 | 10.4 | 11.7 | 3.5 |
| | | v maple | 10 | 3to14 | 5.8 | 5 | 3.5 | 6.5 | 0.4 |
| | | willow | 5 | 4to14 | 6.6 | 5 | 1.7 | 7.6 | 0.2 |

| | | | | Diam | av | Median | rel | | rel |
|---|---|---|---|---|---|---|---|---|---|
| **Twp-Range** | **11-2W** | **species** | **# WTs** | **Range** | **Diam** | **Diam** | **Freq%** | **QMD** | **DOM%** |
| **# all WTs** | 277 | alder | 20 | 6to20 | 13.8 | 12 | 7.2 | 14.4 | 2.8 |
| **all avDiam** | 20.5 | ash | 3 | 8to18 | 11.3 | 8 | 1.1 | 12.3 | 0.3 |
| **ttl QMD** | 23.2 | redcedar | 51 | 5to60 | 22.1 | 20 | 18.4 | 24.8 | 21.0 |
| **SrvyYr** | 1853 | cherry | 1 | 5 | 5.0 | 5 | 0.4 | 5.0 | 0.0 |
| | | crabapple | 1 | 7 | 7.0 | 7 | 0.4 | 7.0 | 0.0 |
| | | cottonwood | 6 | 8to24 | 18.7 | 23 | 2.2 | 19.9 | 1.6 |
| | | dogwood | 1 | 15 | 15.0 | 15 | 0.4 | 15.0 | 0.2 |
| | | Douglas-fir | 148 | 3to65 | 23.6 | 20 | 53.4 | 26.0 | 66.9 |
| | | hemlock | 15 | 6to36 | 16.9 | 16 | 5.4 | 18.7 | 3.5 |
| | | bl maple | 23 | 3to24 | 12.5 | 15 | 8.3 | 14.2 | 3.1 |
| | | oak | 1 | 20 | 20.0 | 20 | 0.4 | 20.0 | 0.3 |
| | | v maple | 3 | 6to8 | 7.3 | 8 | 1.1 | 7.4 | 0.1 |
| | | willow | 4 | 2to12 | 7.3 | 7.5 | 1.4 | 8.1 | 0.2 |

| | | | | Diam | av | Median | rel | | rel |
|---|---|---|---|---|---|---|---|---|---|
| **Twp-Range** | **11-1W** | **species** | **# WTs** | **Range** | **Diam** | **Diam** | **Freq%** | **QMD** | **DOM%** |
| **# all WTs** | 252 | alder | 22 | 8to30 | 12.7 | 12 | 8.7 | 13.6 | 5.2 |
| **all avDiam** | 14.8 | ash | 30 | 2to16 | 8.9 | 8 | 11.9 | 9.6 | 3.6 |
| **ttl QMD** | 17.5 | redcedar | 27 | 4to48 | 20.1 | 12 | 10.7 | 20.9 | 15.3 |
| **SrvyYr** | 1853 | cherry | 1 | 12 | 12.0 | 12 | 0.4 | 12.0 | 0.2 |
| | | cottonwood | 8 | 7to36 | 20.1 | 20 | 3.2 | 23.2 | 5.6 |
| | | Douglas-fir | 118 | 3to60 | 16.6 | 14.5 | 46.8 | 19.4 | 57.5 |
| | | hemlock | 4 | 8to16 | 11.5 | 11 | 1.6 | 11.9 | 0.7 |
| | | hazel | 1 | 5 | 5.0 | 5 | 0.4 | 5.0 | 0.0 |
| | | bl maple | 28 | 6to30 | 15.0 | 12 | 11.1 | 16.4 | 9.8 |
| | | oak | 4 | 4to24 | 15.5 | 17 | 1.6 | 17.2 | 1.5 |
| | | cascara | 1 | 7 | 7.0 | 7 | 0.4 | 7.0 | 0.1 |
| | | v maple | 4 | 5to8 | 6.3 | 6 | 1.6 | 6.3 | 0.2 |
| | | willow | 4 | 4to10 | 6.5 | 6 | 1.6 | 7.0 | 0.3 |

| | | | | Diam | av | Median | rel | | rel |
|---|---|---|---|---|---|---|---|---|---|
| **Twp-Range** | **11-1E** | **species** | **# WTs** | **Range** | **Diam** | **Diam** | **Freq%** | **QMD** | **DOM%** |
| **# all WTs** | 286 | alder | 23 | 6to24 | 12.0 | 12 | 8.0 | 12.9 | 2.7 |
| **all avDiam** | 18.9 | ash | 10 | 6to30 | 14.7 | 12 | 3.5 | 15.9 | 1.8 |
| **ttl QMD** | 22.5 | barberry | 5 | 6 | 6.0 | 6 | 1.7 | 6.0 | 0.1 |
| **SrvyYr** | 1875 | redcedar | 77 | 6to72 | 24.3 | 18 | 26.9 | 27.6 | 40.7 |
| | | cherry | 2 | 8 | 8.0 | 8 | 0.7 | 8.0 | 0.1 |
| | | cottonwood | 2 | 18to36 | 27.0 | 27 | 0.7 | 28.5 | 1.1 |
| | | dogwood | 1 | 12 | 12.0 | 12 | 0.3 | 12.0 | 0.1 |
| | | Douglas-fir | 102 | 6to72 | 21.8 | 19 | 35.7 | 25.5 | 45.9 |
| | | hemlock | 32 | 6to24 | 12.5 | 12 | 11.2 | 13.5 | 4.1 |
| | | bl maple | 24 | 6to24 | 12.8 | 13.5 | 8.4 | 13.6 | 3.1 |
| | | oak | 1 | 20 | 20.0 | 20 | 0.3 | 20.0 | 0.3 |
| | | v maple | 6 | 4to7 | 5.5 | 6 | 2.1 | 5.6 | 0.1 |
| | | willow | 1 | 6 | 6.0 | 6 | 0.3 | 6.0 | 0.0 |

| | | | | Diam | av | Median | rel | | rel |
|---|---|---|---|---|---|---|---|---|---|
| **Twp-Range** | **11-2E** | **species** | **# WTs** | **Range** | **Diam** | **Diam** | **Freq%** | **QMD** | **DOM%** |
| **# all WTs** | 287 | alder | 22 | 5to18 | 12.1 | 11 | 7.7 | 12.7 | 1.8 |
| **all avDiam** | 23.2 | barberry | 1 | 12 | 12.0 | 12 | 0.3 | 12.0 | 0.1 |
| **ttl QMD** | 26.3 | recedar | 71 | 9to60 | 27.0 | 24 | 24.7 | 30.2 | 32.8 |
| **SrvyYr** | 1896 | crabapple | 1 | 12 | 12.0 | 12 | 0.3 | 12.0 | 0.1 |
| | | Douglas-fir | 134 | 7to60 | 25.5 | 24 | 46.7 | 28.0 | 53.2 |
| | | hemlock | 49 | 6to50 | 19.3 | 18 | 17.1 | 21.4 | 11.4 |
| | | bl maple | 6 | 10to20 | 13.8 | 13 | 2.1 | 14.2 | 0.6 |
| | | v maple | 1 | 8 | 8.0 | 8 | 0.3 | 8.0 | 0.0 |
| | | willow | 2 | 4to5 | 4.5 | 4.5 | 0.7 | 4.5 | 0.0 |

| | | | | Diam | av | Median | rel | | rel |
|---|---|---|---|---|---|---|---|---|---|
| **Twp-Range** | **11-3E** | **species** | **# WTs** | **Range** | **Diam** | **Diam** | **Freq%** | **QMD** | **DOM%** |
| **# all WTs** | 284 | alder | 8 | 7to24 | 16.9 | 15 | 2.8 | 17.9 | 0.8 |
| **all avDiam** | 27.8 | barberry | 1 | 7 | 7.0 | 7 | 0.4 | 7.0 | 0.0 |
| **ttl QMD** | 33.1 | redcedar | 39 | 7to72 | 32.4 | 30 | 13.7 | 36.8 | 17.0 |
| **SrvyYr** | 1896 | Douglas-fir | 91 | 6to108 | 32.6 | 24 | 32.0 | 40.7 | 48.4 |
| | | true fir | 8 | 6to84 | 28.5 | 18 | 2.8 | 37.0 | 3.5 |
| | | hemlock | 133 | 6to50 | 24.4 | 24 | 46.8 | 26.4 | 29.8 |
| | | bl maple | 2 | 6to108 | 8.0 | 8 | 0.7 | 8.3 | 0.0 |
| | | spruce | 1 | 28 | 28.0 | 28 | 0.4 | 28.0 | 0.3 |
| | | v maple | 1 | 8 | 8.0 | 8 | 0.4 | 8.0 | 0.0 |

| | | | | Diam | av | Median | rel | | rel |
|---|---|---|---|---|---|---|---|---|---|
| **Twp-Range** | **11-4E** | **species** | **# WTs** | **Range** | **Diam** | **Diam** | **Freq%** | **QMD** | **DOM%** |
| **# all WTs** | 286 | alder | 6 | 5to24 | 9.0 | 6 | 2.1 | 11.2 | 0.7 |
| **all avDiam** | 16.8 | barberry | 3 | 6to11 | 9.0 | 10 | 1.0 | 9.3 | 0.2 |
| **ttl QMD** | 20.2 | redcedar | 18 | 6to60 | 22.3 | 18 | 6.3 | 25.7 | 10.2 |
| **SrvyYr** | 1899 | cottonwood | 2 | 4to5 | 4.5 | 4.5 | 0.7 | 4.5 | 0.0 |
| | | Douglas-fir | 67 | 3to60 | 19.0 | 14 | 23.4 | 24.1 | 33.5 |
| | | true fir | 29 | 6to40 | 16.0 | 16 | 10.1 | 18.2 | 8.3 |
| | | hemlock | 144 | 4to48 | 17.1 | 15 | 50.3 | 19.4 | 46.3 |
| | | bl maple | 5 | 7to12 | 9.4 | 9 | 1.7 | 9.7 | 0.4 |
| | | v maple | 10 | 4to8 | 5.7 | 5.5 | 3.5 | 5.9 | 0.3 |
| | | willow | 1 | 5 | 5.0 | 5 | 0.3 | 5.0 | 0.0 |
| | | yew | 1 | 12 | 12.0 | 12 | 0.3 | 12.0 | 0.1 |

| | | | | Diam | av | Median | rel | | rel |
|---|---|---|---|---|---|---|---|---|---|
| **Twp-Range** | **10-11W** | **species** | **# WTs** | **Range** | **Diam** | **Diam** | **Freq%** | **QMD** | **DOM%** |
| **# all WTs** | 149 | alder | 30 | 4to18 | 7.8 | 6 | 20.1 | 8.6 | 4.2 |
| **all avDiam** | 14.3 | redcedar | 1 | 60 | 60.0 | 60 | 0.7 | 60.0 | 6.8 |
| **ttl QMD** | 18.8 | crabapple | 4 | 5to6 | 5.3 | 5 | 2.7 | 5.3 | 0.2 |
| **SrvyYr** | 1858 | Douglas-fir | 10 | 6to40 | 19.1 | 18 | 6.7 | 21.7 | 9.0 |
| | | hemlock | 44 | 4to48 | 19.6 | 16 | 29.5 | 23.2 | 45.0 |
| | | pine | 4 | 6to10 | 7.5 | 6 | 2.7 | 7.7 | 0.4 |
| | | spruce | 46 | 4to60 | 14.7 | 10 | 30.9 | 19.6 | 33.6 |
| | | willow | 10 | 4to14 | 5.6 | 4.5 | 6.7 | 6.3 | 0.8 |

| | | | | Diam | av | Median | rel | | rel |
|---|---|---|---|---|---|---|---|---|---|
| **Twp-Range** | **10-10W** | **species** | **# WTs** | **Range** | **Diam** | **Diam** | **Freq%** | **QMD** | **DOM%** |
| **# all WTs** | 259 | alder | 9 | 4to20 | 10.6 | 10 | 3.5 | 11.5 | 1.0 |
| **all avDiam** | 17.1 | redcedar | 10 | 5to50 | 20.4 | 15 | 3.9 | 25.2 | 5.5 |
| **ttl QMD** | 21.1 | crabapple | 1 | 6 | 6.0 | 6 | 0.4 | 6.0 | 0.0 |
| **SrvyYr** | 1858 | Douglas-fir | 1 | 15 | 15.0 | 15 | 0.4 | 15.0 | 0.2 |
| | | hemlock | 198 | 4to56 | 17.0 | 15.5 | 76.4 | 20.4 | 71.9 |
| | | spruce | 39 | 4to72 | 18.9 | 14 | 15.1 | 25.0 | 21.2 |
| | | v maple | 1 | 6 | 6.0 | 6 | 0.4 | 6.0 | 0.0 |

| | | | | Diam | av | Median | rel | | rel |
|---|---|---|---|---|---|---|---|---|---|
| **Twp-Range** | **10-9W** | **species** | **# WTs** | **Range** | **Diam** | **Diam** | **Freq%** | **QMD** | **DOM%** |
| **# all WTs** | 279 | alder | 12 | 6to30 | 12.9 | 11.5 | 4.3 | 14.3 | 2.2 |
| **all avDiam** | 15.1 | barberry | 6 | 5to15 | 8.2 | 6.5 | 2.2 | 8.9 | 0.4 |
| **ttl QMD** | 19.8 | redcedar | 4 | 10to60 | 41.3 | 47.5 | 1.4 | 45.4 | 7.6 |
| **SrvyYr** | 1875 | crabapple | 1 | 5 | 5.0 | 5 | 0.4 | 5.0 | 0.0 |
| | | Douglas-fir | 6 | 5to20 | 11.5 | 8 | 2.2 | 13.0 | 0.9 |
| | | hemlock | 183 | 4to48 | 14.7 | 12 | 65.6 | 17.3 | 50.2 |
| | | bl maple | 3 | 7to18 | 10.7 | 10 | 1.1 | 12.6 | 0.4 |
| | | spruce | 27 | 4to80 | 31.6 | 18 | 9.7 | 38.8 | 37.2 |
| | | v maple | 35 | 4to8 | 5.3 | 5 | 12.5 | 5.4 | 1.0 |
| | | willow | 2 | 4to6 | 5.0 | 5 | 0.7 | 5.1 | 0.0 |

| | | | | Diam | av | Median | rel | | rel |
|---|---|---|---|---|---|---|---|---|---|
| **Twp-Range** | **10-8W** | **species** | **# WTs** | **Range** | **Diam** | **Diam** | **Freq%** | **QMD** | **DOM%** |
| **# all WTs** | 276 | alder | 33 | 6to36 | 16.6 | 14 | 12.0 | 18.5 | 8.6 |
| **all avDiam** | 17.0 | barberry | 1 | 8 | 8.0 | 8 | 0.4 | 8.0 | 0.0 |
| **ttl QMD** | 21.8 | redcedar | 11 | 7to70 | 34.5 | 36 | 4.0 | 39.4 | 13.1 |
| **SrvyYr** | 1888 | dogwood | 1 | 4 | 4.0 | 4 | 0.4 | 4.0 | 0.0 |
| | | Douglas-fir | 39 | 6to72 | 18.0 | 14 | 14.1 | 22.8 | 15.4 |
| | | hemlock | 131 | 4to40 | 15.4 | 12 | 47.5 | 17.9 | 31.9 |
| | | bl maplw | 2 | 10to20 | 15.0 | 15 | 0.7 | 15.8 | 0.4 |
| | | pigeonwood | 2 | 4t05 | 4.5 | 4.5 | 0.7 | 4.5 | 0.0 |
| | | spruce | 33 | 4to74 | 26.6 | 20 | 12.0 | 34.5 | 29.9 |
| | | v maple | 19 | 3to15 | 4.8 | 4 | 6.9 | 5.5 | 0.4 |
| | | willow | 4 | 3to5 | 4.3 | 4.5 | 1.4 | 4.3 | 0.1 |

| | | | | Diam | av | Median | rel | | rel |
|---|---|---|---|---|---|---|---|---|---|
| **Twp-Range** | **10-7W** | **species** | **# WTs** | **Range** | **Diam** | **Diam** | **Freq%** | **QMD** | **DOM%** |
| **# all WTs** | 288 | alder | 28 | 3to36 | 25.7 | 13.5 | 9.7 | 18.3 | 5.9 |
| **all avDiam** | 18.7 | redcedar | 7 | 12to70 | 35.1 | 30 | 2.4 | 39.4 | 6.9 |
| **ttl QMD** | 23.4 | cottonwood | 2 | 18to36 | 27.0 | 27 | 0.7 | 28.5 | 1.0 |
| **SrvyYr** | 1882 | elder | 4 | 4to5 | 4.5 | 4.5 | 1.4 | 4.5 | 0.1 |
| | | Douglas-fir | 10 | 10to70 | 50.0 | 65 | 3.5 | 55.8 | 19.8 |
| | | true fir | 1 | 20 | 20.0 | 20 | 0.3 | 20.0 | 0.3 |
| | | hemlock | 173 | 4to50 | 18.4 | 18 | 60.1 | 20.5 | 46.2 |
| | | pigeonwood | 2 | 6to8 | 7.0 | 7 | 0.7 | 7.1 | 0.1 |
| | | spruce | 25 | 5to75 | 30.5 | 30 | 8.7 | 36.0 | 20.5 |
| | | cascara | 9 | 4to10 | 7.2 | 8 | 3.1 | 7.5 | 0.3 |
| | | v maple | 26 | 4to12 | 5.4 | 5 | 9.0 | 5.7 | 0.5 |
| | | willow | 1 | 10 | 10.0 | 10 | 0.3 | 10.0 | 0.1 |

| | | | | Diam | av | Median | rel | | rel |
|---|---|---|---|---|---|---|---|---|---|
| **Twp-Range** | **10-6W** | **species** | **# WTs** | **Range** | **Diam** | **Diam** | **Freq%** | **QMD** | **DOM%** |
| **# all WTs** | 289 | alder | 26 | 4to48 | 16.2 | 12 | 9.0 | 19.4 | 4.3 |
| **all avDiam** | 23.8 | redcedar | 25 | 10to80 | 36.7 | 36 | 8.7 | 40.2 | 17.7 |
| **ttl QMD** | 28.1 | crabapple | 2 | 7to12 | 9.5 | 9.5 | 0.7 | 9.8 | 0.1 |
| **SrvyYr** | 1872 | Douglas-fir | 25 | 10to70 | 35.3 | 30 | 8.7 | 40.9 | 18.2 |
| | | hemlock | 158 | 4to70 | 23.8 | 24 | 54.7 | 26.6 | 48.8 |
| | | bl maple | 11 | 5to24 | 14.6 | 16 | 3.8 | 15.8 | 1.2 |
| | | spruce | 27 | 4to70 | 23.6 | 18 | 9.3 | 28.3 | 9.5 |
| | | cascara | 7 | 6to10 | 7.9 | 8 | 2.4 | 8.0 | 0.2 |
| | | v maple | 8 | 4to6 | 4.6 | 4 | 2.8 | 4.7 | 0.1 |

| Twp-Range | 10-5W | species | # WTs | Diam Range | av Diam | Median Diam | rel Freq% | QMD | rel DOM% |
|---|---|---|---|---|---|---|---|---|---|
| # all WTs | 288 | alder | 14 | 8to48 | 15.8 | 13 | 4.9 | 18.6 | 1.7 |
| all avDiam | 25.8 | redcedar | 13 | 15to84 | 52.7 | 50 | 4.5 | 59.6 | 16.6 |
| ttl QMD | 31.1 | Douglas-fir | 5 | 36to84 | 67.2 | 72 | 1.7 | 69.6 | 8.7 |
| SrvyYr | 1894 | true fir | 5 | 8to20 | 13.6 | 10 | 1.7 | 14.6 | 0.4 |
| | | hemlock | 182 | 6to60 | 23.2 | 24 | 63.2 | 26.2 | 44.8 |
| | | larch | 41 | 10to60 | 27.7 | 24 | 14.2 | 30.6 | 13.8 |
| | | bl maple | 4 | 10to22 | 17.5 | 19 | 1.4 | 18.1 | 0.5 |
| | | spruce | 14 | 10to100 | 44.6 | 44 | 4.9 | 51.9 | 13.5 |
| | | cascara | 3 | 6to12 | 8.7 | 8 | 1.0 | 9.0 | 0.1 |
| | | v maple | 7 | 4to8 | 5.7 | 6 | 2.4 | 5.9 | 0.1 |

| Twp-Range | 11-4W | species | # WTs | Diam Range | av Diam | Median Diam | rel Freq% | QMD | rel DOM% |
|---|---|---|---|---|---|---|---|---|---|
| # all WTs | 288 | alder | 6 | 4to24 | 12.0 | 11 | 2.1 | 13.7 | 0.9 |
| all avDiam | 17.7 | barberry | 1 | 10 | 10.0 | 10 | 0.3 | 10.0 | 0.1 |
| ttl QMD | 20.5 | redcedar | 6 | 14to48 | 31.3 | 27 | 2.1 | 33.8 | 5.7 |
| SrvyYr | 1894 | Douglas-fir | 30 | 20to60 | 35.1 | 36 | 10.4 | 36.4 | 33.0 |
| | | true fir | 7 | 4to24 | 10.0 | 8 | 2.4 | 11.7 | 0.8 |
| | | hemlock | 219 | 4to48 | 15.5 | 14 | 76.0 | 17.1 | 53.3 |
| | | larch | 11 | 7to36 | 20.5 | 20 | 3.8 | 22.5 | 4.6 |
| | | bl maple | 1 | 6 | 6.0 | 6 | 0.3 | 6.0 | 0.0 |
| | | spruce | 1 | 40 | 40.0 | 40 | 0.3 | 40.0 | 1.3 |
| | | v maple | 5 | 3to6 | 4.4 | 4 | 1.7 | 4.5 | 0.1 |
| | | yew | 1 | 16 | 16.0 | 16 | 0.3 | 16.0 | 0.2 |

| Twp-Range | 10-3W | species | # WTs | Diam Range | av Diam | Median Diam | rel Freq% | QMD | rel DOM% |
|---|---|---|---|---|---|---|---|---|---|
| # all WTs | 289 | alder | 10 | 11to20 | 14.9 | 14.5 | 3.5 | 15.2 | 2.0 |
| all avDiam | 16.5 | ash | 1 | 7 | 7.0 | 7 | 0.3 | 7.0 | 0.0 |
| ttl QMD | 19.9 | barberry | 1 | 7 | 7.0 | 7 | 0.3 | 7.0 | 0.0 |
| SrvyYr | 1881 | redcedar | 35 | 6to50 | 22.2 | 20 | 12.1 | 25.4 | 19.7 |
| | | dogwood | 4 | 6to9 | 7.5 | 0.5 | 1.4 | 7.6 | 0.2 |
| | | Douglas-fir | 54 | 6to60 | 22.6 | 20 | 18.7 | 26.8 | 33.8 |
| | | hemlock | 149 | 4to40 | 15.5 | 12 | 51.6 | 18.0 | 42.0 |
| | | bl maple | 9 | 4to20 | 10.1 | 8 | 3.1 | 11.4 | 1.0 |
| | | spruce | 7 | 5to20 | 10.3 | 10 | 2.4 | 11.3 | 0.8 |
| | | v maple | 19 | 4to7 | 5.4 | 5 | 6.6 | 5.5 | 0.5 |

| Twp-Range | 10-2W | species | # WTs | Diam Range | av Diam | Median Diam | rel Freq% | QMD | rel DOM% |
|---|---|---|---|---|---|---|---|---|---|
| # all WTs | 284 | alder | 18 | 4to24 | 12.2 | 12 | 6.3 | 13.3 | 1.7 |
| all avDiam | 21.1 | redcedar | 42 | 4to50 | 20.4 | 18 | 14.8 | 23.8 | 12.5 |
| ttl QMD | 25.9 | cherry | 1 | 5 | 5.0 | 5 | 0.4 | 5.0 | 0.0 |
| SrvyYr | 1858&74 | cottonwood | 2 | 4to6 | 5.0 | 5 | 0.7 | 5.1 | 0.0 |
| | | dogwood | 7 | 4to10 | 8.9 | 8 | 2.5 | 8.5 | 0.3 |
| | | Douglas-fir | 132 | 5to98 | 28.8 | 25.5 | 46.5 | 33.2 | 76.1 |
| | | hemlock | 32 | 6to36 | 14.1 | 12 | 11.3 | 15.7 | 4.1 |
| | | hazel | 1 | 4 | 4.0 | 4 | 0.4 | 4.0 | 0.0 |
| | | bl maple | 30 | 4to30 | 16.0 | 18 | 10.6 | 17.6 | 4.9 |
| | | pine | 1 | 10 | 10.0 | 10 | 0.4 | 10.0 | 0.1 |
| | | cascara | 1 | 8 | 8.0 | 8 | 0.4 | 8.0 | 0.0 |
| | | v maple | 14 | 3to8 | 5.4 | 5 | 4.9 | 5.6 | 0.2 |
| | | willow | 3 | 3to6 | 4.0 | 3 | 1.1 | 4.2 | 0.0 |

| Twp-Range | 10-1W | species | # WTs | Diam Range | av Diam | Median Diam | rel Freq% | QMD | rel DOM% |
|---|---|---|---|---|---|---|---|---|---|
| # all WTs | 280 | alder | 30 | 3to24 | 10.6 | 10 | 10.7 | 11.9 | 3.2 |
| all avDiam | 18.5 | ash | 7 | 3to16 | 9.0 | 8 | 2.5 | 10.1 | 0.5 |
| ttl QMD | 21.7 | redcedar | 61 | 5to48 | 20.3 | 18 | 21.8 | 22.8 | 24.0 |
| SrvyYr | 1858&74 | crabapple | 5 | 5to12 | 7.0 | 5 | 1.8 | 7.5 | 0.2 |
| | | cottonwood | 2 | 5to7 | 6.0 | 6 | 0.7 | 6.1 | 0.1 |
| | | dogwood | 8 | 7to18 | 9.9 | 9 | 2.9 | 10.4 | 0.7 |
| | | Douglas-fir | 123 | 6to60 | 23.5 | 22 | 43.9 | 26.3 | 64.4 |
| | | hemlock | 16 | 5to30 | 14.8 | 15 | 5.7 | 17.0 | 3.5 |
| | | bl maple | 16 | 4to30 | 14.7 | 13.5 | 5.7 | 15.9 | 3.0 |
| | | cascara | 2 | 4to10 | 7.0 | 7 | 0.7 | 7.6 | 0.1 |
| | | v maple | 8 | 4to8 | 5.3 | 5 | 2.9 | 5.4 | 0.2 |
| | | willow | 2 | 3to4 | 3.5 | 3.5 | 0.7 | 3.5 | 0.0 |

| Twp-Range | 10-1E | species | # WTs | Diam Range | av Diam | Median Diam | rel Freq% | QMD | rel DOM% |
|---|---|---|---|---|---|---|---|---|---|
| # all WTs | 285 | alder | 24 | 5to18 | 9.1 | 8 | 8.4 | 9.8 | 1.3 |
| all avDiam | 20.8 | ash | 6 | 2to36 | 15.2 | 13.5 | 2.1 | 18.5 | 1.1 |
| ttl QMD | 25.2 | redcedar | 76 | 6to60 | 23.6 | 20 | 26.7 | 26.9 | 30.5 |
| SrvyYr | 1875 | cherry | 4 | 6to8 | 7.5 | 8 | 1.4 | 7.6 | 0.1 |
| | | crabapple | 1 | 2 | 2.0 | 2 | 0.4 | 2.0 | 0.0 |
| | | cottonwood | 3 | 24to48 | 34.0 | 30 | 1.1 | 35.5 | 2.1 |
| | | Douglas-fir | 144 | 4to85 | 23.3 | 20 | 50.5 | 27.9 | 61.8 |
| | | hemlock | 12 | 6to24 | 14.6 | 13.5 | 4.2 | 15.8 | 1.6 |
| | | bl maple | 12 | 5to36 | 10.6 | 8 | 4.2 | 13.2 | 1.2 |
| | | spruce | 1 | 20 | 20.0 | 20 | 0.4 | 20.0 | 0.2 |
| | | v maple | 1 | 6 | 6.0 | 6 | 0.4 | 6.0 | 0.0 |
| | | willow | 1 | 4 | 4.0 | 4 | 0.4 | 4.0 | 0.0 |

| Twp-Range | 10-2E | species | # WTs | Diam Range | av Diam | Median Diam | rel Freq% | QMD | rel DOM% |
|---|---|---|---|---|---|---|---|---|---|
| # all WTs | 284 | alder | 21 | 5to24 | 10.1 | 10 | 7.4 | 11.1 | 2.1 |
| all avDiam | 15.5 | redcedar | 51 | 5to96 | 26.6 | 20 | 18.0 | 33.5 | 46.3 |
| ttl QMD | 20.9 | cherry | 1 | 6 | 6.0 | 6 | 0.4 | 6.0 | 0.0 |
| SrvyYr | 1891 | cottonwood | 1 | 6 | 6.0 | 6 | 0.4 | 6.0 | 0.0 |
| | | Douglas-fir | 135 | 5to84 | 14.6 | 10 | 47.5 | 19.4 | 41.2 |
| | | hemlock | 60 | 5to36 | 11.4 | 9.5 | 21.1 | 13.1 | 8.3 |
| | | bl maple | 9 | 6to14 | 9.9 | 10 | 3.2 | 10.2 | 0.8 |
| | | spruce | 2 | 24to30 | 27.0 | 27 | 0.7 | 27.2 | 1.2 |
| | | v maple | 3 | 4to6 | 5.0 | 5 | 1.1 | 5.1 | 0.1 |
| | | willow | 1 | 6 | 6.0 | 6 | 0.4 | 6.0 | 0.0 |

| Twp-Range | 10-3E | species | # WTs | Diam Range | av Diam | Median Diam | rel Freq% | QMD | rel DOM% |
|---|---|---|---|---|---|---|---|---|---|
| # all WTs | 288 | alder | 11 | 6to25 | 12.6 | 12 | 3.8 | 13.5 | 1.1 |
| all avDiam | 19.3 | redcedar | 45 | 5to120 | 31.1 | 30 | 15.6 | 37.7 | 36.8 |
| ttl QMD | 24.6 | Douglas-fir | 75 | 6to84 | 20.1 | 14 | 26.0 | 26.8 | 30.9 |
| SrvyYr | 1891 | hemlock | 147 | 5to50 | 16.0 | 14 | 51.0 | 18.6 | 29.2 |
| | | bl maple | 5 | 6to10 | 6.8 | 6 | 1.7 | 7.0 | 0.1 |
| | | spruce | 5 | 12to36 | 23.6 | 26 | 1.7 | 25.4 | 1.8 |

| Twp-Range | 9-10W | species | # WTs | Diam Range | av Diam | Median Diam | rel Freq% | QMD | rel DOM% |
|---|---|---|---|---|---|---|---|---|---|
| # all WTs | 141 | alder | 22 | 4to18 | 8.3 | 7 | 15.6 | 9.1 | 3.4 |
| all avDiam | 15.0 | redcedar | 2 | 10to54 | 32.0 | 32 | 1.4 | 38.8 | 5.7 |
| ttl QMD | 19.4 | crabapple | 1 | 6 | 6.0 | 6 | 0.7 | 6.0 | 0.1 |
| SrvyYr | 1858 | Douglas-fir | 6 | 4to60 | 20.5 | 15.5 | 4.3 | 27.6 | 8.6 |
| | | hemlock | 90 | 4to48 | 14.9 | 10 | 63.8 | 18.7 | 59.1 |
| | | spruce | 19 | 4to50 | 20.5 | 14 | 13.5 | 25.4 | 23.1 |
| | | willow | 1 | 4 | 4.0 | 4 | 0.7 | 4.0 | 0.0 |

| Twp-Range | 9-9W | species | # WTs | Diam Range | av Diam | Median Diam | rel Freq% | QMD | rel DOM% |
|---|---|---|---|---|---|---|---|---|---|
| # all WTs | 66 | alder | 3 | 4to24 | 12.0 | 8 | 4.5 | 14.8 | 3.7 |
| all avDiam | 12.7 | redcedar | 1 | 24 | 24.0 | 24 | 1.5 | 24.0 | 3.2 |
| ttl QMD | 16.4 | Douglas-fir | 4 | 5to24 | 14.3 | 15 | 6.1 | 16.3 | 6.0 |
| SrvyYr | 1858 | hemlock | 50 | 3to48 | 11.5 | 30 | 75.8 | 14.9 | 62.0 |
| | | bl maple | 1 | 4 | 4.0 | 4 | 1.5 | 4.0 | 0.1 |
| | | spruce | 5 | 5to48 | 25.4 | 24 | 7.6 | 29.7 | 24.7 |
| | | v maple | 2 | 5to6 | 5.5 | 5.5 | 3.0 | 5.5 | 0.3 |

| Twp-Range | 9-8W | species | # WTs | Diam Range | av Diam | Median Diam | rel Freq% | QMD | rel DOM% |
|---|---|---|---|---|---|---|---|---|---|
| # all WTs | 74 | alder | 3 | 4to30 | 15.3 | 12 | 4.1 | 18.8 | 2.8 |
| all avDiam | 18.5 | redcedar | 1 | 30 | 30.0 | 30 | 1.4 | 30.0 | 2.4 |
| ttl QMD | 22.6 | Douglas-fir | 4 | 8to23 | 10.8 | 8 | 5.4 | 13.0 | 1.8 |
| SrvyYr | 1872 | hemlock | 45 | 5to45 | 27.9 | 14 | 60.8 | 21.1 | 53.1 |
| | | bl maple | 5 | 4to36 | 11.8 | 6 | 6.8 | 17.0 | 3.8 |
| | | spruce | 14 | 8to50 | 22.3 | 19 | 18.9 | 25.8 | 24.8 |
| | | v maple | 2 | 5to65 | 35.0 | 35 | 2.7 | 46.1 | 11.3 |

| Twp-Range | 9-7W | species | # WTs | Diam Range | av Diam | Median Diam | rel Freq% | QMD | rel DOM% |
|---|---|---|---|---|---|---|---|---|---|
| # all WTs | 89 | alder | 7 | 5to36 | 13.9 | 10 | 7.9 | 17.2 | 12.5 |
| all avDiam | 10.7 | redcedar | 1 | 16 | 16.0 | 16 | 1.1 | 16.0 | 1.6 |
| ttl QMD | 13.6 | dogwood | 5 | 6to20 | 9.2 | 6 | 5.6 | 10.7 | 3.5 |
| SrvyYr | 1877 | elder | 2 | 4 | 4.0 | 4 | 2.2 | 4.0 | 0.2 |
| | | Douglas-fir | 2 | 10to15 | 12.5 | 12.5 | 2.2 | 12.8 | 2.0 |
| | | hemlock | 45 | 4to50 | 12.7 | 10 | 50.6 | 15.4 | 64.7 |
| | | bl maple | 17 | 3to30 | 6.7 | 4 | 19.1 | 9.1 | 8.5 |
| | | spruce | 1 | 30 | 30.0 | 30 | 1.1 | 30.0 | 5.5 |
| | | cascara | 4 | 3to8 | 5.3 | 5 | 4.5 | 5.6 | 0.8 |
| | | v maple | 5 | 5to6 | 5.4 | 5 | 5.6 | 5.4 | 0.9 |

| Twp-Range | 9-6W | species | # WTs | Diam Range | av Diam | Median Diam | rel Freq% | QMD | rel DOM% |
|---|---|---|---|---|---|---|---|---|---|
| # all WTs | 206 | alder | 23 | 5to40 | 14.7 | 12 | 11.2 | 17.1 | 5.9 |
| all avDiam | 18.6 | ash | 1 | 8 | 8.0 | 8 | 0.5 | 8.0 | 0.1 |
| ttl QMD | 23.6 | redcedar | 10 | 7to65 | 17.6 | 10 | 4.9 | 24.6 | 5.3 |
| SrvyYr | 1871 | crabapple | 4 | 4to12 | 6.8 | 5.5 | 1.9 | 7.5 | 0.2 |
| | | cottonwood | 9 | 8to30 | 16.9 | 18 | 4.4 | 18.2 | 2.6 |
| | | Douglas-fir | 4 | 8to24 | 17.5 | 19 | 1.9 | 18.5 | 1.2 |
| | | hemlock | 85 | 4to72 | 22.2 | 18 | 41.3 | 26.5 | 52.4 |
| | | bl maple | 7 | 5to40 | 20.7 | 18 | 3.4 | 24.8 | 3.8 |
| | | spruce | 39 | 4to60 | 20.6 | 15 | 18.9 | 28.7 | 28.1 |
| | | cascara | 3 | 5to12 | 9.0 | 10 | 1.5 | 9.5 | 0.2 |
| | | hawthorn | 1 | 4 | 4.0 | 4 | 0.5 | 4.0 | 0.0 |
| | | v maple | 13 | 4to60 | 4.4 | 4 | 6.3 | 4.5 | 0.2 |
| | | willow | 7 | 3to8 | 4.7 | 4 | 3.4 | 4.9 | 0.1 |

| Twp-Range | 9-5W | species | # WTs | Diam Range | av Diam | Median Diam | rel Freq% | QMD | rel DOM% |
|---|---|---|---|---|---|---|---|---|---|
| # all WTs | 287 | alder | 6 | 4to30 | 12.0 | 10.5 | 2.1 | 14.8 | 0.8 |
| all avDiam | 19.7 | redcedar | 11 | 7to90 | 25.9 | 18 | 3.8 | 34.3 | 7.6 |
| ttl QMD | 24.3 | Douglas-fir | 23 | 14to50 | 27.3 | 20 | 8.0 | 29.6 | 11.8 |
| SrvyYr | 1871 | true fir | 5 | 6to24 | 16.0 | 16 | 1.7 | 17.3 | 0.9 |
| | | hemlock | 194 | 5to60 | 19.0 | 16 | 67.6 | 22.2 | 56.2 |
| | | bl maple | 7 | 3to36 | 17.3 | 10 | 2.4 | 21.9 | 2.0 |
| | | spruce | 17 | 9to96 | 38.8 | 34 | 5.9 | 45.0 | 20.2 |
| | | cascara | 2 | 5to13 | 9.0 | 9 | 0.7 | 9.9 | 0.1 |
| | | v maple | 22 | 3to15 | 5.0 | 4 | 7.7 | 5.5 | 0.4 |

| Twp-Range | 9-4W | species | # WTs | Diam Range | av Diam | Median Diam | rel Freq% | QMD | rel DOM% |
|---|---|---|---|---|---|---|---|---|---|
| # all WTs | 280 | alder | 11 | 9to70 | 19.5 | 15 | 3.9 | 25.4 | 4.7 |
| all avDiam | 19.6 | redcedar | 22 | 5to60 | 29.5 | 18 | 7.9 | 35.2 | 18.1 |
| ttl QMD | 23.2 | Douglas-fir | 92 | 4to72 | 23.0 | 20 | 32.9 | 26.5 | 42.9 |
| SrvyYr | 1879 | true fir | 6 | 6to30 | 12.3 | 10 | 2.1 | 14.8 | 0.9 |
| | | hemlock | 138 | 4to48 | 16.3 | 15 | 49.3 | 18.1 | 30.1 |
| | | bl maple | 8 | 8to48 | 21.4 | 18 | 2.9 | 25.1 | 3.3 |
| | | v maple | 3 | 5 | 5.0 | 5 | 1.1 | 5.0 | 0.0 |

| Twp-Range | 9-3W | species | # WTs | Diam Range | av Diam | Median Diam | rel Freq% | QMD | rel DOM% |
|---|---|---|---|---|---|---|---|---|---|
| # all WTs | 280 | alder | 11 | 9to70 | 19.5 | 15 | 3.9 | 25.4 | 4.7 |
| all avDiam | 19.6 | redcedar | 22 | 5to60 | 29.5 | 18 | 7.9 | 35.2 | 18.1 |
| ttl QMD | 23.2 | Douglas-fir | 92 | 4to72 | 23.0 | 20 | 32.9 | 26.5 | 42.9 |
| SrvyYr | 1879 | true fir | 6 | 6to30 | 12.3 | 10 | 2.1 | 14.8 | 0.9 |
| | | hemlock | 138 | 4to48 | 16.3 | 15 | 49.3 | 18.1 | 30.1 |
| | | bl maple | 8 | 8to48 | 21.4 | 18 | 2.9 | 25.1 | 3.3 |
| | | v maple | 3 | 5 | 5.0 | 5 | 1.1 | 5.0 | 0.0 |

| Twp-Range | 9-2W | species | # WTs | Diam Range | av Diam | Median Diam | rel Freq% | QMD | rel DOM% |
|---|---|---|---|---|---|---|---|---|---|
| # all WTs | 284 | alder | 13 | 4to20 | 11.3 | 10 | 4.6 | 12.4 | 1.1 |
| all avDiam | 20.2 | ash | 5 | 4to24 | 18.4 | 20 | 1.8 | 19.8 | 1.1 |
| ttl QMD | 24.9 | redcedar | 48 | 10to72 | 29.4 | 28 | 16.9 | 32.5 | 28.7 |
| SrvyYr | 1858&79 | cherry | 18 | 3to8 | 4.9 | 4 | 6.3 | 5.3 | 0.3 |
| | | cottonwood | 8 | 5to40 | 14.1 | 10 | 2.8 | 17.8 | 1.4 |
| | | dogwood | 4 | 4to10 | 7.5 | 8 | 1.4 | 7.8 | 0.1 |
| | | elder | 1 | 6 | 6.0 | 6 | 0.4 | 6.0 | 0.0 |
| | | Douglas-fir | 85 | 6to72 | 28.9 | 30 | 29.9 | 32.4 | 50.6 |
| | | hemlock | 38 | 4to40 | 18.1 | 15.5 | 13.4 | 21.0 | 9.5 |
| | | hazel | 2 | 4to6 | 5.0 | 5 | 0.7 | 5.1 | 0.0 |
| | | bl maple | 40 | 4to48 | 14.5 | 11 | 14.1 | 17.1 | 6.6 |
| | | v maple | 19 | 3to10 | 5.5 | 6 | 6.7 | 5.9 | 0.4 |
| | | willow | 3 | 2to6 | 3.7 | 3 | 1.1 | 4.0 | 0.0 |

| Twp-Range | 9-1W | species | # WTs | Diam Range | av Diam | Median Diam | rel Freq% | QMD | rel DOM% |
|---|---|---|---|---|---|---|---|---|---|
| # all WTs | 269 | alder | 8 | 4to15 | 9.3 | 9 | 3.0 | 9.9 | 0.5 |
| all avDiam | 20.3 | ash | 1 | 15 | 15.0 | 15 | 0.4 | 15.0 | 0.1 |
| ttl QMD | 23.8 | redcedar | 60 | 6to60 | 20.5 | 20 | 22.3 | 26.6 | 27.9 |
| SrvyYr | 1858 | cherry | 2 | 5to6 | 5.5 | 5.5 | 0.7 | 5.5 | 0.0 |
| | | cottonwood | 3 | 5to10 | 7.7 | 8 | 1.1 | 7.9 | 0.1 |
| | | dogwood | 1 | 6 | 6.0 | 6 | 0.4 | 6.0 | 0.0 |
| | | Douglas-fir | 87 | 4to72 | 26.7 | 24 | 32.3 | 29.5 | 49.6 |
| | | hemlock | 84 | 5to48 | 15.8 | 14 | 31.2 | 18.0 | 17.9 |
| | | bl maple | 13 | 10to30 | 19.2 | 20 | 4.8 | 20.1 | 3.5 |
| | | v maple | 9 | 4to7 | 5.2 | 5 | 3.3 | 5.3 | 0.2 |
| | | yew | 1 | 15 | 15.0 | 15 | 0.4 | 15.0 | 0.1 |

| Twp-Range | 9-1E | species | # WTs | Diam Range | av Diam | Median Diam | rel Freq% | QMD | rel DOM% |
|---|---|---|---|---|---|---|---|---|---|
| # all WTs | 285 | alder | 5 | 6to10 | 8.4 | 8 | 1.8 | 8.5 | 0.2 |
| all avDiam | 18.0 | ash | 1 | 20 | 20.0 | 20 | 0.4 | 20.0 | 0.3 |
| ttl QMD | 22.6 | redcedar | 58 | 6to46 | 19.1 | 18 | 20.4 | 21.1 | 17.7 |
| SrvyYr | 1872 | cherry | 3 | 4to6 | 4.7 | 4 | 1.1 | 4.8 | 0.0 |
| | | crabapple | 1 | 2 | 2.0 | 2 | 0.4 | 2.0 | 0.0 |
| | | dogwood | 1 | 8 | 8.0 | 8 | 0.4 | 8.0 | 0.0 |
| | | Douglas-fir | 100 | 4to72 | 24.0 | 18 | 35.1 | 30.4 | 63.4 |
| | | grand fir | 2 | 18to20 | 19.0 | 19 | 0.7 | 19.0 | 0.5 |
| | | hemlock | 113 | 5to40 | 13.2 | 10 | 39.6 | 15.1 | 17.8 |
| | | willow | 1 | 6 | 6.0 | 6 | 0.4 | 6.0 | 0.0 |

| Twp-Range | 9-2E | species | # WTs | Diam Range | av Diam | Median Diam | rel Freq% | QMD | rel DOM% |
|---|---|---|---|---|---|---|---|---|---|
| # all WTs | 284 | alder | 2 | 10to20 | 15.0 | 15 | 0.7 | 15.8 | 0.3 |
| all avDiam | 20.3 | redcedar | 60 | 5to144 | 32.0 | 30 | 21.1 | 40.4 | 49.9 |
| ttl QMD | 26.3 | Douglas-fir | 50 | 5to60 | 21.2 | 18 | 17.6 | 25.6 | 16.7 |
| SrvyYr | 1891 | true fir | 1 | 10 | 10.0 | 10 | 0.4 | 10.0 | 0.1 |
| | | hemlock | 170 | 5to84 | 16.0 | 10 | 59.9 | 19.5 | 32.9 |
| | | bl maple | 1 | 16 | 16.0 | 16 | 0.4 | 16.0 | 0.1 |

| Twp-Range | 8-5W | species | # WTs | Diam Range | av Diam | Median Diam | rel Freq% | QMD | rel DOM% |
|---|---|---|---|---|---|---|---|---|---|
| # all WTs | 181 | alder | 11 | 4to18 | 9.1 | 8 | 6.1 | 10.2 | 1.9 |
| all avDiam | 14.4 | redcedar | 5 | 6to40 | 23.6 | 26 | 2.8 | 27.2 | 6.2 |
| ttl QMD | 18.2 | cherry | 1 | 10 | 10.0 | 10 | 0.6 | 10.0 | 0.2 |
| SrvyYr | 1873 | Douglas-fir | 29 | 5to60 | 23.5 | 20 | 16.0 | 28.1 | 38.2 |
| | | true fir | 12 | 4to40 | 9.2 | 6.5 | 6.6 | 13.1 | 3.4 |
| | | hemlock | 110 | 3to48 | 13.6 | 10 | 60.8 | 16.2 | 47.9 |
| | | bl maple | 6 | 4to9 | 6.2 | 5.5 | 3.3 | 6.5 | 0.4 |
| | | madrone | 1 | 4 | 4.0 | 4 | 0.6 | 4.0 | 0.0 |
| | | spruce | 1 | 30 | 30.0 | 30 | 0.6 | 30.0 | 1.5 |
| | | v maple | 3 | 3to6 | 5.0 | 6 | 1.7 | 5.2 | 0.1 |
| | | willow | 2 | 4to10 | 7.0 | 7 | 1.1 | 7.6 | 0.2 |

| Twp-Range | 8-4W | species | # WTs | Diam Range | av Diam | Median Diam | rel Freq% | QMD | rel DOM% |
|---|---|---|---|---|---|---|---|---|---|
| # all WTs | 80 | alder | 4 | 9to15 | 12.0 | 12 | 5.0 | 12.2 | 1.5 |
| all avDiam | 17.7 | redcedar | 7 | 4to60 | 25.7 | 27 | 8.8 | 31.4 | 17.6 |
| ttl QMD | 22.1 | Douglas-fir | 12 | 6to70 | 27.0 | 22 | 15.0 | 33.4 | 34.1 |
| SrvyYr | 1856 | hemlock | 45 | 3to48 | 16.8 | 15 | 56.3 | 19.4 | 43.2 |
| | | bl maple | 4 | 10to20 | 13.8 | 12.5 | 5.0 | 14.4 | 2.1 |
| | | cascara | 1 | 20 | 20.0 | 20 | 1.3 | 20.0 | 1.0 |
| | | v maple | 4 | 4to8 | 6.0 | 6 | 5.0 | 6.2 | 0.4 |
| | | willow | 3 | 4 | 4.0 | 4 | 3.8 | 4.0 | 0.1 |

| Twp-Range | 8-3W | species | # WTs | Diam Range | av Diam | Median Diam | rel Freq% | QMD | rel DOM% |
|---|---|---|---|---|---|---|---|---|---|
| # all WTs | 163 | alder | 9 | 10to18 | 13.8 | 12 | 5.5 | 14.1 | 3.7 |
| all avDiam | 13.2 | ash | 11 | 3to40 | 13.7 | 10 | 6.7 | 16.7 | 6.3 |
| ttl QMD | 17.3 | redcedar | 7 | 12to50 | 24.7 | 24 | 4.3 | 27.8 | 11.0 |
| SrvyYr | 1858 | cherry | 1 | 12 | 12.0 | 12 | 0.6 | 12.0 | 0.3 |
| | | crabapple | 2 | 3to6 | 4.5 | 4.5 | 1.2 | 4.7 | 0.1 |
| | | cottonwood | 4 | 6to12 | 9.5 | 10 | 2.5 | 9.8 | 0.8 |
| | | dogwood | 1 | 15 | 15.0 | 15 | 0.6 | 15.0 | 0.5 |
| | | Douglas-fir | 48 | 6to60 | 21.8 | 18 | 29.4 | 25.8 | 65.6 |
| | | hemlock | 20 | 3to24 | 10.7 | 10 | 12.3 | 11.8 | 5.7 |
| | | hazel | 1 | 3 | 3.0 | 3 | 0.6 | 3.0 | 0.0 |
| | | bl maple | 9 | 4to18 | 9.7 | 10 | 5.5 | 10.6 | 2.1 |
| | | oak | 2 | 6to18 | 12.0 | 12 | 1.2 | 13.4 | 0.7 |
| | | v maple | 48 | 3to12 | 5.4 | 4 | 29.4 | 5.9 | 3.4 |

| Twp-Range | 8-2W | species | # WTs | Diam Range | av Diam | Median Diam | rel Freq% | QMD | rel DOM% |
|---|---|---|---|---|---|---|---|---|---|
| # all WTs | 267 | alder | 22 | 6to30 | 12.6 | 12 | 8.2 | 13.7 | 2.8 |
| all avDiam | 18.9 | ash | 14 | 6to40 | 15.9 | 13.5 | 5.2 | 18.3 | 3.2 |
| ttl QMD | 23.5 | redcedar | 26 | 6to50 | 26.9 | 24 | 9.7 | 29.7 | 15.6 |
| SrvyYr | 1856 | cherry | 5 | 3to7 | 4.4 | 3 | 1.9 | 4.7 | 0.1 |
| | | crabapple | 11 | 4to8 | 6.4 | 6 | 4.1 | 6.5 | 0.3 |
| | | cottonwood | 11 | 10to50 | 23.6 | 20 | 4.1 | 27.5 | 5.7 |
| | | dogwood | 4 | 6to10 | 8.0 | 8 | 1.5 | 8.1 | 0.2 |
| | | Douglas-fir | 115 | 5to80 | 24.8 | 20 | 43.1 | 28.7 | 64.3 |
| | | hemlock | 3 | 8to15 | 11.7 | 12 | 1.1 | 12.0 | 0.3 |
| | | bl maple | 22 | 3to36 | 14.7 | 12 | 8.2 | 17.9 | 4.8 |
| | | oak | 2 | 18to50 | 34.0 | 34 | 0.7 | 37.6 | 1.9 |
| | | cascara | 1 | 8 | 8.0 | 8 | 0.4 | 8.0 | 0.0 |
| | | hawthorn | 1 | 6 | 6.0 | 6 | 0.4 | 6.0 | 0.0 |
| | | v maple | 8 | 3to8 | 4.8 | 4.5 | 3.0 | 5.0 | 0.1 |
| | | willow | 22 | 3to18 | 6.0 | 5 | 8.2 | 6.9 | 0.7 |

| Twp-Range | 8-1W | species | # WTs | Diam Range | av Diam | Median Diam | rel Freq% | QMD | rel DOM% |
|---|---|---|---|---|---|---|---|---|---|
| # all WTs | 259 | alder | 8 | 6to16 | 10.0 | 9 | 3.1 | 10.4 | 0.8 |
| all avDiam | 15.7 | ash | 1 | 12 | 12.0 | 12 | 0.4 | 12.0 | 0.1 |
| ttl QMD | 20.1 | redcedar | 43 | 3to50 | 21.5 | 18 | 16.6 | 25.2 | 26.1 |
| SrvyYr | 1872 | cherry | 7 | 3to6 | 5.0 | 5 | 2.7 | 5.1 | 0.2 |
| | | dogwood | 1 | 10 | 10.0 | 10 | 0.4 | 10.0 | 0.1 |
| | | elder | 1 | 10 | 10.0 | 10 | 0.4 | 10.0 | 0.1 |
| | | Douglas-fir | 79 | 3to72 | 22.1 | 20 | 30.5 | 26.4 | 52.7 |
| | | true fir | 3 | 18to30 | 33.7 | 20 | 1.2 | 23.3 | 1.6 |
| | | hemlock | 60 | 3to36 | 13.3 | 12 | 23.2 | 15.4 | 13.6 |
| | | hazel | 3 | 3to5 | 4.0 | 4 | 1.2 | 4.1 | 0.0 |
| | | bl maple | 19 | 3to36 | 11.6 | 8 | 7.3 | 14.6 | 3.9 |
| | | v maple | 25 | 1to8 | 4.1 | 4 | 9.7 | 4.4 | 0.5 |
| | | willow | 9 | 3to10 | 6 | 6 | 3.5 | 6.4 | 0.3 |

| Twp-Range | 8-1E | species | # WTs | Diam Range | av Diam | Median Diam | rel Freq% | QMD | rel DOM% |
|---|---|---|---|---|---|---|---|---|---|
| # all WTs | 283 | alder | 5 | 12to20 | 14.4 | 14 | 1.8 | 14.7 | 0.9 |
| all avDiam | 15.3 | barberry | 4 | 4to7 | 5.8 | 6 | 1.4 | 5.9 | 0.1 |
| ttl QMD | 20.4 | redcedar | 25 | 5to72 | 19.4 | 12 | 8.8 | 25.3 | 13.6 |
| SrvyYr | 1872 | cherry | 6 | 3to8 | 5.2 | 4.5 | 2.1 | 5.5 | 0.2 |
| | | elder | 1 | 10 | 10.0 | 10 | 0.4 | 10.0 | 0.1 |
| | | Douglas-fir | 67 | 5to60 | 23.1 | 16 | 23.7 | 28.5 | 46.4 |
| | | true fir | 1 | 10 | 10.0 | 10 | 0.4 | 10.0 | 0.1 |
| | | hemlock | 114 | 4to72 | 15.8 | 14 | 40.3 | 19.1 | 35.3 |
| | | hazel | 1 | 3 | 3.0 | 3 | 0.4 | 3 | 0.0 |
| | | bl maple | 12 | 4to36 | 13.0 | 11 | 4.2 | 15.8 | 2.6 |
| | | v maple | 45 | 1to8 | 4.3 | 4 | 15.9 | 4.5 | 0.8 |
| | | willow | 2 | 5to8 | 6.5 | 6.5 | 0.7 | 6.67 | 0.1 |

| Twp-Range | 8-2E | species | # WTs | Diam Range | av Diam | Median Diam | rel Freq% | QMD | rel DOM% |
|---|---|---|---|---|---|---|---|---|---|
| # all WTs | 287 | alder | 6 | 10to14 | 12.3 | 13 | 2.1 | 12.5 | 0.2 |
| all avDiam | 31.0 | redcedar | 9 | 10to48 | 29.6 | 30 | 3.1 | 32.7 | 2.5 |
| ttl QMD | 36.4 | crabapple | 1 | 5 | 5.0 | 5 | 0.3 | 5.0 | 0.0 |
| SrvyYr | 1893 | Douglas-fir | 111 | 4to96 | 44.1 | 40 | 38.7 | 48.9 | 69.8 |
| | | hemlock | 145 | 3to48 | 23.0 | 22 | 50.5 | 25.0 | 23.8 |
| | | larch | 4 | 18to72 | 47.5 | 50 | 1.4 | 51.7 | 2.8 |
| | | bl maple | 3 | 20to30 | 24.7 | 24 | 1.0 | 25.0 | 0.5 |
| | | v maple | 8 | 3to6 | 4.0 | 3.5 | 2.8 | 4.2 | 0.0 |

| Twp-Range | 7-2W | species | # WTs | Diam Range | av Diam | Median Diam | rel Freq% | QMD | rel DOM% |
|---|---|---|---|---|---|---|---|---|---|
| # all WTs | 97 | alder | 3 | 8to10 | 9.3 | 10 | 3.1 | 9.4 | 0.9 |
| all avDiam | 14.0 | ash | 32 | 2to50 | 13.4 | 10 | 33.0 | 17.0 | 29.8 |
| ttl QMD | 17.9 | redcedar | 7 | 10to30 | 16.1 | 15 | 7.2 | 17.3 | 6.8 |
| SrvyYr | 1856 | cottonwood | 14 | 7to50 | 28.1 | 26 | 14.4 | 31.7 | 45.3 |
| | | Douglas-fir | 6 | 5to24 | 15.5 | 16 | 6.2 | 17.3 | 5.8 |
| | | hawthorn | 2 | 6to8 | 7.0 | 7 | 2.1 | 7.1 | 0.3 |
| | | hemlock | 1 | 8 | 8.0 | 8 | 1.0 | 8.0 | 0.2 |
| | | bl maple | 2 | 3to8 | 5.5 | 5.5 | 2.1 | 6.0 | 0.2 |
| | | oak | 2 | 20to30 | 25.0 | 25 | 2.1 | 25.5 | 4.2 |
| | | willow | 28 | 3to15 | 7.9 | 8 | 28.9 | 8.5 | 6.5 |

| Twp-Range | 7-1W | species | # WTs | Diam Range | av Diam | Median Diam | rel Freq% | QMD | rel DOM% |
|---|---|---|---|---|---|---|---|---|---|
| # all WTs | 285 | alder | 18 | 4to12 | 7.6 | 6.5 | 6.3 | 7.9 | 0.5 |
| all avDiam | 21.1 | ash | 2 | 8to18 | 13.0 | 13 | 0.7 | 14.0 | 0.2 |
| ttl QMD | 27.5 | redcedar | 59 | 4to80 | 25.3 | 20 | 20.7 | 30.1 | 24.8 |
| SrvyYr | 1856&72 | cherry | 17 | 2to10 | 4.0 | 3 | 6.0 | 4.4 | 0.2 |
| | | Douglas-fir | 114 | 3to80 | 31.9 | 30 | 40.0 | 36.2 | 69.3 |
| | | true fir | 1 | 18 | 18.0 | 18 | 0.4 | 18.0 | 0.2 |
| | | hemlock | 12 | 8to50 | 20.5 | 17 | 4.2 | 23.7 | 3.1 |
| | | hawthorn | 2 | 6to8 | 7.0 | 7 | 0.7 | 7.1 | 0.0 |
| | | bl maple | 26 | 1to18 | 7.9 | 8 | 9.1 | 9.3 | 1.0 |
| | | oak | 3 | 8to20 | 12.7 | 10 | 1.1 | 13.7 | 0.3 |
| | | v maple | 16 | 1to6 | 3.3 | 3 | 5.6 | 3.6 | 0.1 |
| | | willow | 15 | 2to20 | 5.2 | 3 | 5.3 | 6.8 | 0.3 |

| Twp-Range | 7-1E | species | # WTs | Diam Range | av Diam | Median Diam | rel Freq% | QMD | rel DOM% |
|---|---|---|---|---|---|---|---|---|---|
| # all WTs | 236 | alder | 5 | 7to14 | 10.2 | 9 | 2.1 | 10.5 | 0.3 |
| all avDiam | 21.6 | barberry | 2 | 3to6 | 4.5 | 4.5 | 0.9 | 4.7 | 0.0 |
| ttl QMD | 29.8 | redcedar | 2 | 16to70 | 43.0 | 43 | 0.9 | 50.8 | 2.5 |
| SrvyYr | 1872 | cherry | 11 | 1to7 | 3.5 | 3 | 4.7 | 3.8 | 0.1 |
| | | elder | 1 | 4 | 4.0 | 4 | 0.4 | 4.0 | 0.0 |
| | | Douglas-for | 161 | 2to80 | 28.6 | 30 | 68.5 | 35.3 | 96.2 |
| | | hemlock | 11 | 3to20 | 10.7 | 10 | 4.7 | 12.1 | 0.8 |
| | | hazel | 2 | 2to3 | 2.5 | 2.5 | 0.9 | 2.6 | 0.0 |
| | | bl maple | 10 | 1to26 | 6.3 | 4 | 4.3 | 9.3 | 0.4 |
| | | hawthorn | 2 | 3 | 3.0 | 3 | 0.9 | 3.0 | 0.0 |
| | | v maple | 28 | 1to6 | 3.6 | 3 | 11.9 | 3.8 | 0.2 |
| | | willow | 1 | 2 | 2.0 | 2 | 0.4 | 2.0 | 0.0 |

| Twp-Range | 7-2E | species | # WTs | Diam Range | av Diam | Median Diam | rel Freq% | QMD | rel DOM% |
|---|---|---|---|---|---|---|---|---|---|
| # all WTs | 286 | alder | 24 | 10to24 | 12.0 | 12 | 8.4 | 12.3 | 1.9 |
| all avDiam | 20.8 | barberry | 3 | 4to6 | 4.7 | 4 | 1.0 | 4.8 | 0.0 |
| ttl QMD | 25.7 | redcedar | 10 | 10to48 | 24.2 | 12 | 3.5 | 29.8 | 4.7 |
| SrvyYr | 1894 | Douglas-fir | 156 | 4to72 | 24.4 | 14 | 54.5 | 30.2 | 75.5 |
| | | hemlock | 83 | 4to48 | 17.7 | 16 | 29.0 | 19.4 | 16.6 |
| | | bl maple | 6 | 12to24 | 19.3 | 20 | 2.1 | 19.7 | 1.2 |
| | | v maple | 4 | 3to6 | 5.3 | 6 | 1.4 | 5.4 | 0.1 |
